# LRRK2 Kinase Activity Regulates Parkinson’s Disease-Relevant Lipids at the Lysosome

**DOI:** 10.1101/2022.12.19.521070

**Authors:** Michael T. Maloney, Xiang Wang, Rajarshi Ghosh, Shan V. Andrews, Romeo Maciuca, Shababa T. Masoud, Richard M. Caprioli, John Chen, Chi-Lu Chiu, Sonnet S. Davis, Audrey Cheuk-Nga Ho, Hoang N. Nguyen, Nicholas E. Propson, Michelle L. Reyzer, Oliver B. Davis, Matthew C. Deen, Sha Zhu, Gilbert Di Paolo, David J. Vocadlo, Anthony A. Estrada, Javier de Vicente, Joseph W. Lewcock, Annie Arguello, Jung H. Suh, Sarah Huntwork-Rodriguez, Anastasia G. Henry

## Abstract

Lysosomal dysfunction is a hallmark of Parkinson’s disease (PD), and variants in several PD-associated genes, including *LRRK2*, perturb lysosomal homeostasis. Based on this, LRRK2 kinase inhibition is being explored as a therapeutic approach for the treatment of PD. LRRK2 inhibitors reduce levels of BMP, an endolysosomal lipid involved in glycosphingolipid (GSL) catabolism, in urine from preclinical models and clinical subjects, however, the mechanisms by which LRRK2 regulates BMP and the functional significance of this change to disease are undefined. We establish that LRRK2 regulates secretion of BMP- and GSL-containing vesicles from kidney into urine and modulates BMP and GSL levels in the brain. BMP accumulates within lysosomes as a secondary response to LRRK2’s effects on the activity of glucocerebrosidase (GCase), a PD-linked enzyme involved in GSL catabolism. Alterations in BMP and GCase substrate turnover are observed in CSF from LRRK2-PD patients, highlighting the relevance of LRRK2-dependent lysosomal dysfunction in disease.

## Introduction

Lysosomal dysfunction has emerged as a principal contributor to the susceptibility and pathogenesis of several neurodegenerative diseases including PD. Whole exome sequencing and genome-wide association studies focused on identifying genetic risk factors for PD have converged on genes associated with PD risk that function in the autophagic and endolysosomal pathways such as *LRRK2*, *GBA1*, and *TMEM175*(1-5). The link between lysosomal dysfunction and PD is bolstered by an increased burden of variants in several genes that cause rare monogenic lysosomal storage disorders (LSDs) in PD patients, suggesting common mechanisms of lysosomal dysfunction drive both severe LSDs and PD(6-8). Defects in lysosomal homeostasis have also been observed in PD patient postmortem brain samples or biofluids, including depletion of the lysosomal compartment, glycosphingolipid (GSL) storage, and accumulation of lysosomal proteins (9-13). These data suggest that approaches aimed at correcting lysosomal dysfunction may provide new therapeutic avenues for the treatment of PD, for which there still remains no disease-modifying therapy.

LRRK2 kinase inhibition is currently being explored in late-stage clinical studies as one such approach to improve lysosomal function in PD. Variants in *LRRK2* are associated with increased risk for Parkinson’s and Crohn’s disease, and a putative protective variant has been identified (N551K R1398H) linked to reduced risk for disease(14-16). LRRK2 variants are proposed to modify PD risk by regulating the enzyme’s kinase activity, as most pathogenic variants lead to increased kinase activity and the protective variant is associated with reduced activity(17-19). LRRK2 phosphorylates a subset of Rab GTPases, known master regulators of the secretory and endocytic pathways, and PD-risk linked LRRK2 variants increase phosphorylation of these Rab substrates, including Rab10(20, 21). This phosphorylation is thought to impair Rab function by perturbing interactions with downstream effectors, resulting in defects in various aspects of intracellular trafficking, including autophagy and endolysosomal function. Consistent with this hypothesis, LRRK2 has been shown to act as both a sensor and trigger of lysosomal dysfunction as the kinase localizes to endolysosomal membranes upon damage and can modulate autophagic flux and lysosomal proteolysis(22-25). The detailed mechanisms by which LRRK2 regulates lysosomal function and the relevance of this dysfunction to PD, however, are still not clear.

We reasoned that insight into the mechanism by which LRRK2 modulates lysosomal homeostasis and the significance of such regulation in disease may come from the connection between LRRK2 kinase activity and levels of BMP (bis (monoacylglycerol)phosphate), an anionic phospholipid concentrated on intraluminal vesicles (ILVs) in late endosomes and lysosomes(26). BMP promotes the hydrolysis of GSLs by serving as an anionic lipid anchor and stimulator of lysosomal lipase activities, including the PD-linked enzyme glucocerebrosidase (GCase) encoded by *GBA1*(27, 28). LRRK2 kinase inhibition leads to reduced urine BMP levels in preclinical models and in human subjects(29, 30), and human subjects that carry the LRRK2 G2019S variant associated with increased kinase activity have higher levels of BMP in urine compared to controls(31). These observations collectively suggest LRRK2 kinase activity may be linked to lysosomal function through its action on BMP and GSL homeostasis. Currently, however, existing data only demonstrate that LRRK2 regulates the extracellular concentration of BMP, at least in urine, and it is difficult to interpret what such changes mean with respect to LRRK2’s impact on endolysosomal BMP levels, GSL catabolism, and lysosomal function more broadly. Moreover, whether LRRK2 similarly regulates BMP within brain is unknown.

Accordingly, a deeper understanding of the mechanisms by which LRRK2 regulates BMP and its functional consequences is needed to understand the potential of LRRK2 kinase inhibition to improve lysosomal dysfunction in PD as well as to determine the utility of BMP as a lysosomal biomarker of LRRK2 activity in the clinic.

Here, we define the mechanisms underlying LRRK2-dependent regulation of BMP levels and show that LRRK2-dependent changes in BMP correlate with GSL dysregulation and lysosomal dysfunction in preclinical models and PD patients. This regulation of lysosomal function by LRRK2 occurs in both kidney and brain and can be observed in PD patient CSF. We uncover the mechanisms by which LRRK2 regulates intracellular BMP levels, demonstrating that LRRK2 can regulate the secretion of BMP and promote BMP accumulation as a secondary consequence of LRRK2-mediated effects on GCase activity. Increased LRRK2 kinase activity correlates with defects in lysosomal proteolysis, and LRRK2 kinase inhibition reduces BMP and GCase substrate accumulation in lysosomes and ultimately restores lysosomal function. Together, these results support the use of BMP and GCase substrates as biomarkers of LRRK2-dependent regulation of lysosomal function in the clinic and highlight the therapeutic potential of LRRK2 inhibition to correct lysosomal dysfunction observed in PD.

## Results

### Variants in LRRK2 that impact kinase activity regulate BMP levels in urine from human subjects

Previous analyses in human subjects show that LRRK2 kinase activity regulates BMP levels in urine. Carriers of the kinase-activating LRRK2 G2019S variant have elevated urine BMP levels, and LRRK2 kinase inhibition resulted in reduced urine BMP levels in clinical studies(30, 31). To further investigate the relationship between risk and protective variants in LRRK2 that modulate kinase activity and BMP levels in humans, we used publicly available urine BMP data and whole genome sequencing (WGS) data in the Parkinson’s Progression Markers Initiative (PPMI). After restricting PPMI samples with available WGS data to those with available urine BMP data and those of predicted European ancestry (see Materials and Methods), 1,069 samples (derived from subjects with and without PD) were available for data analysis. This dataset was enriched for carriers of the PD risk variants LRRK2 G2019S (N = 289; 27%) and GBA N370S/N409S (N = 238; 22%). In models adjusting for sex, age, disease status and principal components derived from the WGS data, urine levels of total 22:6/22:6 BMP species were significantly increased in carriers of the PD risk variant G2019S (p = 5.78E-82; Fig. 1A and Supplementary Table S1), consistent with previous work(31, 32). To delve more deeply into the relationship between LRRK2 kinase activity and BMP, we next assessed whether the LRRK2 N551K protective variant impacted BMP levels in urine. Our recent work showed that the LRRK2 protective variant N551K R1398H is associated with reduced LRRK2 kinase activity, as reflected by levels of the LRRK2 kinase substrate phospho-Rab10 (pRab10) in both cellular models and human subjects(19). We observed a significant decrease in total 22:6/22:6 urine BMP levels in carriers of the PD protective variant N551K (p = 7.33E-4; Fig. 1A and Supplementary Table S1). These relationships were maintained (consistent direction of effect and p < 0.05) in sensitivity analyses that removed the adjustment by disease status or additionally adjusted for GBA N370S/N409S status, N551K status (for the G2019S effect) or G2019S status (for the N551K effect). This analysis provides evidence that genetic variants that either increase or decrease LRRK2 kinase activity have the concurrent effect on extracellular levels of BMP in urine. Additionally, the reduction in urine BMP observed in carriers of N551K LRRK2 protective allele further supports therapeutic strategies focused on reduction of LRRK2 kinase inhibition.

**Figure 1:**
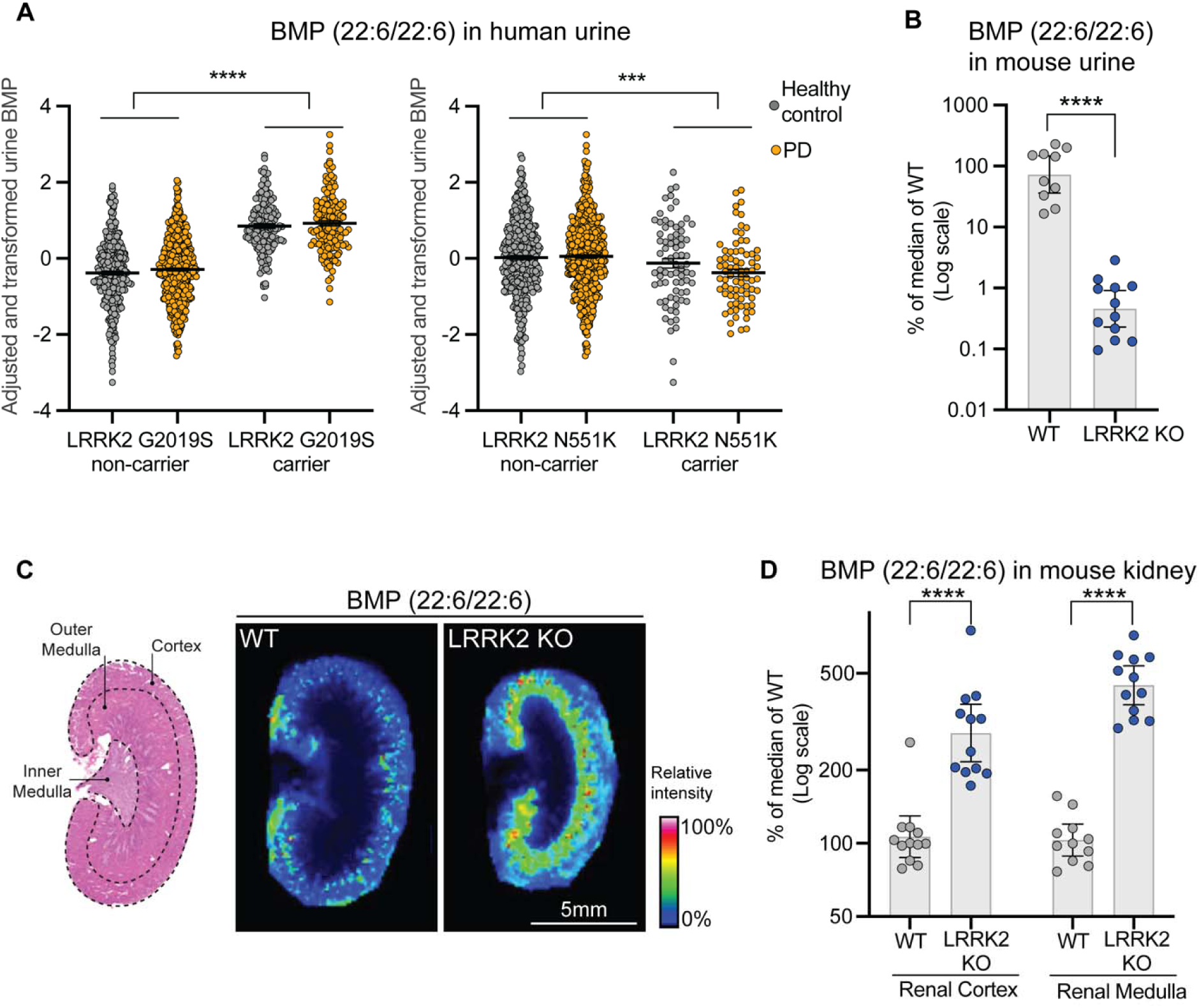
LRRK2 activity regulates peripheral BMP levels in human subjects and in preclinical models. A) Association of urine BMP (22:6/22:6) levels and carrier status at the PD-risk LRRK2 G2019S variant and the PD-protective LRRK2 N551K variant in PPMI data. Urine BMP levels were normalized to creatinine levels, log transformed, and fit in a linear model against sex, age, disease status, and the first five principal components derived from whole genome sequencing data. Inverse normal transformed residuals from this linear model are plotted on the y-axis and used in association testing with LRRK2 variant status. Data are shown as mean ± SEM. B) Urine BMP (22:6/22:6) levels from LRRK2 KO mice (n=12) showed a significant reduction (down to 1%) compared to wild-type (WT) littermates (n=10). The relative abundance of urine BMP (22:6/22:6) levels were normalized to creatinine, measured by liquid chromatography tandem mass spectrometry (LC-MS/MS), and then presented as a percent of median values of WT group. C) Matrix-assisted laser desorption/ionization mass spectrometry images were acquired from longitudinal kidney sections of WT and LRRK2 KO mice. Left panel: Representative hematoxylin and eosin photomicrograph of the kidney showing demarcated regions (cortex, outer medulla, and inner medulla). Right panel: Representative mass spectrometry image showing the distribution of the signal at a mass/charge ratio (m/z) of 865.502, corresponding to BMP (22:6/22:6). Images depict the relative intensity of the signal from 0 to 100% with intensity normalized using total ion current. D) The relative abundance of BMP (22:6/22:6) in the renal cortex and medulla from LRRK2 KO mice and WT littermates was measured by LC-MS/MS and normalized as a percent of median values of the WT group; n=12 animals for each group. B, D) Data are shown as geometric mean ratio (%) and 95% confidence intervals (CI), with statistical significance assessed based on Benjamini-Hochberg (BH)-adjusted p-values. ***P ≤ 0.001, ****P ≤ 0.0001.

**Table 1:**
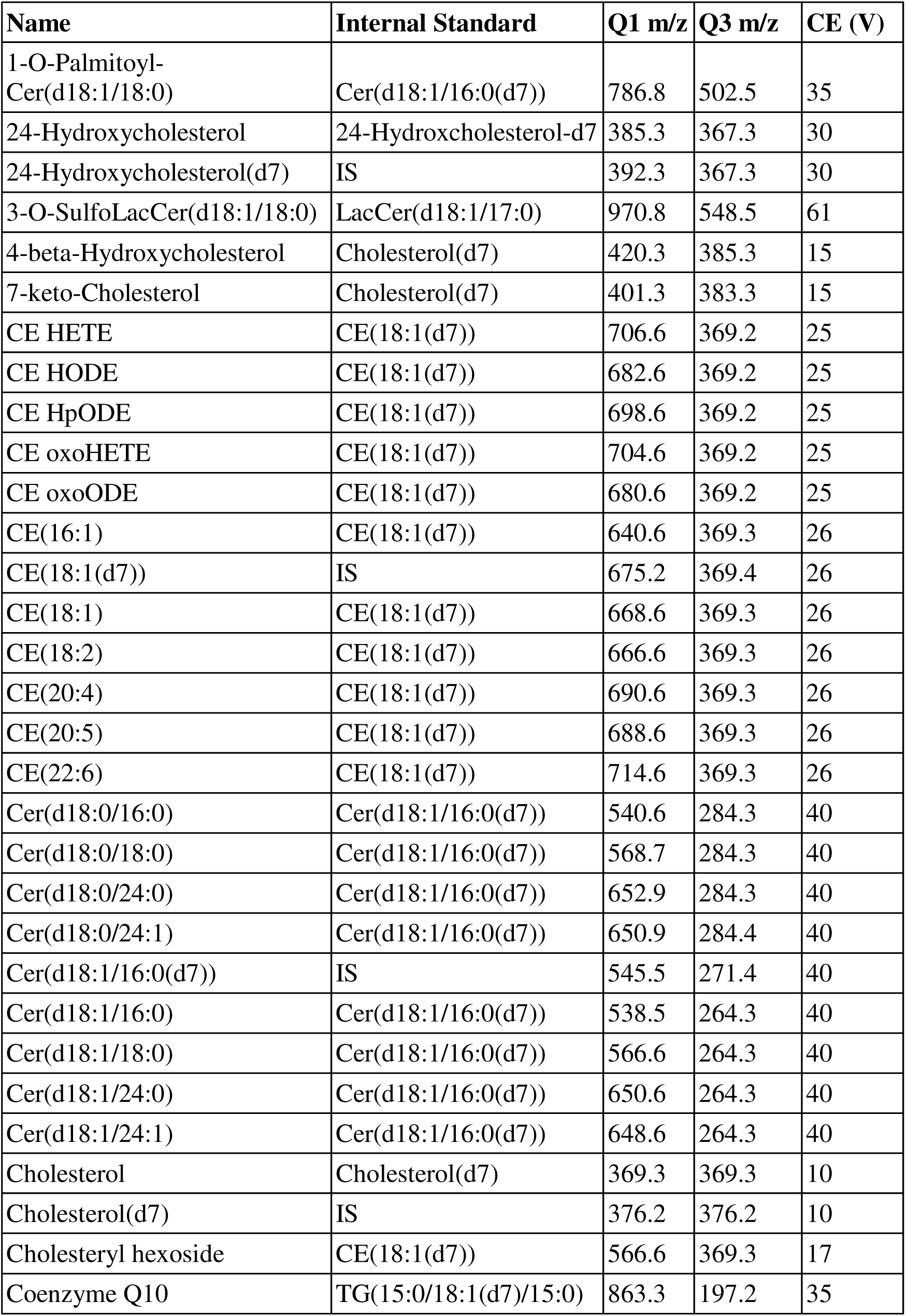
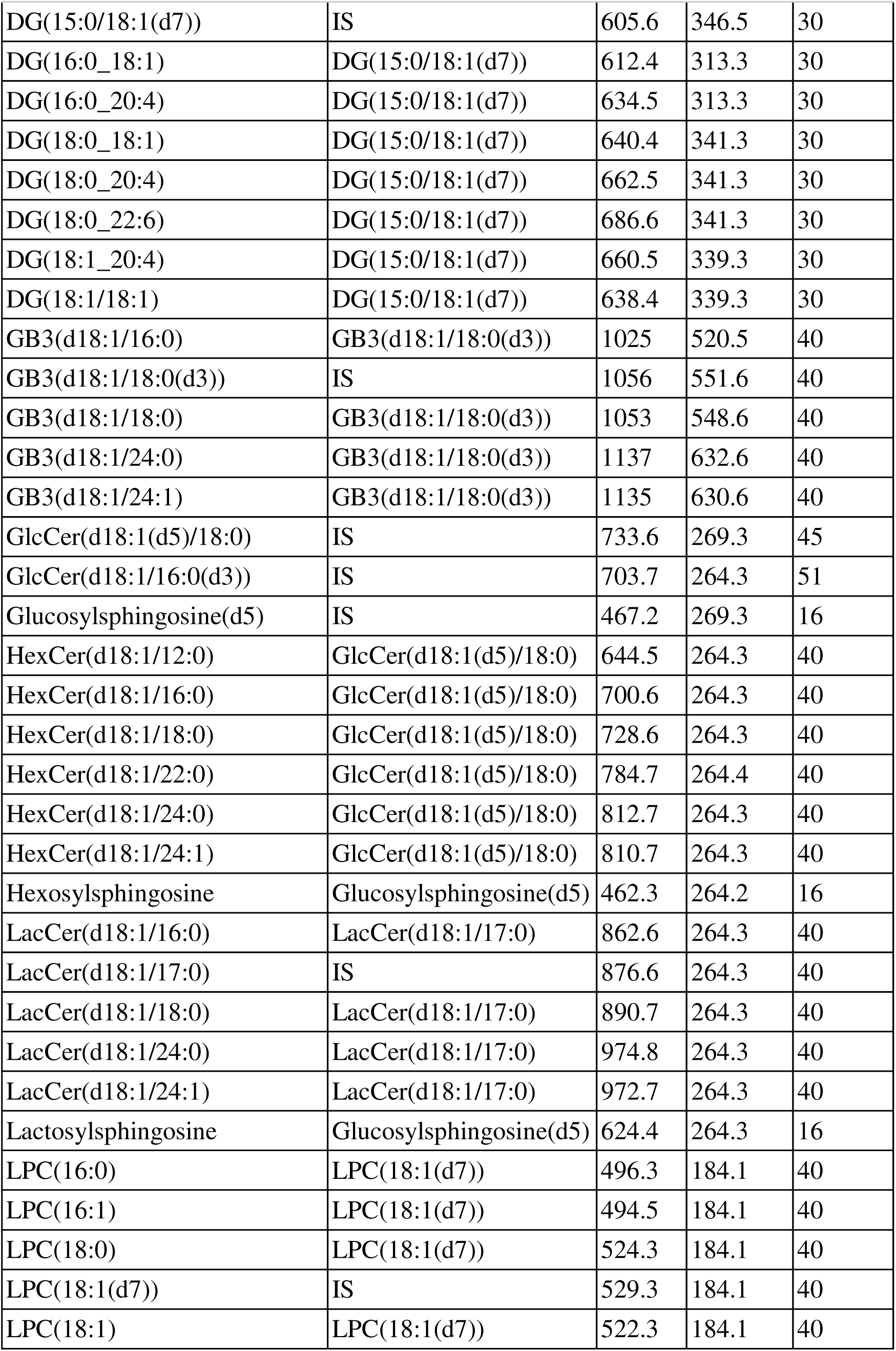
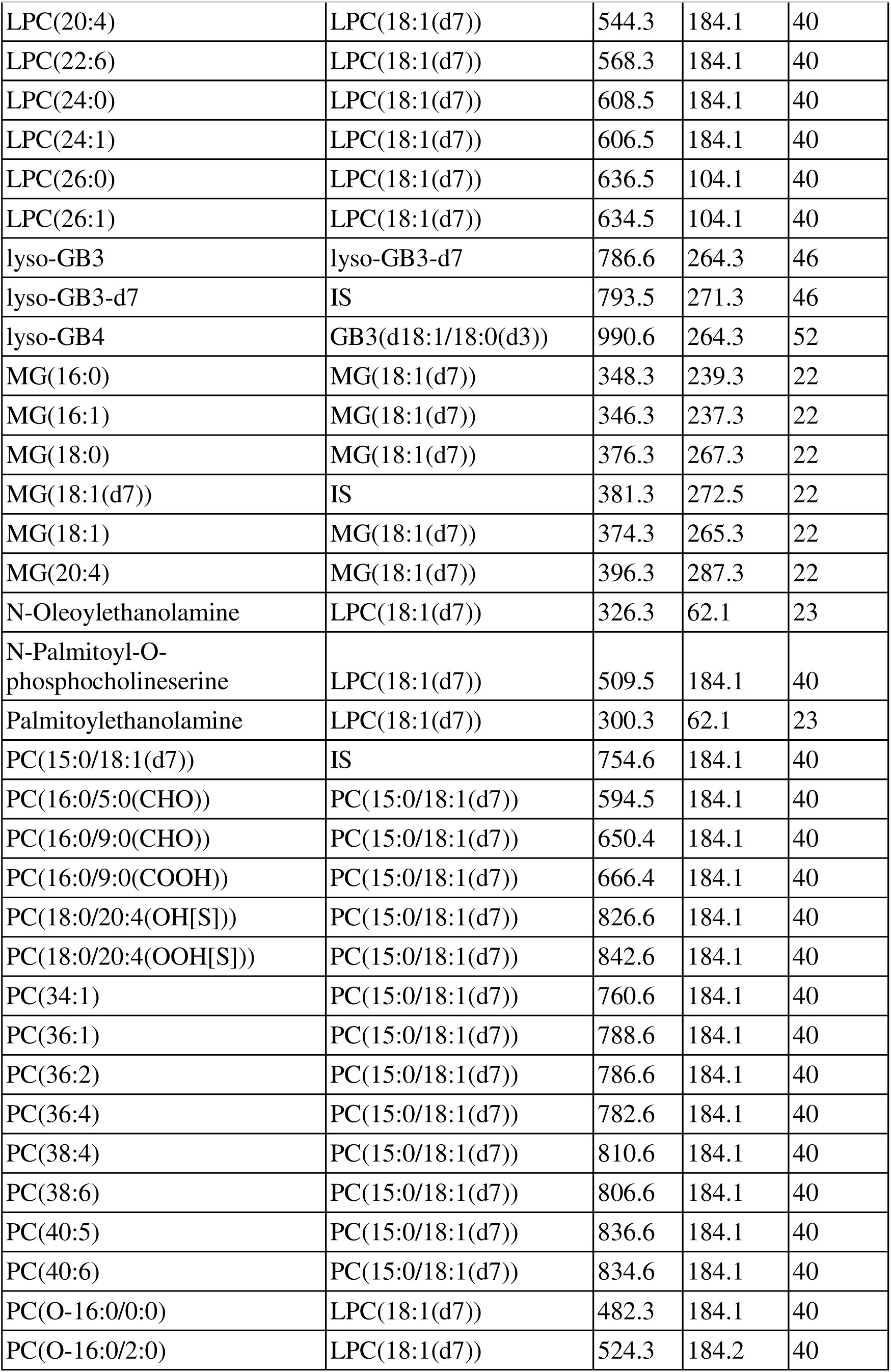
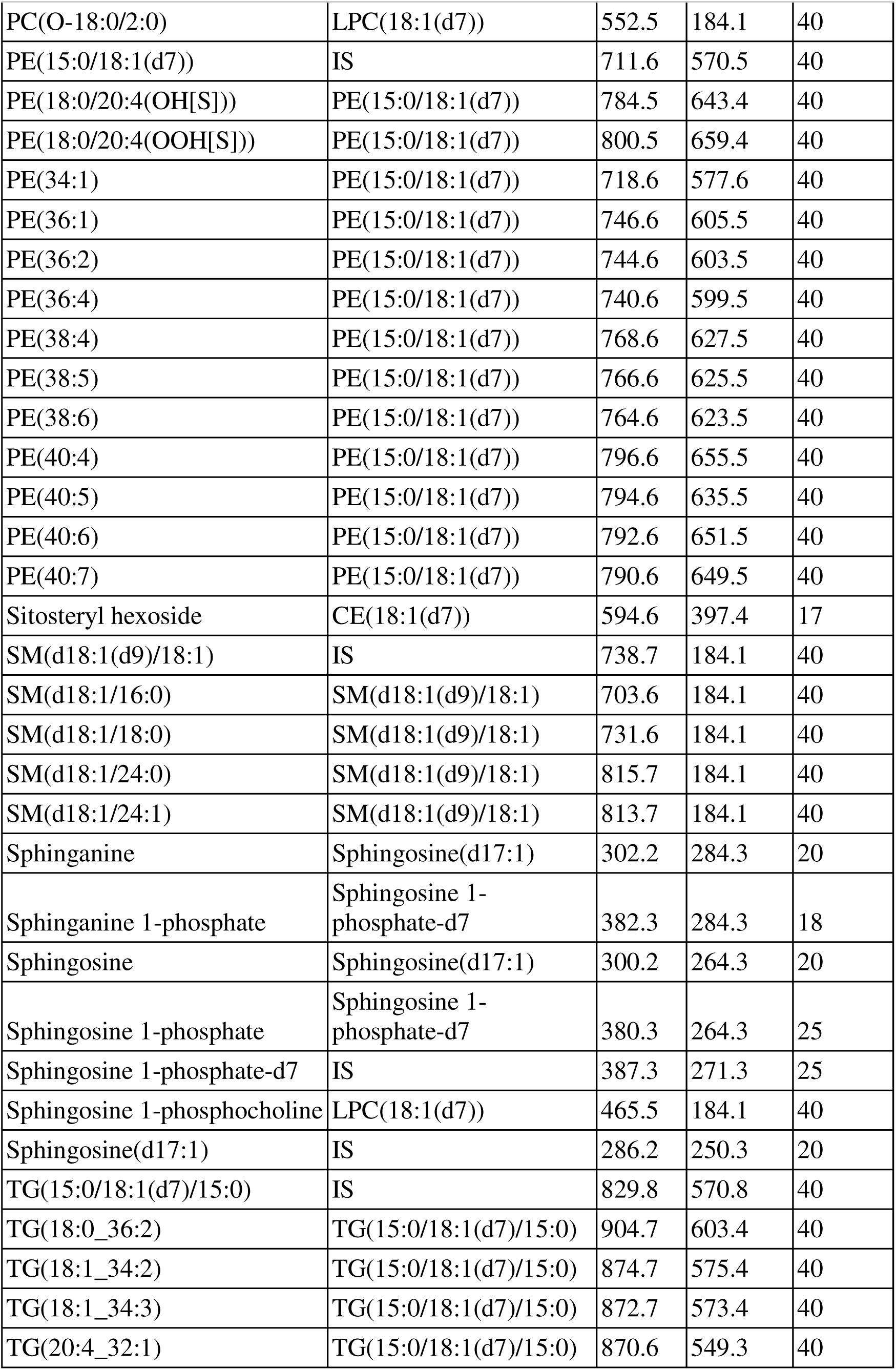
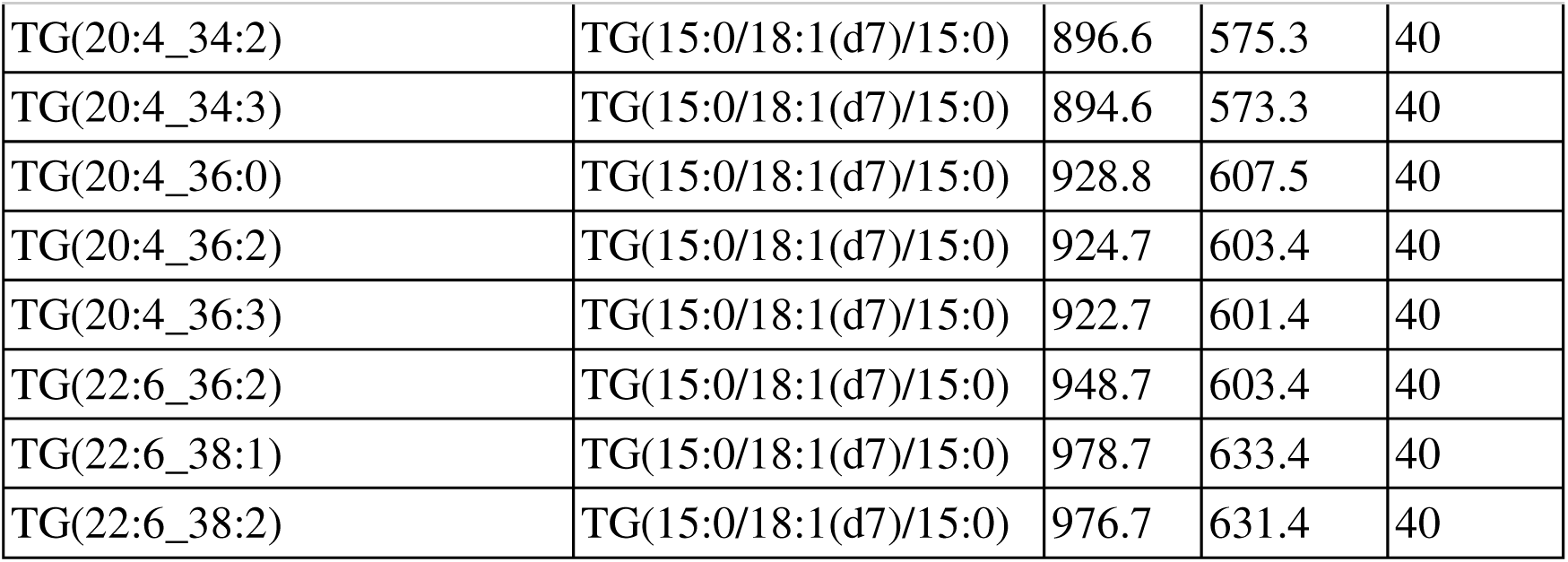
Lipidomics in positive mode parameters

| Name | Internal Standard | Q1 m/z | Q3 m/z | CE (V) |
| --- | --- | --- | --- | --- |
| 1-O-Palmitoyl-Cer(d18:1/18:0) | Cer(d18:1/16:0(d7)) | 786.8 | 502.5 | 35 |
| 24-Hydroxycholesterol | 24-Hydroxcholesterol-d7 | 385.3 | 367.3 | 30 |
| 24-Hydroxycholesterol(d7) | IS | 392.3 | 367.3 | 30 |
| 3-O-SulfoLacCer(d18:1/18:0) | LacCer(d18:1/17:0) | 970.8 | 548.5 | 61 |
| 4-beta-Hydroxycholesterol | Cholesterol(d7) | 420.3 | 385.3 | 15 |
| 7-keto-Cholesterol | Cholesterol(d7) | 401.3 | 383.3 | 15 |
| CE HETE | CE(18:1(d7)) | 706.6 | 369.2 | 25 |
| CE HODE | CE(18:1(d7)) | 682.6 | 369.2 | 25 |
| CE HpODE | CE(18:1(d7)) | 698.6 | 369.2 | 25 |
| CE oxoHETE | CE(18:1(d7)) | 704.6 | 369.2 | 25 |
| CE oxoODE | CE(18:1(d7)) | 680.6 | 369.2 | 25 |
| CE(16:1) | CE(18:1(d7)) | 640.6 | 369.3 | 26 |
| CE(18:1(d7)) | IS | 675.2 | 369.4 | 26 |
| CE(18:1) | CE(18:1(d7)) | 668.6 | 369.3 | 26 |
| CE(18:2) | CE(18:1(d7)) | 666.6 | 369.3 | 26 |
| CE(20:4) | CE(18:1(d7)) | 690.6 | 369.3 | 26 |
| CE(20:5) | CE(18:1(d7)) | 688.6 | 369.3 | 26 |
| CE(22:6) | CE(18:1(d7)) | 714.6 | 369.3 | 26 |
| Cer(d18:0/16:0) | Cer(d18:1/16:0(d7)) | 540.6 | 284.3 | 40 |
| Cer(d18:0/18:0) | Cer(d18:1/16:0(d7)) | 568.7 | 284.3 | 40 |
| Cer(d18:0/24:0) | Cer(d18:1/16:0(d7)) | 652.9 | 284.3 | 40 |
| Cer(d18:0/24:1) | Cer(d18:1/16:0(d7)) | 650.9 | 284.4 | 40 |
| Cer(d18:1/16:0(d7)) | IS | 545.5 | 271.4 | 40 |
| Cer(d18:1/16:0) | Cer(d18:1/16:0(d7)) | 538.5 | 264.3 | 40 |
| Cer(d18:1/18:0) | Cer(d18:1/16:0(d7)) | 566.6 | 264.3 | 40 |
| Cer(d18:1/24:0) | Cer(d18:1/16:0(d7)) | 650.6 | 264.3 | 40 |
| Cer(d18:1/24:1) | Cer(d18:1/16:0(d7)) | 648.6 | 264.3 | 40 |
| Cholesterol | Cholesterol(d7) | 369.3 | 369.3 | 10 |
| Cholesterol(d7) | IS | 376.2 | 376.2 | 10 |
| Cholesteryl hexoside | CE(18:1(d7)) | 566.6 | 369.3 | 17 |
| Coenzyme Q10 | TG(15:0/18:1(d7)/15:0) | 863.3 | 197.2 | 35 |
| DG(15:0/18:1(d7)) | IS | 605.6 | 346.5 | 30 |
| DG(16:0_18:1) | DG(15:0/18:1(d7)) | 612.4 | 313.3 | 30 |
| DG(16:0_20:4) | DG(15:0/18:1(d7)) | 634.5 | 313.3 | 30 |
| DG(18:0_18:1) | DG(15:0/18:1(d7)) | 640.4 | 341.3 | 30 |
| DG(18:0_20:4) | DG(15:0/18:1(d7)) | 662.5 | 341.3 | 30 |
| DG(18:0_22:6) | DG(15:0/18:1(d7)) | 686.6 | 341.3 | 30 |
| DG(18:1_20:4) | DG(15:0/18:1(d7)) | 660.5 | 339.3 | 30 |
| DG(18:1/18:1) | DG(15:0/18:1(d7)) | 638.4 | 339.3 | 30 |
| GB3(d18:1/16:0) | GB3(d18:1/18:0(d3)) | 1025 | 520.5 | 40 |
| GB3(d18:1/18:0(d3)) | IS | 1056 | 551.6 | 40 |
| GB3(d18:1/18:0) | GB3(d18:1/18:0(d3)) | 1053 | 548.6 | 40 |
| GB3(d18:1/24:0) | GB3(d18:1/18:0(d3)) | 1137 | 632.6 | 40 |
| GB3(d18:1/24:1) | GB3(d18:1/18:0(d3)) | 1135 | 630.6 | 40 |
| GlcCer(d18:1(d5)/18:0) | IS | 733.6 | 269.3 | 45 |
| GlcCer(d18:1/16:0(d3)) | IS | 703.7 | 264.3 | 51 |
| Glucosylsphingosine(d5) | IS | 467.2 | 269.3 | 16 |
| HexCer(d18:1/12:0) | GlcCer(d18:1(d5)/18:0) | 644.5 | 264.3 | 40 |
| HexCer(d18:1/16:0) | GlcCer(d18:1(d5)/18:0) | 700.6 | 264.3 | 40 |
| HexCer(d18:1/18:0) | GlcCer(d18:1(d5)/18:0) | 728.6 | 264.3 | 40 |
| HexCer(d18:1/22:0) | GlcCer(d18:1(d5)/18:0) | 784.7 | 264.4 | 40 |
| HexCer(d18:1/24:0) | GlcCer(d18:1(d5)/18:0) | 812.7 | 264.3 | 40 |
| HexCer(d18:1/24:1) | GlcCer(d18:1(d5)/18:0) | 810.7 | 264.3 | 40 |
| Hexosylsphingosine | Glucosylsphingosine(d5) | 462.3 | 264.2 | 16 |
| LacCer(d18:1/16:0) | LacCer(d18:1/17:0) | 862.6 | 264.3 | 40 |
| LacCer(d18:1/17:0) | IS | 876.6 | 264.3 | 40 |
| LacCer(d18:1/18:0) | LacCer(d18:1/17:0) | 890.7 | 264.3 | 40 |
| LacCer(d18:1/24:0) | LacCer(d18:1/17:0) | 974.8 | 264.3 | 40 |
| LacCer(d18:1/24:1) | LacCer(d18:1/17:0) | 972.7 | 264.3 | 40 |
| Lactosylsphingosine | Glucosylsphingosine(d5) | 624.4 | 264.3 | 16 |
| LPC(16:0) | LPC(18:1(d7)) | 496.3 | 184.1 | 40 |
| LPC(16:1) | LPC(18:1(d7)) | 494.5 | 184.1 | 40 |
| LPC(18:0) | LPC(18:1(d7)) | 524.3 | 184.1 | 40 |
| LPC(18:1(d7)) | IS | 529.3 | 184.1 | 40 |
| LPC(18:1) | LPC(18:1(d7)) | 522.3 | 184.1 | 40 |
| LPC(20:4) | LPC(18:1(d7)) | 544.3 | 184.1 | 40 |
| LPC(22:6) | LPC(18:1(d7)) | 568.3 | 184.1 | 40 |
| LPC(24:0) | LPC(18:1(d7)) | 608.5 | 184.1 | 40 |
| LPC(24:1) | LPC(18:1(d7)) | 606.5 | 184.1 | 40 |
| LPC(26:0) | LPC(18:1(d7)) | 636.5 | 104.1 | 40 |
| LPC(26:1) | LPC(18:1(d7)) | 634.5 | 104.1 | 40 |
| lyso-GB3 | lyso-GB3-d7 | 786.6 | 264.3 | 46 |
| lyso-GB3-d7 | IS | 793.5 | 271.3 | 46 |
| lyso-GB4 | GB3(d18:1/18:0(d3)) | 990.6 | 264.3 | 52 |
| MG(16:0) | MG(18:1(d7)) | 348.3 | 239.3 | 22 |
| MG(16:1) | MG(18:1(d7)) | 346.3 | 237.3 | 22 |
| MG(18:0) | MG(18:1(d7)) | 376.3 | 267.3 | 22 |
| MG(18:1(d7)) | IS | 381.3 | 272.5 | 22 |
| MG(18:1) | MG(18:1(d7)) | 374.3 | 265.3 | 22 |
| MG(20:4) | MG(18:1(d7)) | 396.3 | 287.3 | 22 |
| N-Oleoylethanolamine | LPC(18:1(d7)) | 326.3 | 62.1 | 23 |
| N-Palmitoyl-O-phosphocholineserine | LPC(18:1(d7)) | 509.5 | 184.1 | 40 |
| Palmitoylethanolamine | LPC(18:1(d7)) | 300.3 | 62.1 | 23 |
| PC(15:0/18:1(d7)) | IS | 754.6 | 184.1 | 40 |
| PC(16:0/5:0(CHO)) | PC(15:0/18:1(d7)) | 594.5 | 184.1 | 40 |
| PC(16:0/9:0(CHO)) | PC(15:0/18:1(d7)) | 650.4 | 184.1 | 40 |
| PC(16:0/9:0(COOH)) | PC(15:0/18:1(d7)) | 666.4 | 184.1 | 40 |
| PC(18:0/20:4(OH[S])) | PC(15:0/18:1(d7)) | 826.6 | 184.1 | 40 |
| PC(18:0/20:4(OOH[S])) | PC(15:0/18:1(d7)) | 842.6 | 184.1 | 40 |
| PC(34:1) | PC(15:0/18:1(d7)) | 760.6 | 184.1 | 40 |
| PC(36:1) | PC(15:0/18:1(d7)) | 788.6 | 184.1 | 40 |
| PC(36:2) | PC(15:0/18:1(d7)) | 786.6 | 184.1 | 40 |
| PC(36:4) | PC(15:0/18:1(d7)) | 782.6 | 184.1 | 40 |
| PC(38:4) | PC(15:0/18:1(d7)) | 810.6 | 184.1 | 40 |
| PC(38:6) | PC(15:0/18:1(d7)) | 806.6 | 184.1 | 40 |
| PC(40:5) | PC(15:0/18:1(d7)) | 836.6 | 184.1 | 40 |
| PC(40:6) | PC(15:0/18:1(d7)) | 834.6 | 184.1 | 40 |
| PC(O-16:0/0:0) | LPC(18:1(d7)) | 482.3 | 184.1 | 40 |
| PC(O-16:0/2:0) | LPC(18:1(d7)) | 524.3 | 184.2 | 40 |
| PC(O-18:0/2:0) | LPC(18:1(d7)) | 552.5 | 184.1 | 40 |
| PE(15:0/18:1(d7)) | IS | 711.6 | 570.5 | 40 |
| PE(18:0/20:4(OH[S])) | PE(15:0/18:1(d7)) | 784.5 | 643.4 | 40 |
| PE(18:0/20:4(OOH[S])) | PE(15:0/18:1(d7)) | 800.5 | 659.4 | 40 |
| PE(34:1) | PE(15:0/18:1(d7)) | 718.6 | 577.6 | 40 |
| PE(36:1) | PE(15:0/18:1(d7)) | 746.6 | 605.5 | 40 |
| PE(36:2) | PE(15:0/18:1(d7)) | 744.6 | 603.5 | 40 |
| PE(36:4) | PE(15:0/18:1(d7)) | 740.6 | 599.5 | 40 |
| PE(38:4) | PE(15:0/18:1(d7)) | 768.6 | 627.5 | 40 |
| PE(38:5) | PE(15:0/18:1(d7)) | 766.6 | 625.5 | 40 |
| PE(38:6) | PE(15:0/18:1(d7)) | 764.6 | 623.5 | 40 |
| PE(40:4) | PE(15:0/18:1(d7)) | 796.6 | 655.5 | 40 |
| PE(40:5) | PE(15:0/18:1(d7)) | 794.6 | 635.5 | 40 |
| PE(40:6) | PE(15:0/18:1(d7)) | 792.6 | 651.5 | 40 |
| PE(40:7) | PE(15:0/18:1(d7)) | 790.6 | 649.5 | 40 |
| Sitosteryl hexoside | CE(18:1(d7)) | 594.6 | 397.4 | 17 |
| SM(d18:1(d9)/18:1) | IS | 738.7 | 184.1 | 40 |
| SM(d18:1/16:0) | SM(d18:1(d9)/18:1) | 703.6 | 184.1 | 40 |
| SM(d18:1/18:0) | SM(d18:1(d9)/18:1) | 731.6 | 184.1 | 40 |
| SM(d18:1/24:0) | SM(d18:1(d9)/18:1) | 815.7 | 184.1 | 40 |
| SM(d18:1/24:1) | SM(d18:1(d9)/18:1) | 813.7 | 184.1 | 40 |
| Sphinganine | Sphingosine(d17:1) | 302.2 | 284.3 | 20 |
| Sphinganine 1-phosphate | Sphingosine 1-phosphate-d7 | 382.3 | 284.3 | 18 |
| Sphingosine | Sphingosine(d17:1) | 300.2 | 264.3 | 20 |
| Sphingosine 1-phosphate | Sphingosine 1-phosphate-d7 | 380.3 | 264.3 | 25 |
| Sphingosine 1-phosphate-d7 | IS | 387.3 | 271.3 | 25 |
| Sphingosine 1-phosphocholine | LPC(18:1(d7)) | 465.5 | 184.1 | 40 |
| Sphingosine(d17:1) | IS | 286.2 | 250.3 | 20 |
| TG(15:0/18:1(d7)/15:0) | IS | 829.8 | 570.8 | 40 |
| TG(18:0_36:2) | TG(15:0/18:1(d7)/15:0) | 904.7 | 603.4 | 40 |
| TG(18:1_34:2) | TG(15:0/18:1(d7)/15:0) | 874.7 | 575.4 | 40 |
| TG(18:1_34:3) | TG(15:0/18:1(d7)/15:0) | 872.7 | 573.4 | 40 |
| TG(20:4_32:1) | TG(15:0/18:1(d7)/15:0) | 870.6 | 549.3 | 40 |
| TG(20:4_34:2) | TG(15:0/18:1(d7)/15:0) | 896.6 | 575.3 | 40 |
| TG(20:4_34:3) | TG(15:0/18:1(d7)/15:0) | 894.6 | 573.3 | 40 |
| TG(20:4_36:0) | TG(15:0/18:1(d7)/15:0) | 928.8 | 607.5 | 40 |
| TG(20:4_36:2) | TG(15:0/18:1(d7)/15:0) | 924.7 | 603.4 | 40 |
| TG(20:4_36:3) | TG(15:0/18:1(d7)/15:0) | 922.7 | 601.4 | 40 |
| TG(22:6_36:2) | TG(15:0/18:1(d7)/15:0) | 948.7 | 603.4 | 40 |
| TG(22:6_38:1) | TG(15:0/18:1(d7)/15:0) | 978.7 | 633.4 | 40 |
| TG(22:6_38:2) | TG(15:0/18:1(d7)/15:0) | 976.7 | 631.4 | 40 |

### LRRK2 kinase activity regulates urinary BMP levels by modulating the release of BMP- and glycosphingolipid-containing vesicles from kidney

To better understand the mechanism by which LRRK2 activity regulates urinary BMP levels, we first explored the impact of LRRK2 deletion on BMP in urine and kidney from mice. Consistent with previous reports, we confirmed that BMP (22:6/22:6) levels were significantly reduced in urine from LRRK2 KO mice compared to WT controls (Fig. 1B)(29). It is possible that LRRK2 regulates BMP synthesis in the kidney and loss of LRRK2 would therefore result in a similar reduction of BMP in tissue. Alternatively, LRRK2 may drive the secretion of BMP-containing intraluminal vesicles through exosome release to the extracellular space, and loss of LRRK2 would instead result in the accumulation of BMP in kidney. To discriminate between these possibilities, we first performed imaging mass spectrometry (IMS) to assess the spatial distribution of BMP in kidneys from wildtype (WT) and LRRK2 KO mice, which revealed accumulation of BMP (22:6/22:6) within the renal cortex and outer medulla of LRRK2 KO mice compared to wildtype controls (Fig. 1C). Quantification of the levels of BMP in the homogenized tissues from dissected renal cortex and medulla using LC-MS/MS confirmed that BMP (22:6/22:6) levels were significantly increased in LRRK2 KO mouse kidney, with higher levels of BMP observed in the renal medulla consistent with our findings using IMS (Fig. 1D).

Our data suggests that LRRK2 may regulate exosome secretion more broadly and therefore impact levels of additional lipids enriched in ILVs beyond BMP, including BMP-related lipid and glycosphingolipid species (29). To test this, we performed targeted lipidomics on urine and kidney from WT and LRRK2 KO mice using LC-MS/MS. Similar to the reduction in BMP observed in urine from LRRK2 KO mice, glucosylceramide (GlcCer) levels were also decreased (Fig. 2A). In contrast, we observed a significant accumulation of multiple species of glycosphingolipids and components of the BMP pathway, including hemi-BMP, LPG and PG, in both kidney regions examined from LRRK2 KO mice (Fig 2B and C, Supplementary Table S2). Having demonstrated that loss of LRRK2 leads to an increase of BMP and glycosphingolipids in kidney and a corresponding reduction in urine, we next assessed how modulation of LRRK2 kinase activity impacts the levels of lipids enriched in exosomes. We hypothesized that genetic variants that increase LRRK2 kinase activity, such as the LRRK2 G2019S variant, would reduce BMP and glycosphingolipid levels in kidney and that LRRK2 kinase inhibition would, conversely, increase levels of these lipid species. To assess this, we dosed LRRK2 G2019S KI and WT littermates with either vehicle or a tool LRRK2 kinase inhibitor (MLi-2) in diet for 35 days and assessed the consequences on lipid levels in urine and kidney by mass spectrometry (Fig. 2D)(33, 34). Both pharmacokinetics (unbound concentration of the MLi-2) and LRRK2 pS935 analysis demonstrated high level (>90%) of LRRK2 kinase inhibition (Supplementary Fig. S1A-B). In contrast to the elevated levels of BMP observed in urine from human LRRK2 G2019S carriers, BMP levels were unchanged in urine from LRRK2 G2019S KI mice compared to WT littermates (Supplementary Fig. S1C). Consistent with previous findings in both preclinical and clinical studies, following LRRK2 kinase inhibition in both LRRK2 G2019S KI and WT mice, the levels of urine BMP were significantly reduced compared to vehicle treated mice (Fig. 2E, Supplementary Fig. S1C, Supplementary Table S2)(29, 30, 35). Beyond changes in BMP, several species of GlcCer were also reduced following LRRK2 kinase inhibition in mouse urine (Fig. 2E, Supplementary Table 2). In kidney, we observed a significant reduction in BMP and glycosphingolipid levels in LRRK2 G2019S mice compared to WT littermates (Fig. 2F and Supplementary Fig. S1D), and LRRK2 inhibition increased levels of BMP-related lipids and several classes of glycosphingolipids (Fig. 2F-G and Supplementary Table S2). These collective results suggest that the LRRK2-dependent secretion of BMP- and glycosphingolipid-containing vesicles from kidney is driven by LRRK2 kinase activity.

**Figure 2:**
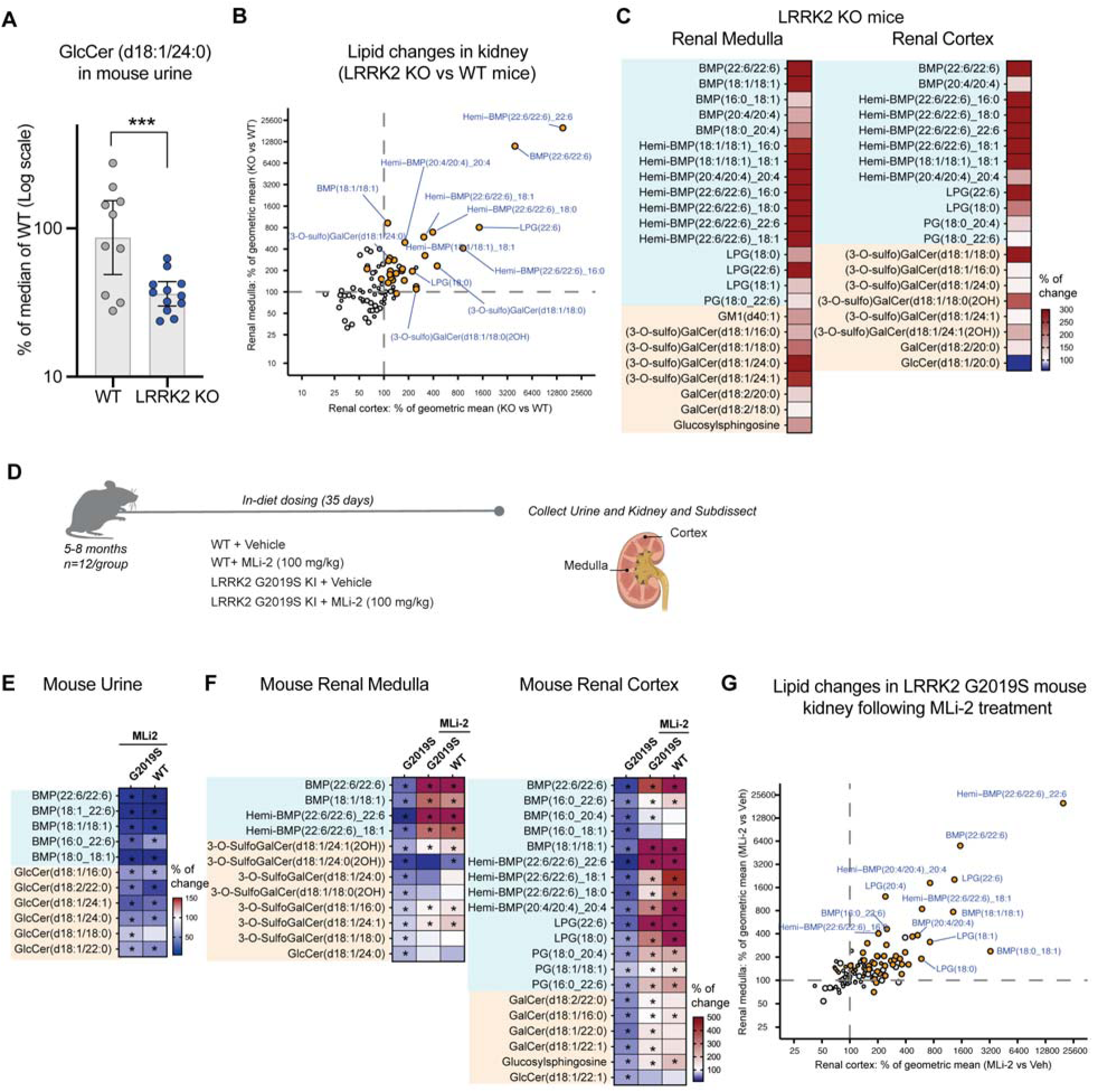
LRRK2 activity regulates the release of BMP and glycosphingolipids from kidney to urine. A) GlcCer (d18:1/24:0) levels in urine from LRRK2 KO mice (n=12) and WT littermates (n=10) were measured by LC-MS/MS. Data are presented as a percent of median values of the WT group, shown as geometric means with 95% CI, with p-values based on an ANCOVA model with statistical significance assessed using BH-adjusted p-values. ***P ≤ 0.001. B) The scatter plot represents the lipids changed in LRRK2 KO mice relative to WT littermates measured from renal cortex (X axis) and renal medulla (Y axis). The data are shown as estimated (geometric) mean ratio (%) for LRRK2 KO Vs WT, and the BMP related lipids and glycosphingolipids species were highlighted in orange if the BH-adjusted p-value for genotype difference was <0.05 for either the cortex or the medulla renal regions. Values > 200 fold increases were capped at 200X. C) A heatmap representing genotype-dependent differences on BMP related lipids and glycosphingolipids in the kidney from LRRK2 KO mice compared with WT control. The plot was generated as % of change by normalizing the average of LRRK2 KO group to the average of WT control group, in both renal medulla and renal cortex. Analytes included had BH-adjusted p-values <0.05 for genotype difference and were grouped based on lipid class, with BMP related lipids shaded in cyan, and glycosphingolipid species shaded in orange. White in the color scale depicts wild-type amounts, as 100%, red shows an accumulation (capped at 300%), and blue shows a reduction in the LRRK2 KO mice compared with WT control. D) Schematic of the in vivo dosing study design in LRRK2 G2019S knockin (KI) mice and WT littermates that were administered either a tool LRRK2 kinase inhibitor MLi-2 (100 mg/kg) or vehicle control in chow. A panel of lipid species were measured in urine and in the renal cortex and renal medulla 35 days after dosing using LC-MS/MS; n=12 animals per group. E) A heatmap representing the treatment effects of LRRK2 inhibitor MLi-2 on BMP-relevant species and glycosphingolipids in the urine from LRRK2 G2019S and WT mice, respectively. The plot was generated as % of change by normalizing the average for MLi-2-treatment group to the average of vehicle group of the same genotype (i.e. LRRK2 G2019S-MLi-2 vs. LRRK2 G2019S vehicle). Analytes included had BH-adjusted p-values <0.10 for treatment effect in LRRK2 G2019S mice and were grouped based on lipid class. *BH-adjusted p<0.10. White in the color scale depicts the vehicle control amounts, as 100%, red shows an accumulation, and blue shows a reduction in the treatment group. F) The heatmap demonstrates BMP-relevant species and glycosphingolipids are reduced in the kidney from LRRK2 G2019S KI mice compared to WT and are corrected upon treatment with MLi-2. The plot was generated as % of change by normalizing the average of different groups to the average of the WT vehicle group. The analytes included had nominal p-values <0.10 for genotype difference and were grouped based on lipid class. *p<0.10 annotated in the MLi-2 treatment groups are based on the MLi-2 vs. vehicle comparisons within the same genotype. White in the color scale depicts the WT-vehicle amounts, as 100%, red shows an accumulation (capped at 500%), and blue shows a reduction. G) The scatter plot demonstrates the degree of lipid changes upon MLi-2 treatment observed in LRRK2 G2019S KI mice relative to the LRRK2 G2019S KI-vehicle group from renal cortex (X axis) and renal medulla (Y axis). The data are shown as geometric mean ratio (%) for MLi-2 vs. vehicle, and the BMP-relevant species and glycosphingolipids were highlighted in orange if the BH-adjusted p-value for treatment effect was <0.05 for either the cortex or medulla renal regions. Values> 200 fold increases were capped at 200X.

**Table 2:** Lipidomics in negative mode parameters

| Name | Internal Standard | Q1 m/z | Q3 m/z | CE (V) |
| --- | --- | --- | --- | --- |
| (3-O-sulfo)GalCer(d18:1/16:0) | (3-O-sulfo)GalCer(d18:1/18:0(d3)) | 778.5 | 97 | -150 |
| (3-O-sulfo)GalCer(d18:1/18:0(2OH)) | (3-O-sulfo)GalCer(d18:1/18:0(d3)) | 822.6 | 97 | -150 |
| (3-O-sulfo)GalCer(d18:1/18:0(d3)) | IS | 809.6 | 97 | -150 |
| (3-O-sulfo)GalCer(d18:1/18:0) | (3-O-sulfo)GalCer(d18:1/18:0(d3)) | 806.6 | 97 | -150 |
| (3-O-sulfo)GalCer(d18:1/24:0(2OH)) | (3-O-sulfo)GalCer(d18:1/18:0(d3)) | 906.7 | 97 | -150 |
| (3-O-sulfo)GalCer(d18:1/24:0) | (3-O-sulfo)GalCer(d18:1/18:0(d3)) | 890.7 | 97 | -150 |
| (3-O-sulfo)GalCer(d18:1/24:1(2OH)) | (3-O-sulfo)GalCer(d18:1/18:0(d3)) | 904.7 | 97 | -150 |
| (3-O-sulfo)GalCer(d18:1/24:1) | (3-O-sulfo)GalCer(d18:1/18:0(d3)) | 888.7 | 97 | -150 |
| Arachidonic acid | Arachidonic acid-d8 | 303.2 | 303.2 | -10 |
| Arachidonic acid_MRM | Arachidonic acid-d8_MRM | 303.2 | 259.1 | -19 |
| Arachidonic acid-d8 | IS | 311.3 | 311.3 | -10 |
| Arachidonic acid-d8_MRM | IS | 311.3 | 267.1 | -19 |
| BMP(14:0/14:0) | IS | 665.3 | 227.2 | -50 |
| BMP(16:0_18:1) | BMP(14:0/14:0) | 747.5 | 255.4 | -50 |
| BMP(16:0_20:4) | BMP(14:0/14:0) | 769.5 | 255.4 | -50 |
| BMP(16:0_22:6) | BMP(14:0/14:0) | 795.5 | 255.4 | -50 |
| BMP(16:1/16:1) | BMP(14:0/14:0) | 717.5 | 253.1 | -50 |
| BMP(18:0_18:1) | BMP(14:0/14:0) | 775.5 | 281.4 | -50 |
| BMP(18:0_20:4) | BMP(14:0/14:0) | 797.5 | 283.4 | -50 |
| BMP(18:0_22:6) | BMP(14:0/14:0) | 823.5 | 283.4 | -50 |
| BMP(18:1/18:1) | BMP(14:0/14:0) | 773.5 | 281.3 | -50 |
| BMP(20:4/20:4) | BMP(14:0/14:0) | 817.5 | 303.3 | -50 |
| BMP(22:6/22:6) | BMP(14:0/14:0) | 865.5 | 327.3 | -50 |
| Cholesterol sulfate | (3-O-sulfo)GalCer(d18:1/18:0(d3)) | 465.3 | 96.7 | -80 |
| CL(14:0/14:0/14:0/14:0) | IS | 619.5 | 227.2 | -50 |
| CL(72:6/18:2) | CL(14:0/14:0/14:0/14:0) | 725.7 | 279.2 | -50 |
| CL(72:7/18:2) | CL(14:0/14:0/14:0/14:0) | 724.7 | 279.2 | -50 |
| CL(72:8-2(OOH)/18:2) | CL(14:0/14:0/14:0/14:0) | 755.7 | 279.2 | -50 |
| CL(72:8/18:2) | CL(14:0/14:0/14:0/14:0) | 723.7 | 279.3 | -50 |
| CL(74:9/18:2) | CL(14:0/14:0/14:0/14:0) | 736.7 | 279.2 | -50 |
| DHA | Arachidonic acid-d8_MRM | 327.2 | 229.1 | -19 |
| EPA | Arachidonic acid-d8_MRM | 301.3 | 257.1 | -19 |
| FAHFA(18:1/9-O-18:0) | PG(15:0/18:1(d7)) | 563.6 | 281 | -50 |
| GD1a/b(d36:1) | GM3(d18:1/18:0(d5)) | 917.5 | 290.1 | -65 |
| GD3(d34:1) | GM3(d18:1/18:0(d5)) | 720.9 | 290.1 | -65 |
| GD3(d36:1) | GM3(d18:1/18:0(d5)) | 734.9 | 290.1 | -65 |
| GM3(d18:1/18:0(d5)) | IS | 1184.8 | 290.1 | -65 |
| GM3(d34:1) | GM3(d18:1/18:0(d5)) | 1151.7 | 290.1 | -65 |
| GM3(d36:1) | GM3(d18:1/18:0(d5)) | 1179.8 | 290.1 | -65 |
| GQ1b(d36:1) | GM3(d18:1/18:0(d5)) | 1208.6 | 290.1 | -65 |
| Hemi-BMP(14:0/14:0)_14:0 | IS | 875.5 | 227.3 | -50 |
| Hemi-BMP(18:1/18:1)_16:0 | Hemi-BMP(14:0/14:0)_14:0 | 1011.7 | 281.3 | -50 |
| Hemi-BMP(18:1/18:1)_18:0 | Hemi-BMP(14:0/14:0)_14:0 | 1039.7 | 281.3 | -50 |
| Hemi-BMP(18:1/18:1)_18:1 | Hemi-BMP(14:0/14:0)_14:0 | 1037.7 | 281.3 | -50 |
| Hemi-BMP(20:4/20:4)_16:0 | Hemi-BMP(14:0/14:0)_14:0 | 1056.8 | 303.3 | -50 |
| Hemi-BMP(20:4/20:4)_18:0 | Hemi-BMP(14:0/14:0)_14:0 | 1185.8 | 303.3 | -50 |
| Hemi-BMP(20:4/20:4)_18:1 | Hemi-BMP(14:0/14:0)_14:0 | 1183.8 | 303.3 | -50 |
| Hemi-BMP(20:4/20:4)_20:4 | Hemi-BMP(14:0/14:0)_14:0 | 1104.8 | 303.3 | -50 |
| Hemi-BMP(22:6/22:6)_16:0 | Hemi-BMP(14:0/14:0)_14:0 | 1103.7 | 327.3 | -50 |
| Hemi-BMP(22:6/22:6)_18:0 | Hemi-BMP(14:0/14:0)_14:0 | 1131.7 | 327.3 | -50 |
| Hemi-BMP(22:6/22:6)_18:1 | Hemi-BMP(14:0/14:0)_14:0 | 1129.7 | 327.3 | -50 |
| Hemi-BMP(22:6/22:6)_22:6 | Hemi-BMP(14:0/14:0)_14:0 | 1175.7 | 327.3 | -50 |
| Linoleic acid | Arachidonic acid-d8 | 279.2 | 279.2 | -38 |
| Linolenic acid | Arachidonic acid-d8 | 277.2 | 277.2 | -10 |
| LPE(16:0) | LPE(18:1(d7)) | 452.2 | 255.3 | -50 |
| LPE(18:0) | LPE(18:1(d7)) | 480.31 | 283.3 | -50 |
| LPE(18:1(d7)) | IS | 485.3 | 288.3 | -50 |
| LPG(16:0) | LPE(18:1(d7)) | 483.3 | 255.3 | -50 |
| LPG(18:0) | LPE(18:1(d7)) | 511.3 | 283.3 | -50 |
| LPG(18:1) | LPE(18:1(d7)) | 509.3 | 281.3 | -50 |
| LPG(20:4) | LPE(18:1(d7)) | 531.3 | 303.3 | -50 |
| LPG(22:6) | LPE(18:1(d7)) | 555.3 | 327.3 | -50 |
| LPI(16:0) | LPE(18:1(d7)) | 571.3 | 241.1 | -50 |
| LPI(18:0) | LPE(18:1(d7)) | 599.3 | 241.1 | -50 |
| Oleic acid | Arachidonic acid-d8 | 281.2 | 281.2 | -10 |
| PA(15:0/18:1(d7)) | IS | 666.52 | 241.3 | -50 |
| PA(16:0_18:1) | PA(15:0/18:1(d7)) | 673.5 | 255.3 | -50 |
| PA(18:0_18:1) | PA(15:0/18:1(d7)) | 701.5 | 283.3 | -50 |
| PA(18:0_20:4) | PA(15:0/18:1(d7)) | 723.5 | 283.3 | -50 |
| PA(18:0_22:6) | PA(15:0/18:1(d7)) | 747.5 | 283.3 | -50 |
| PA(18:1/18:1) | PA(15:0/18:1(d7)) | 699.5 | 281.3 | -50 |
| Palmitic acid | Arachidonic acid-d8 | 255.1 | 255.1 | -10 |
| Palmitoleic acid | Arachidonic acid-d8 | 253.1 | 253.1 | -10 |
| PE(15:0/18:1(d7)) | IS | 709.6 | 241.3 | -50 |
| PE(O-16:0/20:4) | PE(15:0/18:1(d7)) | 724.5 | 303.2 | -50 |
| PE(O-16:0/22:6) | PE(15:0/18:1(d7)) | 748.5 | 327.2 | -50 |
| PE(O-18:0/20:4) | PE(15:0/18:1(d7)) | 752.6 | 303.2 | -50 |
| PE(O-18:0/22:6) | PE(15:0/18:1(d7)) | 776.6 | 327.2 | -50 |
| PE(P-16:0/20:4) | PE(15:0/18:1(d7)) | 722.6 | 303.3 | -50 |
| PE(P-16:0/20:5) | PE(15:0/18:1(d7)) | 720.6 | 301.3 | -50 |
| PE(P-16:0/22:4) | PE(15:0/18:1(d7)) | 750.6 | 331.3 | -50 |
| PE(P-16:0/22:6) | PE(15:0/18:1(d7)) | 746.6 | 327.3 | -50 |
| PE(P-18:0/18:1) | PE(15:0/18:1(d7)) | 728.6 | 281.3 | -50 |
| PE(P-18:0/18:2) | PE(15:0/18:1(d7)) | 726.6 | 279.2 | -50 |
| PE(P-18:0/20:4) | PE(15:0/18:1(d7)) | 750.6 | 303.3 | -50 |
| PE(P-18:0/20:5) | PE(15:0/18:1(d7)) | 748.6 | 301.3 | -50 |
| PE(P-18:0/22:6) | PE(15:0/18:1(d7)) | 774.6 | 327.3 | -50 |
| PE(P-18:1/20:4) | PE(15:0/18:1(d7)) | 748.5 | 303.3 | -50 |
| PE(P-18:1/22:6) | PE(15:0/18:1(d7)) | 772.5 | 327.3 | -50 |
| PEth(16:0_18:1) | PE(15:0/18:1(d7)) | 772.5 | 255.1 | -50 |
| PEth(18:1/18:1) | PE(15:0/18:1(d7)) | 773.6 | 281.2 | -50 |
| PG(15:0/18:1(d7)) | IS | 740.55 | 241.3 | -50 |
| PG(16:0_18:1) | PG(15:0/18:1(d7)) | 747.5 | 255.3 | -50 |
| PG(16:0_20:4) | PG(15:0/18:1(d7)) | 769.5 | 255.3 | -50 |
| PG(16:0_22:6) | PG(15:0/18:1(d7)) | 795.5 | 255.3 | -50 |
| PG(18:0_18:1) | PG(15:0/18:1(d7)) | 775.5 | 281.3 | -50 |
| PG(18:0_20:4) | PG(15:0/18:1(d7)) | 797.5 | 283.3 | -50 |
| PG(18:0_22:6) | PG(15:0/18:1(d7)) | 823.5 | 283.3 | -50 |
| PG(18:1/18:1) | PG(15:0/18:1(d7)) | 773.4 | 281.4 | -50 |
| PI(15:0/18:1(d7)) | IS | 828.6 | 241.3 | -50 |
| PI(16:0_18:1) | PI(15:0/18:1(d7)) | 835.6 | 255.3 | -50 |
| PI(16:0_20:4) | PI(15:0/18:1(d7)) | 857.6 | 255.3 | -50 |
| PI(16:0_22:6) | PI(15:0/18:1(d7)) | 881.6 | 255.3 | -50 |
| PI(18:0_18:1) | PI(15:0/18:1(d7)) | 863.6 | 283.3 | -50 |
| PI(18:0_20:4) | PI(15:0/18:1(d7)) | 885.6 | 283.3 | -50 |
| PI(18:0_22:6) | PI(15:0/18:1(d7)) | 909.6 | 283.3 | -50 |
| PI(18:1/18:1) | PI(15:0/18:1(d7)) | 861.6 | 281.3 | -50 |
| PI(20:4/20:4) | PI(15:0/18:1(d7)) | 905.6 | 303.3 | -50 |
| PS(15:0/18:1(d7)) | IS | 753.55 | 241.3 | -50 |
| PS(16:0_18:1) | PS(15:0/18:1(d7)) | 760.6 | 255.3 | -50 |
| PS(16:0_20:4) | PS(15:0/18:1(d7)) | 782.6 | 255.3 | -50 |
| PS(16:0_22:6) | PS(15:0/18:1(d7)) | 806.6 | 255.3 | -50 |
| PS(18:0_18:1) | PS(15:0/18:1(d7)) | 788.6 | 283.3 | -50 |
| PS(18:0_20:4) | PS(15:0/18:1(d7)) | 810.6 | 283.3 | -50 |
| PS(18:0_22:6) | PS(15:0/18:1(d7)) | 834.6 | 283.3 | -50 |
| PS(18:1/18:1) | PS(15:0/18:1(d7)) | 786.6 | 281.3 | -50 |
| PS(22:6/22:6) | PS(15:0/18:1(d7)) | 878.5 | 327.3 | -50 |
| Stearic acid | Arachidonic acid-d8 | 283.2 | 283.2 | -10 |

Note: IS - Internal standard;
The percent of samples with missing area ratio is calculated across the entire set of samples, including the pooled control samples.

We next directly assessed the impact of LRRK2 kinase activity on extracellular vesicle secretion in human subjects dosed with a LRRK2 inhibitor from our clinical study. Using urine samples obtained from human subjects dosed with the LRRK2 kinase inhibitor DNL201 in our recent phase 1b clinical study, we isolated extracellular vesicles (EVs) and analyzed their lipid content via mass-spectrometry(30). BMP was readily detected from EVs isolated from human urine, confirming that BMP is enriched in urinary EVs. We observed a trend toward reduction in total lipid levels and in BMP specifically in EVs isolated from urine from subjects treated with DNL201(n=11) compared to the placebo group (n=7; Supplementary Fig. S1E). While the sample size was limited and not specifically powered for this assessment, these data are consistent with our hypothesis that LRRK2-dependent regulation of exosome release may be a key mechanism by which LRRK2 inhibition reduces BMP levels in urine.

### LRRK2 kinase activity regulates glycosphingolipid and BMP levels in a cell type-specific manner in mouse brain

While urinary BMP levels have been employed as a peripheral biomarker to demonstrate LRRK2-dependent regulation of lysosomal biology in clinical studies, it is unclear whether LRRK2 similarly regulates BMP levels in the CNS. We first assessed the levels of BMP (22:6/22:6) throughout the brain of LRRK2 KO mice and WT littermates using imaging mass spectrometry. Overall, we did not observe gross differences in brain BMP levels between WT and LRRK2 KO mice, in contrast to the effects observed in the kidney (Fig. 3A, Supplementary Fig. S2A-C). While this global analysis suggests that LRRK2 does not profoundly impact BMP levels in the brain, we reasoned that LRRK2 may impact BMP in a cell-type specific manner given the low expression of the kinase throughout the CNS. Previous analyses suggest that LRRK2 expression is highest in mouse astrocytes, with moderate expression in mouse microglia and lower expression in cortical neurons(19, 25). With this in mind, we employed a fluorescence-activated cell sorting (FACS)-based method to isolate enriched populations of astrocytes, microglia, and neurons from LRRK2 KO and LRRK2 G2019S KI mice and assessed lipid changes using mass-spectrometry(36). We observed the most profound lipid changes in astrocytes from LRRK2 KO mice compared to other CNS cell types, with reductions in several classes of glycosphingolipids including GalCer and a trend toward an elevation in BMP (22:6/22:6) (Fig 3B-D, Supplementary Fig S2D-E, Table 2). Alterations in a small number of species of glycosphingolipids and components of the BMP pathway were also observed in microglia and neurons from LRRK2 KO mice (Supplementary Fig. 2D and E).

**Figure 3:**
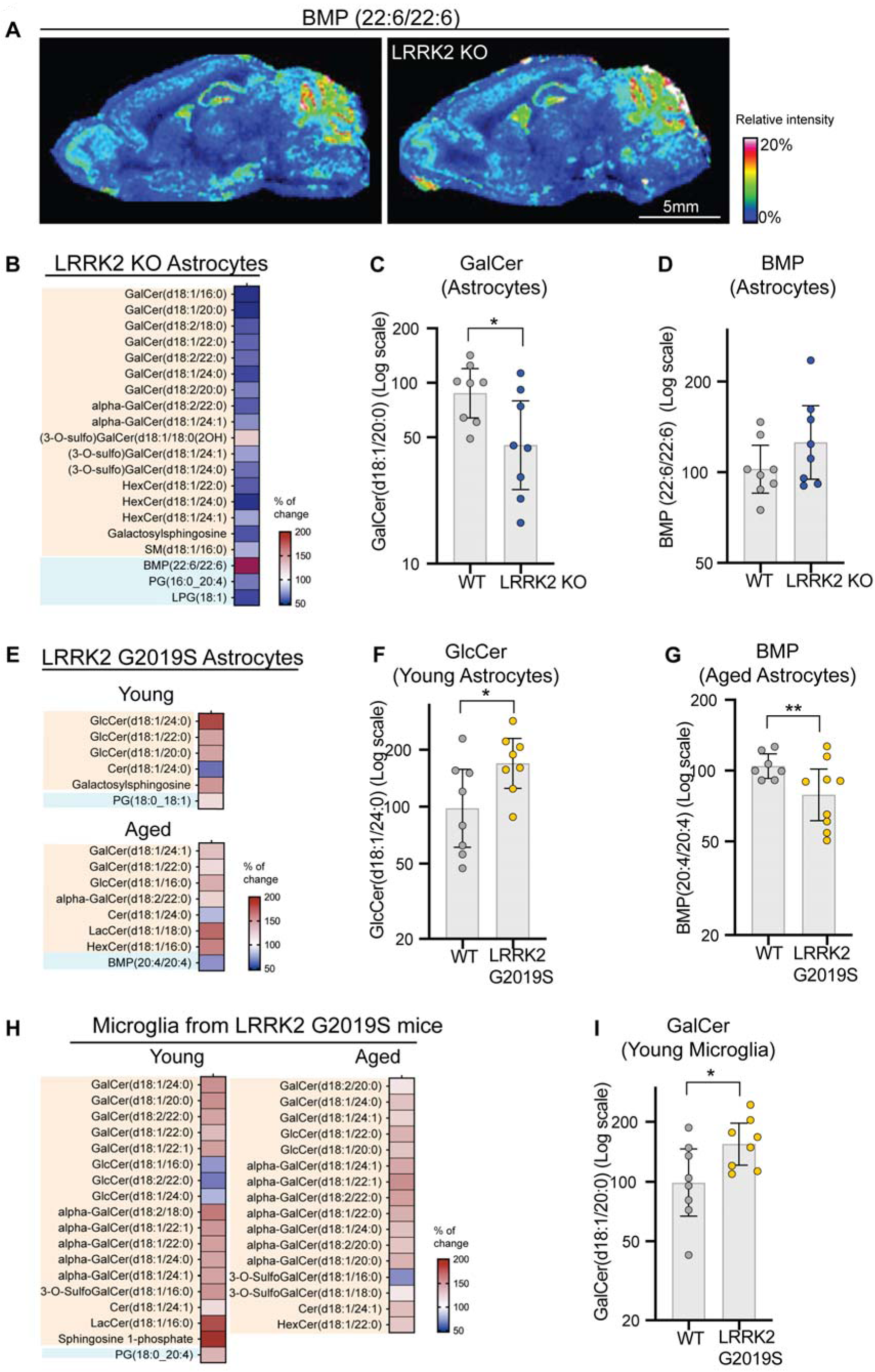
LRRK2 activity regulates glycosphingolipids in mouse brain with modest changes in BMP. A) Matrix-assisted laser desorption/ionization mass spectrometry images were acquired from sagittal brain sections of WT and LRRK2 KO mice. Representative mass spectrometry image showing the distribution of the signal at a mass/charge ratio (m/z) of 865.502, corresponding to BMP (22:6/22:6). Images depict the relative intensity of the signal from 0 to 20% with intensity normalized using total ion current. B) A heatmaps represented the profiles of BMP related lipids and glycosphingolipids in astrocytes from LRRK2 KO mice and LRRK2 G2019S KI mice at 5-6 month-old (young) and 18-month-old (aged); n=8 animals for LRRK2 KO mice and WT littermates. C-D) Representative plots are shown for GalCer (d18:1/20:0) (C) and BMP (22:6/22:6) (D) in astrocytes from LRRK2 KO mice. BMP showed a trend of increase in LRRK2 KO astrocyte with p= 0.064. E) Heatmaps represented the profiles of BMP related lipids and glycosphingolipids in astrocytes from LRRK2 G2019S KI mice at 5-6 month-old (young) and 18-month-old (aged); n=8 animals for LRRK2 G2019S KI and WT littermates at 5–6 month-old; n=7 for WT littermates and n=9 for LRRK2 G2019S KI mice at 18 month-old. (F-G) Representative plots are shown for GlcCer (d18:1/24:0) (F) and BMP (20:4/20:4) in LRRK2 G2019S astrocytes (G). H) Heatmaps represented the profiles of BMP related lipids and glycosphingolipids in microglia from LRRK2 G2019S KI mice at 5-6 month-old (young) and 18-month-old (aged). I) Representative plots are shown for GalCer (d18:1/20:0) in LRRK2 G2019S microglia. The heatmaps (B, E and H) were generated as % of change normalized to WT littermate controls. The analytes included had nominal p-values <0.10 for genotype difference and were grouped based on lipid class. The BMP related lipids were shaded in cyan, and the glycosphingolipid species were shaded in orange. In the color scale, white depicts the WT amounts, as 100%, red shows an accumulation and blue shows a reduction. For plots (C, D, F, G and I), the relative abundance of the analytes was normalized to the median values of WT group. Data are plotted in log scale and shown as geometric mean ratio (%) and 95% CIs with statistical significance assessed at nominal levels. *p < 0.05, **p < 0.01.

To assess the impact of a pathogenic kinase-activating LRRK2 variant on BMP and glycosphingolipids in brain, we performed a similar targeted lipidomic analysis from FACS-enriched CNS cells from LRRK2 G2019S KI mice. We measured lipid levels in CNS cells from young (5-6 months) and old (18 months) LRRK2 G2019S mice to assess whether aging further impacted LRRK2 G2019S-dependent effects on lipids relevant to lysosomal function. Many glycosphingolipids accumulated in astrocytes and microglia from LRRK2 G2019S mice in both age groups assessed (Fig. 3E, H, Supplementary Table 2). Several species of GlcCer and GalCer were significantly elevated in glial cells from LRRK2 G2019S mice, in contrast to the reduction of these lipid species observed in glial cells from LRRK2 KO mice (Fig. 3F, G and I).

Glycosphingolipid dysregulation was most apparent in microglia from LRRK2 G2019S mice, consistent with emerging data from LSDs that suggest that microglia are particularly susceptible to defects in lysosomal function(36, 37). Changes in glycosphingolipids were more modest in neurons where major alterations were only observed upon aging (Supplementary Fig. S2F). BMP levels were reduced in aged LRRK2 G2019S astrocytes and neurons, which contrasted with the elevation of BMP observed in astrocytes from LRRK2 KO mice (Fig. 3B, D, E and G and Supplementary Fig. S2F). Overall, these data establish that LRRK2 regulates BMP levels in specific CNS cell types and plays a major role in regulating glycosphingolipid levels in the brain. As glycosphingolipid alterations were apparent in young LRRK2 G2019S mice and BMP changes were only observed upon aging, these data also suggest that LRRK2-dependent glycosphingolipid dysregulation may contribute to effects on BMP in the brain.

### LRRK2 kinase activity regulates glucosylceramide and BMP levels to maintain proper lysosomal function in cellular models

To better understand the mechanism(s) by which LRRK2 kinase activity modulates glycosphingolipids and BMP, we assessed the impact of a pathogenic LRRK2 variant on lipid accumulation and lysosomal function in A549 cells. This cell line was selected for functional studies based on its high expression of both LRRK2 and Rab10, a previously-identified LRRK2 substrate(21). To enable our mechanistic studies, we focused on the LRRK2 R1441G variant given its strong impact on Rab phosphorylation(38). We found that levels of several species in the BMP pathway, including BMP (22:6/22:6), were significantly elevated in cells expressing the LRRK2 R1441G variant compared to WT cells (Fig. 4A and Supplementary Fig. S3). Further, glycosphingolipid levels were also broadly elevated in LRRK2 R1441G cells compared to WT cells, including nearly all species of GlcCer (Fig. 4B). To determine whether LRRK2 kinase activity regulates glycosphingolipid accumulation in LRRK2 R1441G cells, we treated cells with the LRRK2 kinase inhibitor DNL151 (2 μM) for 72 hours and assessed GlcCer levels by LC-MS/MS. DNL151 treatment fully normalized GlcCer levels in LRRK2 R1441G cells (Fig. 4C). While these data show that LRRK2 kinase inhibition can rescue glycosphingolipid accumulation, the extent to which LRRK2 kinase activity needs to be inhibited to correct lipid storage was unclear. We treated LRRK2 R1441G KI cells with increasing doses of DNL151 to determine what level of LRRK2 kinase inhibition, as assessed by the levels of phosphorylated Rab10, was required to normalize glycosphingolipid levels. Our data showed that approximately 80% LRRK2 kinase inhibition was required to reduce GlcCer levels to that observed in WT cells (Fig. 4D-E). These data are consistent with recent clinical findings with a related LRRK2 inhibitor (DNL201), which show that >80% LRRK2 kinase inhibition is needed to reduce BMP levels in human urine and suggest that this level of inhibition may be necessary to correct lysosomal dysfunction mediated by LRRK2 hyperactivity (30).

**Figure 4:**
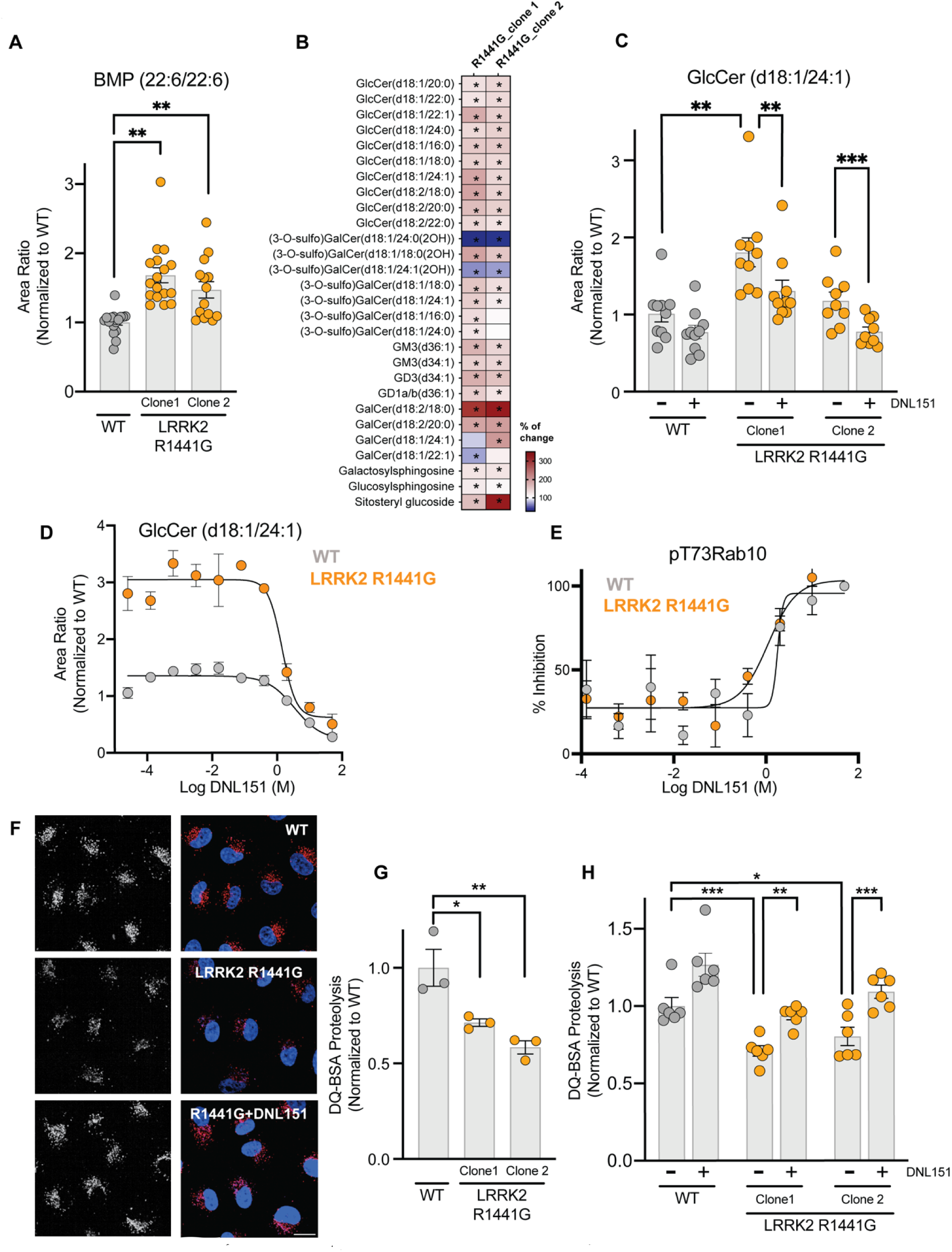
LRRK2 activity regulates glucosylceramide and BMP levels to maintain proper lysosomal function. A) BMP (22:6/22:6) levels were measured in WT parental cells and two clones of LRRK2 R1441G KI A549 cells; data are shown as mean ± SEM; n = 17 independent experiments, and statistical significance was determined using one-way ANOVA following log transformation B) Heatmap showing elevated glycosphingolipids including multiple GlcCer species in whole cell extracts from LRRK2 R1441G KI cells as compared to parental wild-type cells. The heatmaps were generated as % of change by normalizing the average of different groups to the average of the WT group. The analytes included had nominal p-values <0.10 for genotype difference in either clone and were grouped based on lipid class. *p<0.10. White in the color scale depicts the WT-vehicle amounts, as 100%, red shows an accumulation (capped at 350%), and blue shows a reduction transformation. C) WT and LRRK2 R1441G KI A549 cells were treated with vehicle or DNL151 (2μμ) for 72 hours, and the levels of GlcCer (d18:1/24:1) were measured using LC-MS/MS. Data are shown as mean ± SEM; n = 10 independent experiments, and statistical significance was determined using one-way ANOVA following log transformation. D) WT and LRRK2 R1441G KI A549 cells were treated with increasing concentrations of DNL151, and the levels of GlcCer (d18:1/24:1) were measured using LC-MS/MS. Data are shown as mean ± SEM; n = 4 independent experiments, and statistical significance was determined using non-linear fit following log transformation. E) Dose-response curves show the percent inhibition of LRRK2 kinase activity as measured by levels of phosphorylated T73 Rab10 in WT and LRRK2 R1441G cells. F) Representative images of DQ-BSA signals (left panels: black and white, right panels: red) in WT A549 cells (top panel), LRRK2 R1441G KI cells (middle panel) and LRRK2 R1441G KI cells treated with DNL151(bottom panel). Nuclei stained with NucBlue (blue); scale bar = 10 tsm. G) The sum of spot intensities of DQ-BSA signal was quantified per cell. The DQ-BSA signals were normalized to the median within each experiment and to the WT control. Data are shown as mean ± SEM; n=3 independent experiments, and statistical significance was determined using one-way ANOVA following log transformation. H) WT and LRRK2 R1441G cells were treated with vehicle or DNL151 (2μM) for 72 hours, and lysosomal proteolysis was measured using the DQ-BSA-based assay. The sum spot intensities of DQ-BSA signal were quantified per cell. The DQ-BSA signals were normalized to the median within each experiment and to the WT vehicle control. Data are shown as mean ± SEM; n = 6 independent experiments, and statistical significance was determined using one-way ANOVA following log transformation. *P < 0.05, **P < 0.01, ***P < 0.001

To understand the functional impact of BMP and glycosphingolipid accumulation, we next assessed the impact of the pathogenic LRRK2 R1441G variant on lysosomal homeostasis more directly using de-quenched bovine serum albumin (DQ-BSA) to monitor lysosomal proteolysis. Presence of the LRRK2 R1441G variant reduced the endolysosomal turnover of DQ-BSA by approximately 30-40% compared to WT cells, suggesting that increased LRRK2 activity impairs the turnover of proteins via the lysosome in these cells (Fig. 4F and G). Treatment with the LRRK2 kinase inhibitor DNL151 (2 μM) fully rescued lysosomal proteolysis, normalizing signal to that observed in WT cells (Fig. 4H). These data demonstrate that mutant LRRK2 perturbs lysosomal proteolysis and triggers lipid storage in cells and establish that LRRK2 kinase inhibition can correct lipid accumulation that correlates with broad improvements in lysosomal function.

### Endolysosomal GCase activity, GlcCer and BMP levels are controlled by LRRK2 kinase activity

Given our results demonstrating that LRRK2 pathogenic variants impair lysosomal proteolysis and cause elevation of lipids that are known to accumulate upon lysosomal dysfunction, we next assessed whether LRRK2 activity impacts lipid homeostasis and hydrolase activity specifically in the lysosome. We employed an established lysosome immunoprecipitation (Lyso-IP) method that enables rapid isolation of intact lysosomes to profile how LRRK2 activity regulates lysosomal lipids using mass-spectrometry(39, 40). Similar to our observations from whole cell lysates, lysosomes isolated from LRRK2 R1441G KI cells showed broad elevations in GlcCer species and in BMP (20:4/20:4) compared to lysosomes from WT cells (Fig. 5A-C). DNL151 treatment fully normalized GlcCer and BMP (20:4/20:4) accumulation in LRRK2 R1441G cells to WT levels, demonstrating that LRRK2 inhibition can rescue lipid accumulation in lysosomes (Fig. 5B and C).

**Figure 5:**
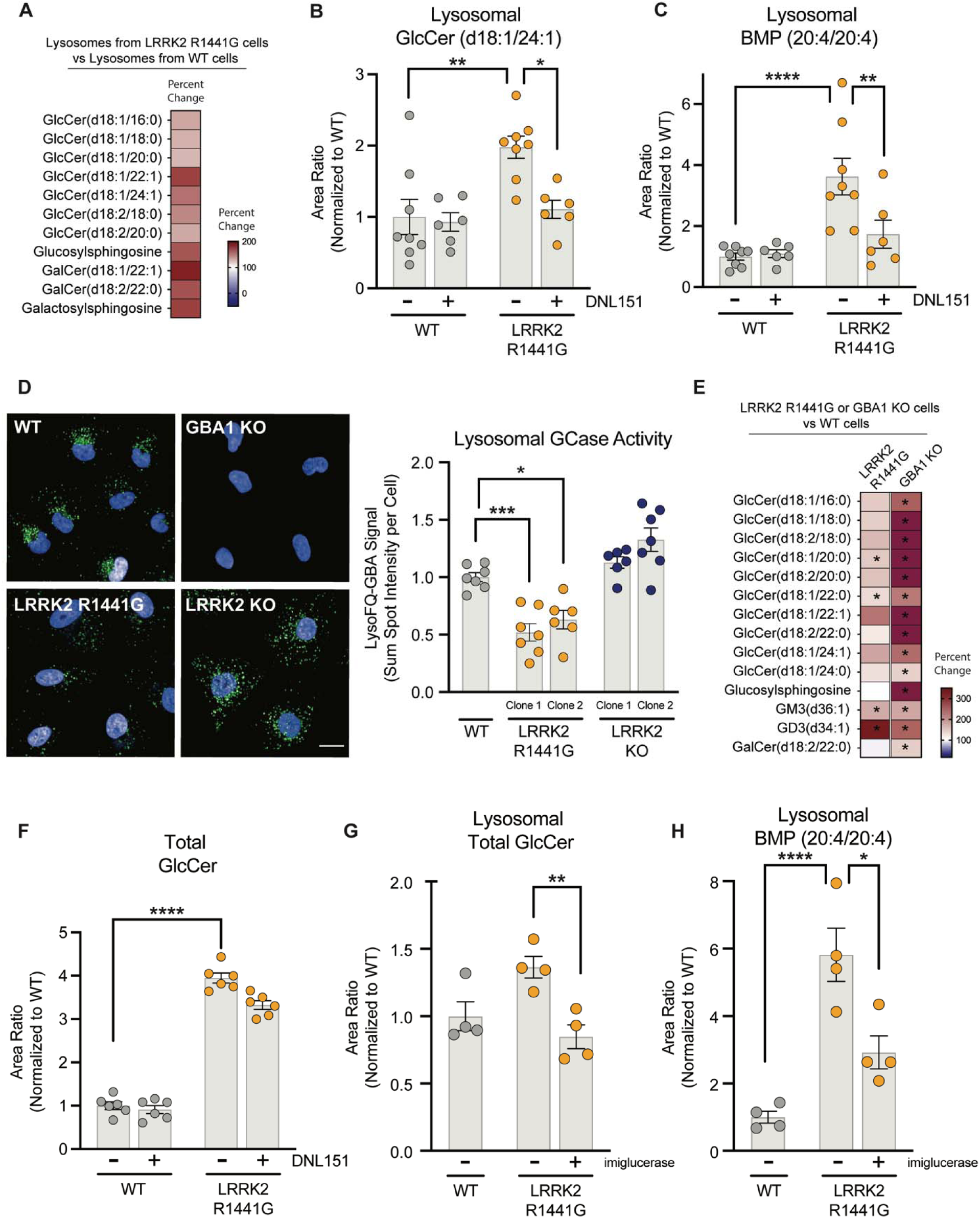
LRRK2 activity regulates endolysosomal GCase activity and GlcCer and BMP levels in lysosomes. A) The levels of glycosphingolipid species were measured in lysosomes isolated from WT and LRRK2 R1441G KI A549 cells. The percent change in signal from LRRK2 R1441G cells compared to WT cells is shown in the heatmap. The analytes included had nominal p-values <0.10 for genotype difference and were grouped based on lipid; n = 12 independent experiments class; n=12 independent experiments. B and C) WT and LRRK2 R1441G KI A549 cells were treated with vehicle or DNL151 (2μμ) for 72 hours, lysosomes were rapidly immunoprecipitated, and the levels of GlcCer (d18:1/24:1) and BMP (20:4/20:4) were measured using LC-MS/MS. Data are shown as mean ± SEM; n = 8 independent experiments, and statistical significance was determined using one-way ANOVA following log transformation. D) Lysosomal GCase activity (shown in green) was assessed using the LysoFQ-GBA probe in WT, GBA1 KO, LRRK2 R1441G KI, and LRRK2 KO A549 cells, nuclei were visualized using NucBlue staining (shown in blue). Representative images are shown; scale bar = 10 μm. The sum of the spot intensities per cell was quantified; data are shown as mean ± SEM; n=7 independent experiments, and statistical significance was determined using one-way ANOVA following log transformation. E) GSL profiling of WT, LRRK2 R1441G KI, and GBA1 KO A549 cells was performed, and the percent change was measured by normalizing the average of each group to the average of WT cells. The analytes included had nominal p-values <0.10 for genotype difference for either LRRK2 R1441G Vs WT or GBA1 KO Vs WT, and were grouped based on lipid class; n=3 independent experiments. * P<0.10. White in the color scale depicts the WT reference level as 100%, red shows an accumulation, and blue shows a reduction. The red scale in was capped at 350%; n=3 independent experiments. F) WT and GBA1 KO A549 cells were treated with vehicle or DNL151 (2μμ) for 72 hours, and the levels of GlcCer species were measured using LC-MS/MS. The sum of all GlcCer species was measured, and data are shown as mean ± SEM; n = 6 independent experiments, and statistical significance was determined using one-way ANOVA following log transformation. G and H) WT and LRRK2 R1441G KI A549 cells were treated with vehicle or imiglucerase (2μM) for 72 hours, and the levels of GlcCer and BMP (20:4/20:4) were measured using LC-MS/MS. Data are shown as mean ± SEM; n = 4 independent experiments, and statistical significance was determined using one-way ANOVA following log transformation. *P < 0.05, **P < 0.01, ***P < 0.001. ****P < 0.0001.

As elevated LRRK2 activity increased the lysosomal levels of GlcCer, a substrate of the lysosomal hydrolase GCase, we hypothesized that the pathogenic LRRK2 R1441G variant impaired GCase activity. Several studies have assessed the impact of LRRK2 variants on GCase activity and reported conflicting results(41-43). A clear understanding of the relationship between LRRK2 and lysosomal GCase activity has been hindered by limitations in existing methods to measure GCase activity with respect to sensitivity, selectivity, and spatial resolution. We employed a recently-described activity-based probe to measure lysosomal GCase activity (LysoFQ-GBA) that has superior signal to noise and retention in lysosomes compared to other probes tested to address this question (44). We confirmed that LysoFQ-GBA specifically detected GCase signal as no detectable signal was observed in *GBA1* KO A549 cells (Fig. 5D). Lysosomal GCase activity was reduced by 40-50% in LRRK2 R1441G cells compared to WT cells and showed a trend toward elevation in LRRK2 KO cells, supporting our hypothesis that LRRK2 regulates GlcCer levels by regulating GCase activity or lysosomal trafficking (Fig. 5D). Further suggesting that lysosomal lipid accumulation in LRRK2 R1441G cells results from impaired GCase activity, we observed a strikingly similar profile of lipid dysregulation in LRRK2 R1441G cells compared to GBA1 KO A549 cells (Fig. 5E). To determine whether LRRK2 regulated GCase activity by controlling its delivery to lysosomes, we assessed the levels of GCase in isolated lysosomes by western blot analysis. GCase protein levels were not changed in lysosomes isolated from LRRK2 R1441G clones and were significantly elevated in lysosomes isolated from LRRK2 KO A549 cells (Supplementary Fig. S3B-C). These data show that the LRRK2 R1441G variant does not impact the delivery of GCase to lysosomes and suggest that LRRK2 regulates additional factors required for optimal GCase activity. Together, these data suggest that pathogenic LRRK2 causes glycosphingolipid and BMP accumulation in lysosomes by impacting GCase function.

To better understand how LRRK2-dependent regulation of GCase activity contributes to its effects on GlcCer and BMP levels, we next determined whether GCase was necessary for LRRK2’s effects on lipid regulation in the lysosome. First, we assessed whether LRRK2 kinase inhibition required GCase activity to effectively reduce GlcCer accumulation by treating GBA1 KO cells with DNL151 (2 μM) for 72 hours and measuring its effect on GlcCer levels. GBA1 KO cells showed significant elevations in GlcCer levels compared to WT cells, and LRRK2 inhibition failed to significantly impact GlcCer accumulation (Fig. 5F). These data suggest that GCase is essential for LRRK2-dependent lysosomal GlcCer regulation. To directly test whether restoration of GCase activity was sufficient to rescue LRRK2-medaited GlcCer accumulation, we treated LRRK2 R1441G cells with recombinant GCase (imiglucerase) and evaluated its effects on lipids in isolated lysosomes using mass spectrometry. Imiglucerase treatment attenuated GlcCer accumulation in isolated lysosomes from LRRK2 R1441G cells (Fig. 5G). Interestingly, imiglucerase treatment also fully normalized BMP (20:4/20:4) levels in lysosomes isolated from LRRK2 R1441G cells (Fig. 5H), suggesting that BMP accumulation is a secondary consequence of reduced GCase activity. Together, our results show that LRRK2 regulates lysosomal BMP as a downstream response to its effect on GCase activity and reveal a novel mechanism by which LRRK2 regulates lysosomal BMP.

### LRRK2 regulates GCase activity and glycosphingolipid levels in human iPSC-derived microglia

To assess whether LRRK2-dependent regulation of GCase activity and BMP occurs in a disease-relevant cell type, we explored the impact of LRRK2 in human iPSC-derived microglia (iMicroglia). Although strong lipid alterations were observed with elevated kinase activity in both cell types *in vivo*, our studies focused specifically on microglia rather than astrocytes given the recent data highlighting that microglia-specific regulatory elements may drive the effects of non-coding LRRK2 risk variants in humans (45, 46). To further expand upon our studies in A549 cells, we examined the consequences of a second LRRK2 pathogenic variant (LRRK2 G2019S) on lysosomal function. We showed that expression of LRRK2 G2019S in iMicroglia resulted in an approximately two-fold increase in LRRK2 activity, as assessed by levels of Rab10 phosphorylation per molecule of LRRK2 (Fig. 6A and Supplementary Fig. S4A-C). Significant accumulation of the GCase substrate glucosylsphingosine (GlcSph) was observed in LRRK2 G2019S iMicroglia while GlcSph levels were reduced in LRRK2 KO iMicroglia compared to WT cells (Fig. 6B). Unlike in our A549 cell studies, we did not observe an alteration in GlcCer levels with expression of a pathogenic LRRK2 variant which may reflect that iMicroglia have greater levels of acid ceramidase (ASAH1) activity, an enzyme that converts GlcCer to glucosylsphingosine (Supplementary Fig. S4D). Consistent with the effects we observed in A549 cells with another pathogenic LRRK2 variant, the LRRK2 G2019S variant increased levels of BMP (20:4/20:4) and LRRK2 KO led to a trend toward reduction in BMP levels in iMicroglia (Fig. 6C). We examined lysosomal GCase activity using the LysoFQ-GBA substrate and confirmed its specificity in iMicroglia by demonstrating loss of signal following treatment with the GCase inhibitor (CBE) (Fig. 6D). Mirroring the effects of LRRK2 observed on glucosylsphingosine, we observe an approximately 50% reduction in lysosomal GCase activity in LRRK2 G2019S iMicroglia and an approximately 50% increase in GCase activity in LRRK2 KO iMicroglia (Fig. 6D). Treatment with recombinant GCase fully rescued glucosylsphingosine and BMP accumulation, demonstrating that LRRK2 acts upstream of GCase to regulate BMP through a similar mechanism in iMicroglia as that observed in A549 cells (Fig. 6E and F). Together, our data show that LRRK2 regulates GCase activity and substrate catabolism and BMP levels in human microglia, demonstrating that LRRK2-dependent regulation of lysosomal function occurs in a key CNS cell type relevant for PD.

**Figure 6:**
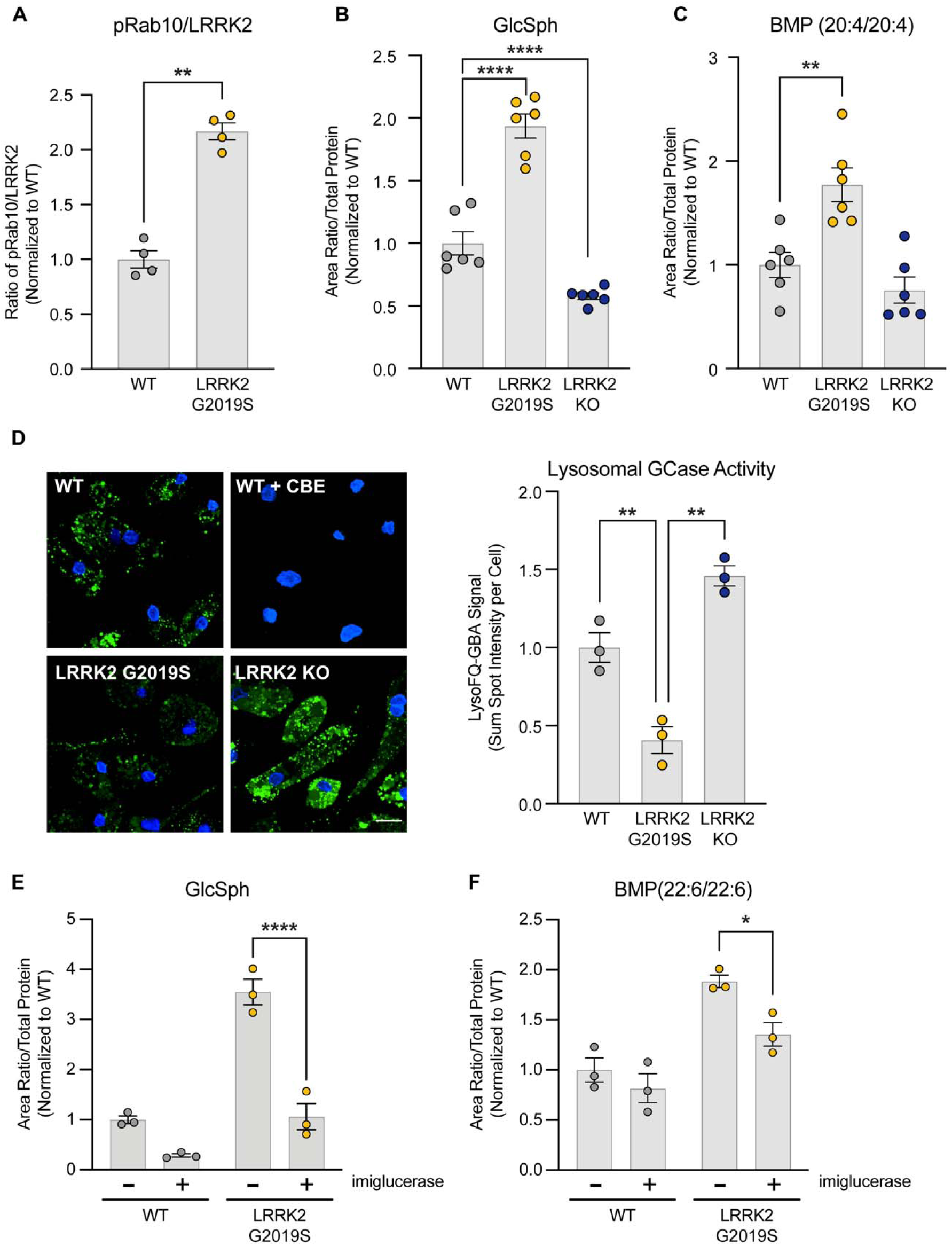
LRRK2 regulates GCase activity and levels of BMP and glucosylsphingosine in human iPSC-derived microglia. A) pT73 Rab10 levels and total LRRK2 were quantified from WT and LRRK2 G2019S KI iMicroglia using MSD-based assays. The ratio of pRab10/LRRK2 levels were quantified, and data are shown as geometric mean ± SEM; n=4 independent experiments, and statistical significance was determined using Student’s t-test. (B and C) The levels of GlcSph (B) and BMP (20:4/20:4) (C) were measured using LC-MS/MS in cell lysates from WT, LRRK2 G2019S KI and LRRK2 KO iMicroglia. Data are shown as geometric mean ± SEM; n=6 independent experiments, and statistical significance was determined using one-way ANOVA and Dunnett’s multiple comparison. (D) Lysosomal GCase activity was assessed using the LysoFQ-GBA probe in WT cells, WT cells treated with the GCase inhibitor CBE, LRRK2 G2019S KI, and LRRK2 KO iMicroglia. The sum of the spot intensities per cell was quantified; data are shown as geometric mean ± SEM; n=3 independent experiments, and statistical significance was determined using one-way ANOVA and Tukey’s method for multiple comparisons. E and F) WT and LRRK2 G2019S iMicroglia were treated with vehicle or imiglucerase (1μM) for 72 hours, and the levels of GlcSph and BMP (22:6/22:6) were measured using LC-MS/MS. Data are shown as geometric mean ± SEM; n=3 independent experiments, and statistical significance was determined using two-way ANOVA and Sidak’s method for multiple comparisons. *P < 0.05, **P < 0.01, ****P < 0.0001.

### Defects in glucosylceramide catabolism correlate with BMP alterations in CSF from PD patients

To assess the relevance of GCase-dependent changes in BMP in human disease, we measured the levels of various GSLs and BMP in CSF from healthy control and PD subjects with or without LRRK2 variants using samples collected from the LRRK2 Cohort Consortium (SI Table 4)(47). BMP and GSL levels were comparable in healthy subjects that carry a LRRK2 variant compared to non-variant carrier controls (Supplementary Fig. S5). In contrast, significant lipid alterations were observed in CSF from LRRK2-PD patients compared to sporadic PD patients. BMP (22:6/22:6) levels were reduced by 36% (unadjusted p = 0.01) in CSF from PD patients that carry a LRRK2 variant relative to the PD patients without LRRK2 variants (Fig. 7A and B). Within the GSL lipid class, GlcCer (d18:1/24:1) increased by 15% (unadjusted p = 0.07), and LacCer (d18:1/24:1) decreased by 53%, (unadjusted p = 0.07) in LRRK2 variant PD subjects compared to PD patients that do not carry a variant in *LRRK2* (Fig. 7A and B, Supplementary Fig. S5). Significant alterations in BMP and GSLs were observed specifically in CSF from LRRK2 variant carriers with PD, suggesting elevated LRRK2 kinase activity drives BMP and GSL alterations in disease.

**Figure 7:**
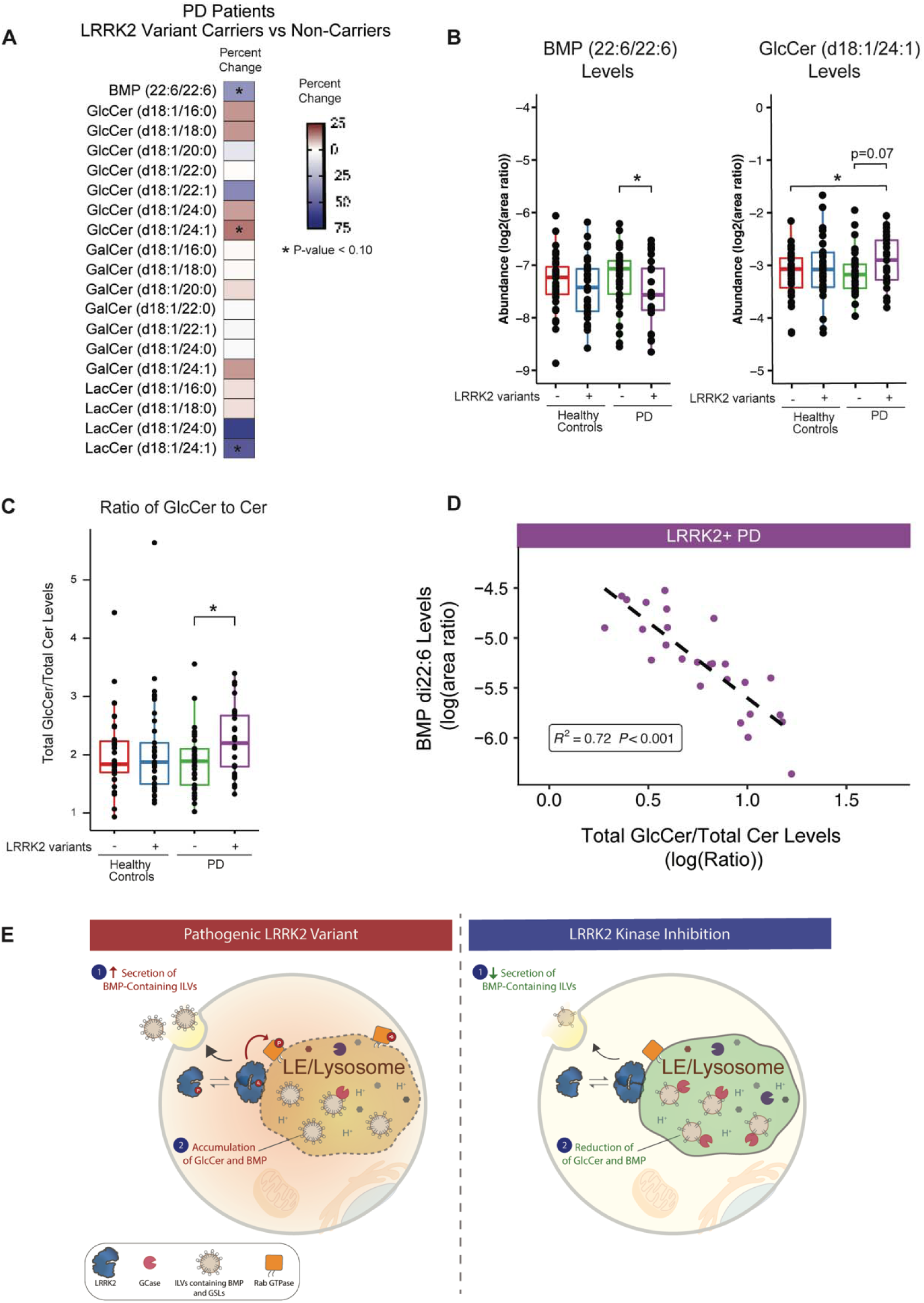
Lipidomic analysis of CSF shows alterations in GlcCer and BMP in PD patients with and without LRRK2 variants. Targeted analyses of CSF lipid levels were determined by LC-MS/MS in human subjects from the LCC cohort (Healthy controls without LRRK2 variants: N=35; Healthy controls with LRRK2 variants: N=37; PD patients without LRRK2 variants: N=37; PD patients with LRRK2 variants: N=26. A) Heatmap showing % change in lipid abundance detected in CSF from PD patients with LRRK2 variant carriers compared to non-carriers. % changes and significance of effects were analyzed using robust linear model with sex and age as covariates. * unadjusted p <0.10. B) Relative abundance of BMP (22:6/22:6) and GlcCer (d18:1/24:1) levels were measured in CSF. Significance of change was analyzed by linear model with pairwise comparisons by Tukey’s honest significant difference test with significance set at unadjusted p value of 0.05. Main box and error bars depict interquartile ranges of top 75^th^ or bottom 25^th^ percentile and largest and smallest value with 1.5 times the interquartile ranges above and below 75^th^ or 25^th^ percentiles. Median 50^th^ percentile is shown as midline within each boxplot. *p < 0.05. C) The ratio of total GlcCer to total ceramide (Cer) levels were measured across study participants. Total GlcCer was calculated as sum of area ratios from GlcCer (d18:1/16:0), GlcCer (d18:1/18:0), GlcCer (d18:1/24:0), and GlcCer (d18:1/24:1). Total Cer was calculated as sum of area ratios from Cer lipids with identically matched acyl chain groups as GlcCer quantified. Significance of change was analyzed by linear model with pairwise comparisons by Tukey’s honest significant difference test. Ranges of values illustrated are the same as B; *p < 0.05. D) Pearson correlation coefficient and significance of correlations between BMP (22:6/22:6) and GlcCer (d18:1/24:1) in LRRK2 variant carriers with PD. (E) Model for mechanisms by which LRRK2 regulates BMP and glycosphingolipid levels. Pathogenic variants in LRRK2 lead to an increased phosphorylation of relevant Rabs at the lysosome, impaired endolysosomal GCase activity, and reduced lysosomal proteolysis. We propose two potential mechanisms that are employed depending on the tissue examined in response to LRRK2-mediated lysosomal dysfunction: (1) in kidney, we propose that LRRK2 hyperactivity promotes increased secretion of BMP- and glycosphingolipid-containing vesicles as a compensatory response to help clear accumulated lipids and proteins and (2) in other cell types, including in the brain, increased LRRK2 activity leads to an increase in intracellular GCase substrate levels, and BMP is upregulated as a compensatory response to boost GCase activity. LRRK2 kinase inhibition corrects lysosomal dysfunction and obviates the need for compensatory measures to restore lysosomal homeostasis, leading to a reduction in BMP and GSL secretion and BMP and GCase substrate accumulation in cells.

Analysis of the substrate levels compared to the levels of product generated has been employed to approximate enzymatic activity and improve sensitivity for identifying genetic associations with lipid phenotypes(48). Therefore, we quantified the ratio of the sum of GlCer species to the sum of corresponding Cer species across healthy subjects and PD patients with or without LRRK2 variants. We observed a significant increase in the total GlcCer/total Cer ratio from 1.9 ± 0.5 (mean ± SD) in PD subjects without LRRK2 variants to 2.2 ± 0.6 (mean ± SD) in PD subjects with LRRK2 variants (unadjusted p = 0.02) (Fig. 7C), suggesting glucosylceramide catabolism is reduced in PD patients that carry a LRRK2 variant. Given our findings that LRRK2 regulates BMP as a secondary response to its effects on GCase activity, we hypothesized that elevations in the ratio of total GlcCer to Cer might correlate with alterations in BMP in CSF from LRRK2-PD patients. We found a strong inverse correlation (r^2^ = 0.72, p < 0.001, Pearson correlation) between the levels of BMP (22:6/22:6) and the ratio of GlcCer to Cer levels in CSF from PD subjects that carry a LRRK2 variant. Together, our results demonstrate that alterations in BMP correlate with reduced glucosylceramide catabolism in CSF from PD patients that carry a variant in LRRK2 and suggest that LRRK2-dependent effects on GCase activity may drive BMP alterations in disease.

## Discussion

Variants in LRRK2 associated with increased kinase activity are proposed to contribute to PD pathogenesis, and growing evidence suggests LRRK2 activity is similarly elevated in patients with sporadic disease, suggesting this may represent a central node of dysfunction in PD. While kinase-activating LRRK2 variants have been shown to perturb lysosomal function(49, 50), many questions remain about how LRRK2 modulates the endolysosomal system and whether such regulation is relevant in PD. Our study reveals that LRRK2 regulates the lysosomal pathway by controlling the levels of BMP, a lysosomal lipid important for lipase activities, and GSLs, including GCase substrates that accumulate in PD. We establish the functional relevance of LRRK2-mediated effects on lysosomal homeostasis by demonstrating that LRRK2 regulates BMP and GSL levels in the brain using preclinical models. We further showed that such LRRK2-associated lipid dysregulation also occurs in CSF from PD patients. Finally, LRRK2 kinase inhibition fully normalized BMP and GSL levels in lysosomes and restored lysosomal proteolysis, highlighting the potential of LRRK2 kinase inhibition as a therapeutic approach to correct lysosomal dysfunction in PD.

Data from preclinical and recent Ph1 and Ph1b studies show that LRRK2 kinase inhibition reduces levels of urine BMP(29, 30), but the implications of such changes with respect to disease or lysosomal function in tissue were poorly defined. Here, we confirmed that LRRK2 G2019S carriers have increased BMP (22:6/22:6) levels in urine(31) and established that carriers of the LRRK2 protective variant (N551K) have decreased urinary BMP levels, mirroring the effects observed with LRRK2 kinase inhibition. As carriers of the protective LRRK2 variant have been shown to have reduced LRRK2 activity, these data provide genetic support that decreased kinase activity is associated with lower urine BMP levels and add to growing data that suggest LRRK2 kinase inhibition recapitulates functional changes observed with a variant that reduces PD risk. BMP and glycosphingolipids are enriched in intraluminal vesicles within multivesicular bodies (MVBs) and can be secreted into the extracellular space when MVBs fuse with the plasma membrane in the form of exosomes(34). Our data demonstrate that LRRK2 deletion or inhibition leads to the intracellular accumulation of BMP and glycosphingolipids in mouse kidney and their corresponding depletion from urine, suggesting LRRK2 may serve to regulate the secretion or content of exosomes. The mechanism by which LRRK2 controls exosome secretion and/or content remains unclear, but a tantalizing possibility is that this may occur through LRRK2’s known kinase activity on specific Rab GTPases. Indeed, several Rab GTPases, including the LRRK2 substrate Rab35, have been shown to directly impact the biogenesis or secretion of exosomes. For these reasons, future studies are warranted to determine whether LRRK2-dependent Rab phosphorylation mediates the effects of LRRK2 activity on exosome release(51, 52). Emerging data suggest that exosome secretion may be employed as a homeostatic response to lysosomal damage to aid in the removal of lysosomal proteins and lipids that are not effectively degraded(33, 53, 54). Accordingly, LRRK2 may normally function to upregulate exosome secretion as a compensatory action to counter defects in lysosomal proteolysis and lipid catabolism, either directly caused by pathogenic variants in LRRK2 or caused by other lysosomal proteins perturbed in PD (Fig. 7E).

Urinary BMP has been effectively used as a biomarker to assess LRRK2-dependent lysosomal changes in the periphery, but the impact of LRRK2 on lysosomal function and lipid homeostasis in the brain and potential relevance of BMP as a CNS biomarker of LRRK2 activity had not been explored. We found BMP levels were not grossly impacted by LRRK2 throughout the mouse brain but were modestly altered in astrocytes and in neurons upon aging in a LRRK2-dependent fashion. In contrast, we observed broad alterations in many classes of glycosphingolipids, including GalCer, GlcCer, and sulfatides, across CNS cells isolated from LRRK2 KO and G2019S mice. While the directionality of BMP changes in CNS cells from LRRK2 mouse models were consistent with those observed in kidney, glycosphingolipid levels were differentially impacted. Glial cells from LRRK2 G2019S mice showed accumulation of several glycosphingolipid species, including the GCase substrate GlcCer, but reduced levels of BMP. As BMP and the glycosphingolipids assessed are enriched in intraluminal vesicles and secreted exosomes, these data suggest that LRRK2 activity affects glycosphingolipid levels in the brain through an additional mechanism beyond its effects on exocytosis. Consistent with data from our LRRK2 mouse models, we also observed reduced levels of BMP and elevated levels of GlcCer in CSF from Parkinson’s disease patients that carry a LRRK2 variant, demonstrating the relevance of these LRRK2-dependent lipid changes in disease. Our data are consistent with previous work showing reductions in lysosomal hydrolase activity and GlcCer accumulation in the substantia nigra from PD patients, and provide additional evidence that defects in lysosomal function and glycosphingolipid metabolism, specifically, may play a key role in PD pathogenesis(55). These results also highlight the potential utility of glycosphingolipids as lysosomal biomarkers of LRRK2 activity in the brain and suggest LRRK2 may more subtly impact BMP levels in CSF.

Our data demonstrate that LRRK2 exerts its effects on glycosphingolipid levels by modulating GCase activity within lysosomes, and we propose that reduced GCase activity may be a primary driver of LRRK2-dependent lysosomal dysfunction. Using a recently-described activity-based probe to measure lysosomal GCase activity, we observed an approximately 50% decrease in lysosomal GCase activity in LRRK2 variant cell models, including iMicroglia(44). We established that this reduction in GCase activity correlated with accumulation of GCase substrates and BMP in lysosomes and with defects in lysosomal proteolysis in cellular models expressing disease-associated LRRK2 variants. Importantly, LRRK2 inhibition normalizes lysosomal dysfunction caused mediated by LRRK2 activity as DNL151 treatment fully corrected lipid accumulation and lysosomal proteolysis in pathogenic LRRK2 cellular models. Previous work showed that LRRK2 inhibition can also correct lysosomal dysfunction in GCase loss-of-function cellular models, and our data suggest that the mechanism by which LRRK2 inhibition may rescue GCase-dependent defects is through LRRK2’s effects on GCase substrate levels(30, 43, 56, 57). Further, we propose that LRRK2 activity regulates BMP levels in lysosomes as a downstream consequence of LRRK2-dependent changes in endolysosomal GCase activity as treatment with exogenous GCase was sufficient to reduce BMP accumulation in LRRK2 variant A549 cells and iPSC-derived microglia. There is precedent for alterations in BMP levels or composition resulting from deficient enzyme activity and primary storage across many LSDs, including in Gaucher disease(58-62). BMP is thought to accumulate either as a secondary effect of broader impairments in lysosomal function and lipid catabolism or as an adaptive response to help degrade accumulated glycosphingolipids(59). Our data from CNS cells isolated from LRRK2 G2019S mice add support to this hypothesis given glycosphingolipid dysregulation was evident as early as 5-6 months of age and BMP alterations only occurred later on in more aged mice. Additional studies are needed to understand whether LRRK2 affects BMP synthesis or catabolism in response to reduced GCase activity and uncover why polyunsaturated fatty acid species of BMP are particularly impacted. Our analysis of BMP and glycosphingolipid levels in kidney and urine from LRRK2 mouse models suggests that LRRK2 can also regulate BMP by controlling its secretion, suggesting that LRRK2 can employ multiple mechanisms to regulate lipid homeostasis and that the directionality of LRRK2’s effects on BMP may be dictated by which mechanism is favored in specific cell types (Fig. 7E).

Our results demonstrate that inhibition of LRRK2 kinase activity can normalize GCase substrate accumulation and downstream deficits in lysosomal function and highlight the therapeutic potential of LRRK2 inhibitors to correct lysosomal dysfunction observed in PD. Preclinical studies have shown that LRRK2 kinase inhibition can rescue lysosomal dysfunction not directly driven by kinase-activating mutations in LRRK2, and our work provides additional support suggesting LRRK2-dependent changes in lysosomal function may occur more broadly in PD patients beyond those that carry a variant in LRRK2(30, 43, 57, 63). Variants in *GBA1*are observed in approximately 10% of PD patients, and increasing data, including that reported in the present work, suggest that GCase activity is impaired in PD patients beyond those that carry deleterious *GBA1* variants(64-68). Our work demonstrates that treatment with DNL151 can rescue lysosomal defects stemming from reduced GCase activity and provides additional preclinical support for the idea that LRRK2 inhibition may normalize lysosomal dysfunction more broadly in PD and could provide benefit to others beyond pathogenic LRRK2 variant carriers. Interestingly, human subjects that carry both the LRRK2 G2019S variant and an additional variant in GBA1 do not have a worse clinical course of PD than those that carry a variant in either gene and, in fact, may have a beneficial effect with respect to cognitive decline(69). DNL151 is currently being tested in late-stage clinical studies in both LRRK2-PD and sporadic PD patients, and the question of whether LRRK2 kinase inhibition can rescue PD-relevant defects in lysosomal homeostasis and modify disease progression in these patient populations will ultimately be resolved in the clinic.

## Materials and Methods

### Genotype analysis of BMP levels in urine from human subjects

We used publicly available data from PPMI to explore the relationship between PD risk and protective variants in LRRK2 and urine BMP levels. For the genetic data, we used WGS data on PPMI subjects available through the Accelerating Medicines Partnership – Parkinson’s Disease (AMP-PD) data portal. Quality control and processing of these data has been previously described(70). A total of 10,418 samples were included in the AMP-PD WGS data (data release 2021_v2-5release_0510), of which 1,807 were from the PPMI study. To predict ancestry of these samples, we merged the data with 1000 Genomes Phase 3 (1000G) reference samples(71). After linkage disequilibrium pruning this merged dataset, we performed principal component analysis (PCA) and then used the first 10 principal components and 1000G super population labels to predict genetic ancestry for all samples in our dataset via the k-nearest neighbors algorithm. Because a vast majority (N = 1,741) of PPMI samples were predicted to be of European ancestry, we elected to restrict our downstream analyses to just this group.

Urine BMP measurements were available in PPMI for the following species of BMP: total di-BMP-22:6, 2,2’di-22:6, and total di-BMP-18:1. These measurements, along with those for urinary creatinine, were generated under PPMI Project ID #145; full details on measurement methodology and data processing can be found in the methods document for this project available through PPMI. After restricting to measurements taken at the PPMI baseline visit, BMP measurements (reported in ng/mg creatinine) were available on 1,232 subjects. Of these, 1,133 samples overlapped with the 1,741 predicted European ancestry samples with available WGS data. Finally, we removed 69 samples with a case/control designation of “Other” in the available AMP-PD metadata, a classification which included prodromal subjects and those with other neurological disorders. Therefore, a final cohort of 1,069 subjects (classified as having PD or as controls only) with joint WGS and urine BMP measurements were used for SNP association testing.

### Animal Care

All procedures in animals were performed with adherence to ethical regulations and protocols approved by Denali Therapeutics Institutional Animal Care and Use Committee. Mice were housed under a 12-hour light/dark cycle and had access to water and standard rodent diet (#25502, irradiated; LabDiet) ad libitum. In-diet dosing studies utilized compound-specific rodent chow outlined below.

### Mouse strains and subject details

LRRK2-KO (C57BL/6-Lrrk2tm1.1Mjff/J) and LRRK2 G2019S KI mice (C57BL/6-Lrrk2tm4.1Arte) mice were obtained from The Jackson Laboratory (Strain #016121) and Taconic Biosciences Inc (Model# 13940), respectively. Genetically modified mice and their wild-type littermates were age- and sex-matched across groups for experiments. 4-6 month old LRRK2 KO and WT mice were used to analyze BMP and glycosphingolipid concentrations in urine (Figure 1B, Figure 2A), kidney (Figure 1D, 2B-C) and FACS-sorted brain cells (Figure 3B-D). LRRK2 G2019S KI and WT mice were sacrificed at 5-6 months of age (young) and 18 months of age (old) to evaluate BMP and glycosphingolipid concentrations in FACS-sorted brain cells (Figure 3E-3I). 6-8 month old LRRK2 G2019S KI and WT mice, treated with vehicle or MLi-2 (LRRK2 inhibitor) diet for 35 days, were used to analyze BMP and glycosphingolipid concentrations in kidney (Figure 2D, F, G) and urine (Figure 2E).

### LRRK2 inhibitor treatment in LRRK2 G2019S KI mice

MLi-2 (MedChemExpress, Monmouth Junction, NJ) was used as a tool LRRK2 inhibitor for in vivo experiments. Grain-based, bacon-flavored rodent chow was formulated into pellets containing MLi-2 (960 mg/kg diet, estimated to be 100 mg/kg per mice) and irradiated for in-diet dosing (Bio-Serv; LabDiet; PicoLab Mouse Diet 20, 5058). Mice were provided with vehicle or MLi-2 diet ad libitum for 35 days. Food weight and mouse body weight were routinely monitored to evaluate diet consumption and animal health.

### Biofluid and tissue collection from LRRK2 mouse models

For studies without MLi-2 treatment, urine was collected over 3 consecutive days to capture animals that did not urinate in initial attempts. For MLi-2 treatment studies, urine was collected on the day of terminal tissue collection. Urine samples were snap-frozen on dry ice and transferred to −80 °C for storage. For terminal tissue collection, mice were anesthetized with 2.5% tribromoethanol. Once deeply anesthetized, animals were transcardially perfused with ice-cold PBS using a peristaltic pump for a minimum of 3 min at a rate of 5 ml/min. After perfusion, brain was collected for subsequent FACS analysis (see section below). Kidney was removed and subdissected into the renal cortex and medulla. Each portion was weighed (20 ± 2 mg), collected in 1.5-ml Eppendorf tubes, frozen on dry ice, and stored at −80 °C. Urine, renal medulla and renal cortex samples were prepared for lipid extraction, and extracted BMP and glycosphingolipid species were quantified.

### MALDI-imaging mass spectrometry analysis of BMP levels in kidney and brain of LRRK2 KO mice

#### Sample preparation

LRRK2 KO mice and their wild-type littermates (5 to 6 months old) were used for MALDI-IMS experiments. Following transcardial perfusion with ice-cold PBS, brain and kidney tissues were collected and flash frozen on aluminum foil that was slowly lowered into liquid nitrogen for approximately 10 seconds. Frozen tissue was stored at −80°C until ready for use. Prior to sectioning, the tissues were removed from the −80°C freezer and placed in the cryostat chamber to equilibrate to −20 °C. Brain and kidney tissues were cut on a cryostat (Leica Biosystems) into 12 μm thick sections and thaw-mounted onto indium-tix oxide (ITO) coated glass slides (Delta Technologies). Additional sections were obtained for H&E staining. After staining, digital micrographs were obtained via a slide scanner (Leica Biosystems).

#### Matrix application

The ITO-coated slides were coated with 1,5-diaminonaphthalene (DAN) MALDI matrix via sublimation (86, 87). For kidney, a custom glass sublimation apparatus was used. Briefly, 100 mg of recrystallized DAN was placed in the bottom of a glass sublimation apparatus (Chemglass Life Sciences). The apparatus was placed on a metal heating block set to 130 °C and DAN was sublimated onto the tissue surface for 4 minutes at a pressure of less than 25 mTorr.

Approximately 2.8 mg of DAN was applied to the ITO slide, determined by weighing the slide before and after matrix application. A home-built sublimation device was used for the brain sections. Briefly, approximately 19 mg of recrystallized DAN was dissolved in 2 ml of acetone and deposited on the bottom of the device. Heat was applied for 10 minutes at >130 °C and all DAN was sublimated. The coated plates were then subjected to mass spectrometer analysis.

#### Imaging mass spectrometry

The tissue sections were imaged on a Solarix 15T FT-ICR MS (Bruker Daltonics), equipped with a SmartBeam II 2 kHz frequency tripled Nd:YAG laser (355 nm). Images were acquired at 125 m (brain) step size in negative ion mode. Each pixel is the average of 1500 laser shots using the small laser focus setting and random-walking within the pixel. The mass spectrometer was externally calibrated with a series of red phosphorus clusters. Data were collected from m/z 345 – 1,000. Images were generated using FlexImaging 3.0 (Bruker Daltonics). BMP was identified by accurate mass, with the mass accuracies typically better than 1 ppm. Brain regions of interest (ROIs) were selected corresponding to the cortex, midbrain, hippocampus, and striatum (Supplemental Fig. 2B). Average spectra for the ROIs were exported from FlexImaging as .csv files and imported into mMass(72, 73). Spectra for all four ROIs were overlaid to directly compare peak intensities. The resulting graphs for m/z 865.502 (corresponding to BMP 22:6/22:6) and m/z 834.530 (corresponding to PS 40:6) are shown in Supplemental Figure 2C and 2D.

### FACS-Based Analysis of BMP and glycosphingolipid levels in mouse CNS cells

Mice were perfused with PBS, and whole brains dissected, and processed into a single cell suspension according to the manufacturers’ protocol using the adult brain dissociation kit (Miltenyi Biotec 130-107-677). Cells were Fc blocked (Biolegend #101320, 1:100) and stained for flow cytometric analysis with Fixable Viability Stain BV510 (BD Biosciences #564406, 1:100) to exclude dead cells, CD11b-BV421 (BD Biosciences 562605, 1:100), CD31-PerCP Cy5.5 (BD Biosciences #562861, 1:100), O1-488 (Thermo/eBio #14-6506-82, 1:50), Thy1-PE (R&D #FAB7335P, 1:100), and EAAT2-633 (Alomone #AGC-022-FR, 1:60). Cells were washed with PBS/1% BSA and strained through a 100μ microglia, EAAT2+ astrocytes, and Thy1+ neurons on a FACS Aria III (BD Biosciences) with a 100 μm nozzle. In order to achieve pure populations of astrocytes, microglia, and neurons negative gates were set to remove O1+ and CD31+ cells which are predominantly oligodendrocytes and endothelial cells respectively. The whole procedures including single cell suspension, staining, and sorting were kept at 4C. Sorted cells were directly collected into methanol extraction buffer for LC-MS/MS analysis (described below).

### LC-MS Based Analysis of Lipids

#### Sample Preparation

##### FACS-sorted brain cells

100,000 cells were sorted directly into tubes containing 800 µl of LC-MS grade methanol containing 2 µl of internal standard mix. These samples were vortex for 5 min, and centrifuged at 21,000 x g for 10 min at 4 °C. The supernatants were transferred to a new glass vial, and half of the sample was aliquoted for GlcCer/GalCer analysis, while half of the sample was used for other lipid panel analysis. The samples were dried down under a constant stream of N_2_, and stored at −80 °C until analysis.

##### Kidney and brain tissues

Approximately 20 mg of kidney samples were placed in Safe-Lock Eppendorf tube (Eppendorf Cat#022600044) containing 5 mm stainless steel beads (QIAGEN Cat#69989). To these tubes, 400 µl of LC-MS grade methanol containing 2 µl internal standard mix solution was added and homogenized for 30 sec at 25 HZ at 4 °C with Tissuelyser.

Homogenized samples were centrifuged at 21,000 x g for 20 min at 4 °C. Cleared methanolic supernatants were transferred into new Eppendorf vials and kept in −20 °C for 1 hour. Samples were centrifuged at 21,000 g for 10 min at 4 °C and supernatants were aliquoted into 96 well plates with glass inserts and dried under a constant stream of N_2_. Dried samples were stored in −80 °C until analysis.

##### Urine exosomes

Urine was collected into a container pre dose or 28 days post dose. Within 30 minutes of the end of the collection period, samples were centrifuged at 2500g for 15 minutes at 4 °C. Urine was transferred to aliquot tubes and stored at −80 °C. Samples were thawed on ice and centrifuged at 1000g for 10 minutes at 4°C to remove particulates. 500 μL urine was transferred for exosome isolation according to the EVTRAP method previously described(74). Following elution, samples were processed for LC-MS analysis. Single step extraction of pelleted exosomes were performed by adding 400 µl of LC-MS grade methanol containing 2 µl internal standard mix solution. Resultant supernatants were transferred into 96 well plates with glass inserts and dried under a constant stream of N2. Dried samples were stored in −80 °C until analysis.

##### Genotype and drug-treatment analysis in cell lysates

A549 cells were seeded into standard 6-well cell culture plates at a density of 90K cells/well. For iMicroglia, cells were seeded at 20K cells/well into CellCarrier-96 Ultra Microplates (Perkin Elmer #6055302). At the time of harvest, cell media was aspirated and cells were washed with ice-cold 0.9% NaCl “Normal Saline”. Cells were then quickly extracted into 400μL per well of ice-cold extraction buffer (9:1; LCMS-grade MeOH:H2O) plus 2μL of internal standards. Cell extracts were scrapped off the well then transferred into a clean Eppendorf tube. Extracts were then shaken at 2000 r.p.m. for 20 minutes at 4 °C, then centrifuged at 21,000 x g at 4 °C for 5 minutes to pellet insoluble material. Next, 100μL of the cleared methanolic supernatant was transferred into glass vials or plates with glass inserts. These samples were dried under a stream of N2 gas for 4 hours and stored in −80C until analysis. To assess the effect of recombinant GCase addition to lipid levels in cells, cells were supplemented with imiglucerase (Sanofi) at a final concentration of 2μM in water. A549 cells were treated with imiglucerase for 96 hours with full replacement/exchange of medium at 48 hours. For iMicroglia plated in NGD+ media, cells were treated with 1μM imiglucerase for 72h. Cells were then processed for downstream analysis as described above.

##### Exosomes from cell culture media

WT or LRRK2 G2019S iMG were grown in NGD+ media for 72h. Supernatant media was collected and centrifuged at 2500g for 10 min to cell debris. Supernatant was then transferred for exosome isolation according to the EVTRAP method previously described(74). Following elution, samples were analyzed by mass spectrometry.

##### Urine and CSF sample extraction

Samples were centrifuged at 1000 x g at 4 °C to remove particulates. Twenty µl of urine and 10 µl of CSF samples were transferred into a 2 ml Safe-lock Eppendorf tubes. To these samples, 200 µl of LC-MS grade methanol containing 2 µl of internal standard mix were added and vortexed for 5 min. Samples were centrifuged at 21,000 x g for 4 °C and stored at 20 C for 1 hr. Following this step, samples were centrifuged once again at 21000 x g for 20 min at 4 °C. Resultant supernatants were transferred to 96 well plates with glass inserts and dried under constant stream of N_2_. Dried samples were stored at −80 °C until analysis.

### Sample preparation for glucosylceramide and galactosylceramide measurements

For analysis and separation of glucosylceramides and galactosylceramides, 50 µl of above extracts were dried under constant stream of N_2_ for 4 hours and resuspended in 200 µl of 92.5/5/2.5 LCMS grade acetonitrile/isopropanol/water fortified with 5 mM ammonium formate and 0.5% formic acid.

### LCMS targeted analysis of lipids

Lipid analyses were performed by liquid chromatography (UHPLC Nexera X2, UHPLC ExionLC) coupled to electrospray mass spectrometry (QTRAP 6500+). For each analysis, 5 μL of sample was injected on a BEH C18 1.7 μm, 2.1×100 mm column (Waters) using a flow rate of 0.25 mL/min at 55°C. For positive ionization mode, mobile phase A consisted of 60:40 acetonitrile/water (v/v) with 10 mM ammonium formate + 0.1% formic acid; mobile phase B consisted of 90:10 isopropyl alcohol/acetonitrile (v/v) with 10 mM ammonium formate + 0.1% formic acid. For negative ionization mode, mobile phase A consisted of 60:40 acetonitrile/water (v/v) with 10 mM ammonium acetate + 0.1% acetic acid; mobile phase B consisted of 90:10 isopropyl alcohol/acetonitrile (v/v) with 10 mM ammonium acetate + 0.1% acetic acid. The gradient was programmed as follows: 0.0-8.0 min from 45% B to 99% B, 8.0-9.0 min at 99% B, 9.0-9.1 min to45% B, and 9.1-10.0 min at 45% B. Electrospray ionization was performed in positive or negative ion mode. For the QTRAP 6500+, we applied the following settings: curtain gas at 30 psi (negative mode) and curtain gas at 40 psi (positive mode); collision gas was set at medium; ion spray voltage at 5500 V (positive mode) or −4500 V (negative mode); temperature at 250 °C (positive mode) or 600 °C (negative mode); ion source Gas 1 at 55 psi; ion source Gas 2 at 60 psi; entrance potential at 10 V (positive mode) or −10 V (negative mode); and collision cell exit potential at 12.5 V (positive mode) or −15.0 V (negative mode). Data acquisition was performed in multiple reaction monitoring mode (MRM) with the collision energy (CE) values reported in Tables 1 and 2. Lipids were quantified using a mixture of non-endogenous internal standards as reported in Table 1 and 2. Quantification was performed using MultiQuant 3.02.

### LCMS targeted analysis of GlcCer and GalCer

Glucosylceramide, galactosylceramide and glucosylsphingosine analyses were performed by liquid chromatography (UHPLC Nexera X2, UHPLC ExionLC) coupled to electrospray mass spectrometry (TQ 6495C). For each analysis, 5 μL of sample was injected on a HALO HILIC 2.0 μm, 3.0 × 150 mm column (Advanced Materials Technology, PN 91813-701) using a flow rate of 0.48mL/min at 45 °C. Mobile phase A consisted of 92.5/5/2.5 ACN/IPA/H2O with 5 mM ammonium formate and 0.5% formic acid. Mobile phase B consisted of 92.5/5/2.5 H2O/IPA/ACN with 5 mM ammonium formate and 0.5% formic acid. The gradient was programmed as follows: 0.0–2 min at 100% B, 2.1 min at 95% B, 4.5 min at 85% B, hold to 6.0 min at 85% B, drop to 0% B at 6.1 min and hold to 7.0 min, ramp back to 100% at 7.1 min and hold to 8.5 min. Electrospray ionization was performed in positive mode. Agilent TQ 6495C was operated with the following settings: gas temp at 180 °C; gas flow 17 L/min; nebulizer 35 psi; sheath gas temp 350°C; sheath gas flow 10 L/min; capillary 3500 V; nozzle voltage 500 V. Glucosylceramide and galactosylceramide species were identified based on their retention times and MRM properties of commercially available reference standards (Avanti Polar Lipids, Birmingham, AL, USA). Quantification was performed using Skyline (v19.1; University of Washington). Table 3 shows specific analytes and internal standards used in this assay.

**Table 3:** GalCer/GlcCer acquisition parameters

| **Name** | **Internal Standard** | **Q1 m/z** | **Q3 m/z** | **CE (V)** |
| --- | --- | --- | --- | --- |
| alpha-GalCer(d18:1/16:0) | GalCer(d18:1/15:0) | 700.6 | 264.3 | 32 |
| alpha-GalCer(d18:1/18:0) | GalCer(d18:1/15:0) | 728.6 | 264.3 | 32 |
| alpha-GalCer(d18:1/20:0) | GalCer(d18:1/15:0) | 756.6 | 264.3 | 32 |
| alpha-GalCer(d18:1/22:0) | GalCer(d18:1/15:0) | 784.7 | 264.3 | 32 |
| alpha-GalCer(d18:1/22:1) | GalCer(d18:1/15:0) | 782.7 | 264.3 | 32 |
| alpha-GalCer(d18:1/24:0) | GalCer(d18:1/15:0) | 812.7 | 264.3 | 32 |
| alpha-GalCer(d18:1/24:1) | GalCer(d18:1/15:0) | 810.7 | 264.3 | 32 |
| alpha-GalCer(d18:2/18:0) | GalCer(d18:1/15:0) | 726.6 | 262.3 | 32 |
| alpha-GalCer(d18:2/20:0) | GalCer(d18:1/15:0) | 754.6 | 262.3 | 32 |
| alpha-GalCer(d18:2/22:0) | GalCer(d18:1/15:0) | 782.7 | 262.3 | 32 |
| Cholesteryl galactoside | GlcCer(d18:1(d5)/18:0) | 566.6 | 369.3 | 13 |
| Cholesteryl glucoside | GlcCer(d18:1(d5)/18:0) | 566.6 | 369.3 | 13 |
| Galactosylsphingosine | Glucosylsphingosine(d5) | 462.2 | 282.3 | 18 |
| GalCer(d18:1/15:0) | IS | 686.3 | 264.3 | 32 |
| GalCer(d18:1/16:0) | GalCer(d18:1/15:0) | 700.6 | 264.3 | 32 |
| GalCer(d18:1/18:0) | GalCer(d18:1/15:0) | 728.6 | 264.3 | 32 |
| GalCer(d18:1/20:0) | GalCer(d18:1/15:0) | 756.6 | 264.3 | 32 |
| GalCer(d18:1/22:0) | GalCer(d18:1/15:0) | 784.7 | 264.3 | 32 |
| GalCer(d18:1/22:1) | GalCer(d18:1/15:0) | 782.7 | 264.3 | 32 |
| GalCer(d18:1/24:0) | GalCer(d18:1/15:0) | 812.7 | 264.3 | 32 |
| GalCer(d18:1/24:1) | GalCer(d18:1/15:0) | 810.7 | 264.3 | 32 |
| GalCer(d18:2/18:0) | GalCer(d18:1/15:0) | 726.6 | 262.3 | 32 |
| GalCer(d18:2/20:0) | GalCer(d18:1/15:0) | 754.6 | 262.3 | 32 |
| GalCer(d18:2/22:0) | GalCer(d18:1/15:0) | 782.7 | 262.3 | 32 |
| GlcCer(d18:1(d5)/18:0) | IS | 733.6 | 269.3 | 32 |
| GlcCer(d18:1/16:0(d3)) | IS | 703.6 | 264.3 | 32 |
| GlcCer(d18:1/16:0) | GlcCer(d18:1/16:0(d3)) | 700.6 | 264.3 | 32 |
| GlcCer(d18:1/18:0) | GlcCer(d18:1(d5)/18:0) | 728.6 | 264.3 | 32 |
| GlcCer(d18:1/20:0) | GlcCer(d18:1(d5)/18:0) | 756.6 | 264.3 | 32 |
| GlcCer(d18:1/22:0) | GlcCer(d18:1(d5)/18:0) | 784.7 | 264.3 | 32 |
| GlcCer(d18:1/22:1) | GlcCer(d18:1(d5)/18:0) | 782.7 | 264.3 | 32 |
| GlcCer(d18:1/24:0) | GlcCer(d18:1(d5)/18:0) | 812.7 | 264.3 | 32 |
| GlcCer(d18:1/24:1) | GlcCer(d18:1(d5)/18:0) | 810.7 | 264.3 | 32 |
| GlcCer(d18:2/18:0) | GlcCer(d18:1(d5)/18:0) | 726.6 | 262.3 | 32 |
| GlcCer(d18:2/20:0) | GlcCer(d18:1(d5)/18:0) | 754.6 | 262.3 | 32 |
| GlcCer(d18:2/22:0) | GlcCer(d18:1(d5)/18:0) | 782.7 | 262.3 | 32 |
| Glucosylsphingosine | Glucosylsphingosine(d5) | 462.2 | 282.3 | 18 |
| Glucosylsphingosine(d5) | IS | 467.2 | 287.3 | 18 |
| Sitosteryl galactoside | GlcCer(d18:1(d5)/18:0) | 594.6 | 397.4 | 13 |
| Sitosteryl glucoside | GlcCer(d18:1(d5)/18:0) | 594.6 | 397.4 | 13 |

Note: IS - Internal standard.

### Cell Line Generation

Cell line engineering of A549 cells to generate homozygous LRRK2 R1441G (CGC/GGC) knock-in, homozygous LRRK2 KO, and homozygous GBA1 knock-out was performed using CRISPR/Cas9. Human LRRK2 G2019S and LRRK2 Knock-out (KO) iPSC lines were generated in human iPSCs obtained from a female clone from Thermo Fisher (#A18945). Sequence information for generating targeting gRNA, ssODN donor and PCR primers are as follows:

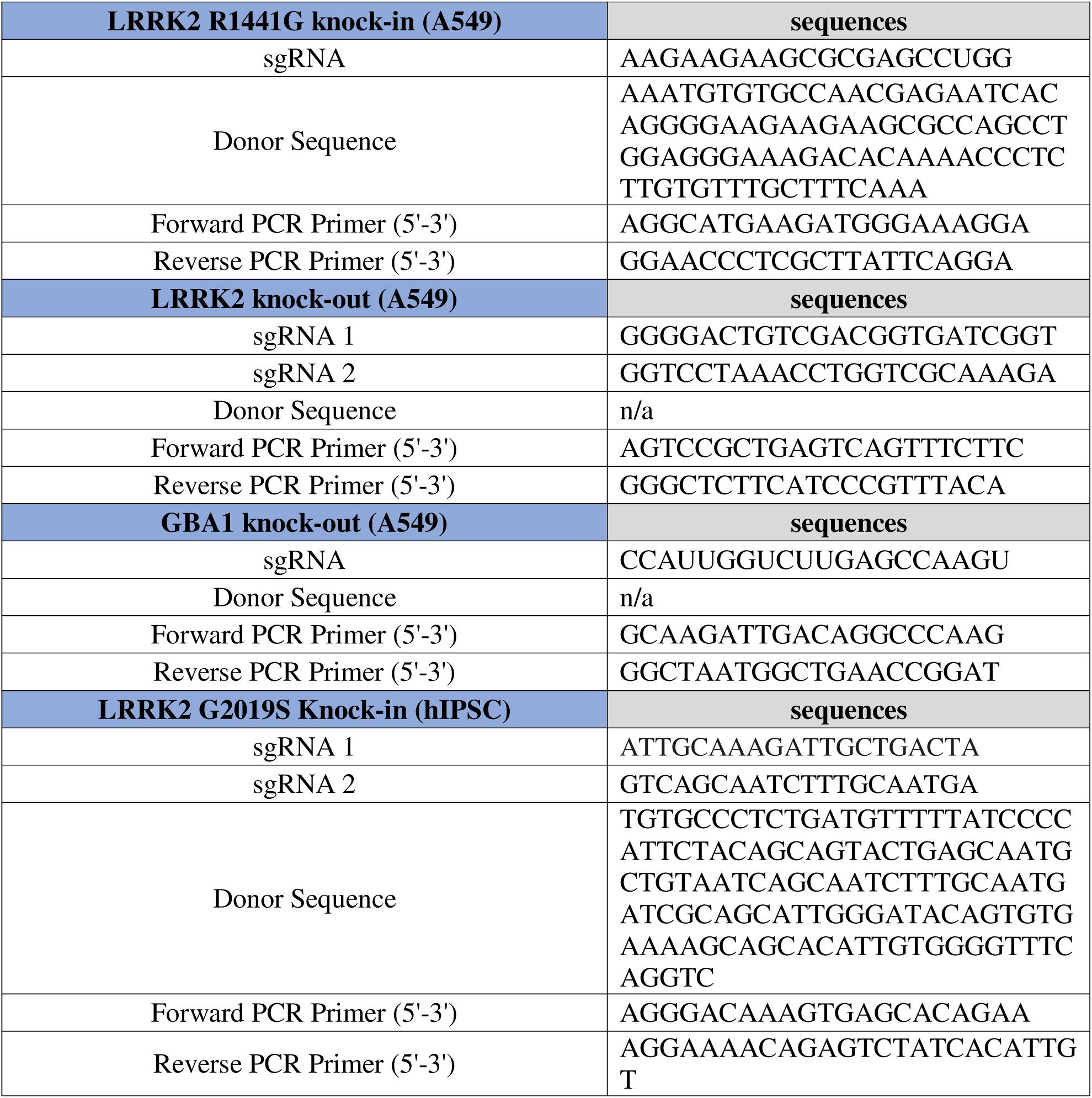
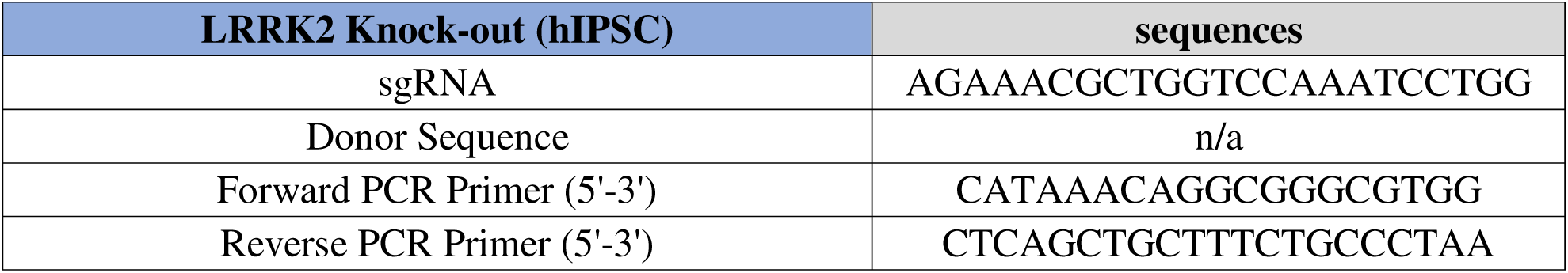

Human LRRK2 G2019S and LRRK2 Knock-out (KO) iPSC lines as described in table above were generated by using a nucleofection-based RNP approach to introduce Cas9 (NEB # M0646M) and sgRNAs against LRRK2 (crRNAs ordered from IDT) were introduced into iPSCs via nucleofection (Lonza P3 kit # V4XP-3032) as described before(75). sgRNAs were designed using the Broad Institute design tool based on a previous study(76). Clones were screened by T7 endonuclease and positive clones were further screened by TOPO cloning (Thermo cat #450030) to identify precise mutations. Clones with indels introduced into both alleles that result in a null mutation were grown up and used as LRRK2 KO clones.

To enable the rapid isolation of lysosomes using immunopurification, A549 cells (wild type, LRRK KO, LRRK2 R1441G and GBA1 KO) cells were transduced with lentivirus carrying the transgene cassette for expression of TMEM192-3x-HA. Stable expression cells were selected using resistance to Hygromycin B (Cat#10687010) supplied in growth medium at 200 g/mL for 21 days. Following selection cells were screened for the stable expression of TMEM192-3x-HA in lysosomes by quantifying the percentage of cells with co-localization of anti-HA and anti-LAMP1 by immunofluorescence, and by monitoring cell lysates for expression TMEM192-3x-HA (∼30 kDa) by western blot.

### Human iMicroglia Differentiation

Human iPSC-induced microglia were generated as previously described with minor modifications (75). Briefly, human iPSCs were routinely passaged as clumps onto Matrigel-coated plates with mTeSR+ media (StemCell Technologies #85850) according to manufacturer instructions. iPSCs were first differentiated into hematopoietic progenitor cells using a combined strategy of incorporating primitive hematopoiesis via a WNT switch(77)and sorting and replating CD235a+ cells in Medium B from a commercially available kit (StemCell Technologies #05310). Briefly: On Day 0 iPSCs were singularized with Accutase treatment for five minutes, pelleted at 300g for 5 mins, and seeded and cultured according to Guttikonda et al. On Day 3 cells were singularized using Accutase (∼10mins) and labeled with anti-CD235a-APC conjugated antibody and sorted using a BD FACS Aria. Cells were replated in prepared Medium B from Stemcell technologies. The manufacturer’s protocol was followed for an additional 7-10 days and numerous CD43+ progenitor cells (HPC) are collected and harvested during day 10-14. HPCs positive for the markers CD43 and negative for CD11b were transferred to a plate containing primary human astrocytes and co-cultured using media C adapted from a previous study(78). Once floating cells in co-culture were >90% mature microglia, cells were plated in fibronectin (Millipore FC010-10MG) - coated (5µg/mL) flat-bottom 48-well tissue culture plates (Corning #3548) for 3 days prior to experiments. Full characterization of human iPSC-derived microglia and additional details on the differentiation protocol have been published previously(75). iPSC-derived microglia were grown “C+++ media” composed of IMDM (GIBCO) media supplemented with 10% defined FBS (GIBCO), 1% Penicillin/Streptomycin (GIBCO), 20 ng/mL of hIL3 (Peprotech), 20 ng/mL of hGM-CSF (Peprotech) and 20 ng/mL of hM-CSF (Peprotech). For experiments, the previously-described C+++ media and “NGD+ media” adapted from Muffat et al. were used(79).

### Analysis of lysosomal proteolysis using DQ-BSA-based assay

A549 cells were seeded in 96 well poly-lysine coated plate, and then treated with DNL151 for 3 days. Cells were loaded with culture media containing 10 mg/ml of DQ Red BSA (Invitrogen D12051) and NucBlue (Thermo R37605) for 30 min, and then washed once and replaced with fresh culture media, followed by live cell imaging with 40X confocal (Opera Phenix high content imager). The image analysis was done with 9 hour time point after loading. Image analysis was performed using Harmony software. Spot analysis was used to identified DQ-BSA positive spots within the cell, with “corrected spot intensity” quantified. The Sum of corrected spot intensity per cell was used to measure DQ-BSA signals.

### Analysis of endo-lysosomal GCase activity using FQ6

LysoFQ-GBA was recently described and obtained from the laboratory of David Vocadlo (44). Briefly, cells were seeded into 96-well PDL-coated plates. FQ6 was added to cells for 1 hour under standard incubation conditions (humidified, 37C, 5% CO2) at a final concentration of 5μM for A549 cells and at 10μM for iMG using ‘C+++’ media. After 1 hour, cells were then rinsed 3x with 37°C wash buffer (live cell imaging solution (Invitrogen #A14291DJ) supplemented with 5.55 mM Glucose), counterstained with NucBlue (ThermoFisher) for staining nuclei for 10 min in imaging solution (wash buffer supplemented with 5% fetal bovine serum) and then imaged in the same buffer.

Imaging was performed on a Perkin Elmer Opera Phenix High Content Imaging System with acquisition of FQ6 (ex:488nm, em:500-550nm) and NucBlue(DAPI) (ex:375nm, em:435-480nm using a 40x water immersion objective. Analysis FQ6 signal “GCase activity” was performed using Harmony software. Harmony Spot Analysis was used to identify FQ6 positive spots within the cell, quantify the “corrected spot intensity”, and then the sum of corrected spot intensities per field of view was normalized to total number of nuclei in the field (to account for differences in cell number) and reported as “SUM Corrected Spot Intensity”.

### Lysosomal isolations

Lysosomes were isolated from A549 cells with stable expression of TMEM192-3x-HA. Rapid isolation of lysosomes was performed as described previously(39). Briefly, cells were seeded into 15cm tissue culture plates at a density of ∼5×10^6 cells/plate. After 72 hours under standard incubation (humidified, 37C, 5% CO2) cells were harvested. Medium was aspirated and cells rinsed 1x with ice-cold PBS. Cells were then scrapped into ice-cold KBPS (136 mM KCl, 10 mM KH_2_PO_4_, pH 7.25) and gently pelleted (300 x g, 5 min., 4C). Cell pellets were reconstituted into 500μM ice-cold KBPS+ (47mL KPBS, 3mL Opti-prep (Sigma D1556), and fresh protease inhibitor cocktail) and sheered by passage through a 21gauge needle. Post nuclear supernatants (PNS) were generated by centrifugation at 800 x g, 10 min., 4C. The PNS was transferred to a clean Eppendorf tube and volume adjusted to 1mL with KBPS. For rapid immunoprecipitation 80μL anti-HA magnetic resin (Thermo-Fisher #88836) is added and samples are rotated end over end at 4C for 15 minutes. Magnetic resin is captured onto a (? Magnet) and the captured lysosomes are washed 2x with 1mL KPBS+. Lysosomes are then processed immediately for downstream analysis.

### Analysis of association of LRRK2 variants with urine BMP levels in human subjects

A total of 1,069 samples were available for statistical testing of the association of LRRK2 variants on urine BMP levels (themselves available as ng/mg creatinine). Four BMP phenotypes were tested: the directly measured levels of total 22:6 and total 18:1 species, and derived ratios of 2,2’ 22:6/total 22:6, and total 22:6/total 18:1. To perform association testing, natural log-transformed BMP phenotypes were fit in a linear model consisting of age, sex, disease status, and the first 5 principal components derived from the WGS data (“Model 1”, see below). The residuals from this model were then inverse normal transformed using the blom() function from the “rcompanion” R package and used in association testing against the LRRK2 G2019S (rs34637584) and N551K (rs7308720) variants. Association testing was carried out using an additive model in plink v1.9(80). Finally, sensitivity analyses were conducted to examine the impact of removing the adjustment by disease status (“Model 2”), or adding adjustment by GBA N370S/N409S status (rs76763715; “Model 3”), LRRK2 N551K status (for G2019S inference only; “Model 4”) or LRRK2 G2019S status (for N551K inference only, “Model 5”). The choice to examine the effect of adjusting by LRRK2 G2019S and GBA N370S/N409S was informed by the large amount of carriers of these major PD risk-conferring variants, as they comprised 27% and 22% of the 1,069 samples, respectively.

### Analysis of CSF lipid levels from human subjects

The lipid analysis on human CSF reported in this manuscript was generated as part of a larger metabolomic characterization whose methods have been previously reported(47). Raw data from this study are available via online LCC data request at https://www.michaeljfox.org/news/lrrk2-cohort-consortium.

### CSF sample acquisition and human participants

CSF samples used in the analyses presented in this article were obtained from the MJFF-sponsored LRRK2 Cohort Consortium (LCC). For up-to-date information on the study, visit https://www.michaeljfox.org/biospecimens. The LRRK2 Cohort Consortium is coordinated and funded by The Michael J. Fox Foundation for Parkinson’s Research. Previous publications have described the LRRK2 Cohort Consortium in detail(81). CSF was collected as part of the LRRK2 cross-sectional study according to guidelines provided in the biologics manual which can be found at http://mjff.prod.acquia-sites.com/sites/default/files/media/document/LRRK2%20Cohort%20Consortium%20Biologics%20Manual%20Final%201.1.pdf.

### Data analysis for LC-MS based measurements

Peak areas for analytes detected were first normalized with spiked in internal standard areas. Pairing of specific analytes to surrogate internal standards are used are listed in supplementary tables 1-3.

### Statistical analysis of lipid levels from tissues and fluids of LRRK2 KO/G2019S mice

Lipid levels were log-transformed and analyzed using analysis of covariance (ANCOVA) models with terms for genotype (and terms for treatment group, or genotype x treatment interaction if applicable), and adjustment for key covariates (sex, age and/or weight). Specific covariates adjustments were study dependent. Effect sizes between groups were estimated via geometric mean ratios, and corresponding 95% confidence intervals (CI), and statistical significance was assessed at two-sided 5% significance levels. For each sample type, adjusted p-values corrected for multiple comparisons were derived using the Benjamini-Hochberg (BH) method.

### Analysis of human CSF from the LCC cohort

Raw area ratios of CSF lipids detected were normalized to equalize the median absolute deviation to 1 and log2 transformed using the limma package (version 3.46). Analysis of % differences in lipid abundance by LRRK2 and disease status was performed by robust linear model procedure using Age and Sex and covariates and the following three-way interaction model: log2(area ratio) ∼ LRRK2*PD status*Sex + Sex*Age. Post-hoc pairwise comparisons were analyzed using the estimated marginal means function provided in the emmeans r package (version 1.7.5). Conversion to % changes were performed using the update function provided in the stats r package (version 4.0.2). Correlation analysis between BMP (22:6/22:6) and GlcCer (d18:1/24:1) species stratified by LRRK2 and PD disease status was performed using the stat_cor function provided by ggpubr package (version 0.4.0). Visualization of heatmaps and boxplots were performed using the ggplot2 (version 3.3.6) and the ComplexHeatmap (version 2.6.2) packages. Data analyses above were performed using the R statistical software (version 4.0.2; R Core Team 2020).

## ACKNOWLEDGEMENTS

Data used in the preparation of this article were obtained from the Accelerating Medicine Partnership® (AMP®) Parkinson’s Disease (AMP PD) Knowledge Platform. For up-to-date information on the study, visit https://www.amp-pd.org. The AMP® PD program is a public-private partnership managed by the Foundation for the National Institutes of Health and funded by the National Institute of Neurological Disorders and Stroke (NINDS) in partnership with the Aligning Science Across Parkinson’s (ASAP) initiative; Celgene Corporation, a subsidiary of Bristol-Myers Squibb Company; GlaxoSmithKline plc (GSK); The Michael J. Fox Foundation for Parkinson’s Research; Pfizer Inc.; Sanofi US Services Inc.; and Verily Life Sciences.

ACCELERATING MEDICINES PARTNERSHIP and AMP are registered service marks of the U.S. Department of Health and Human Services. PPMI is sponsored by The Michael J. Fox Foundation for Parkinson’s Research and supported by a consortium of scientific partners. The PPMI investigators have not participated in reviewing the data analysis or content of the manuscript. For up-to-date information on the study, visit www.ppmi-info.org. The investigators within the LCC contributed to the design and implementation of the LCC and/or provided data and/or collected biospecimens, but did not necessarily participate in the analysis or writing of this report. The full list of LCC investigators can be found at www.michaeljfox.org/lccinvestigators. We thank Michael Schwarzschild and team for their contributions to the LCC CSF metabolomics data set. We thank Jamal Alkabsh for support for LC-MS based analyses and Todd Logan and other members of the lysosomal function pathway team at Denali for useful discussions and feedback.

