## Supplementary Information for "LRRK2 Kinase Activity Regulates Parkinson’s Disease-Relevant Lipids at the Lysosome"

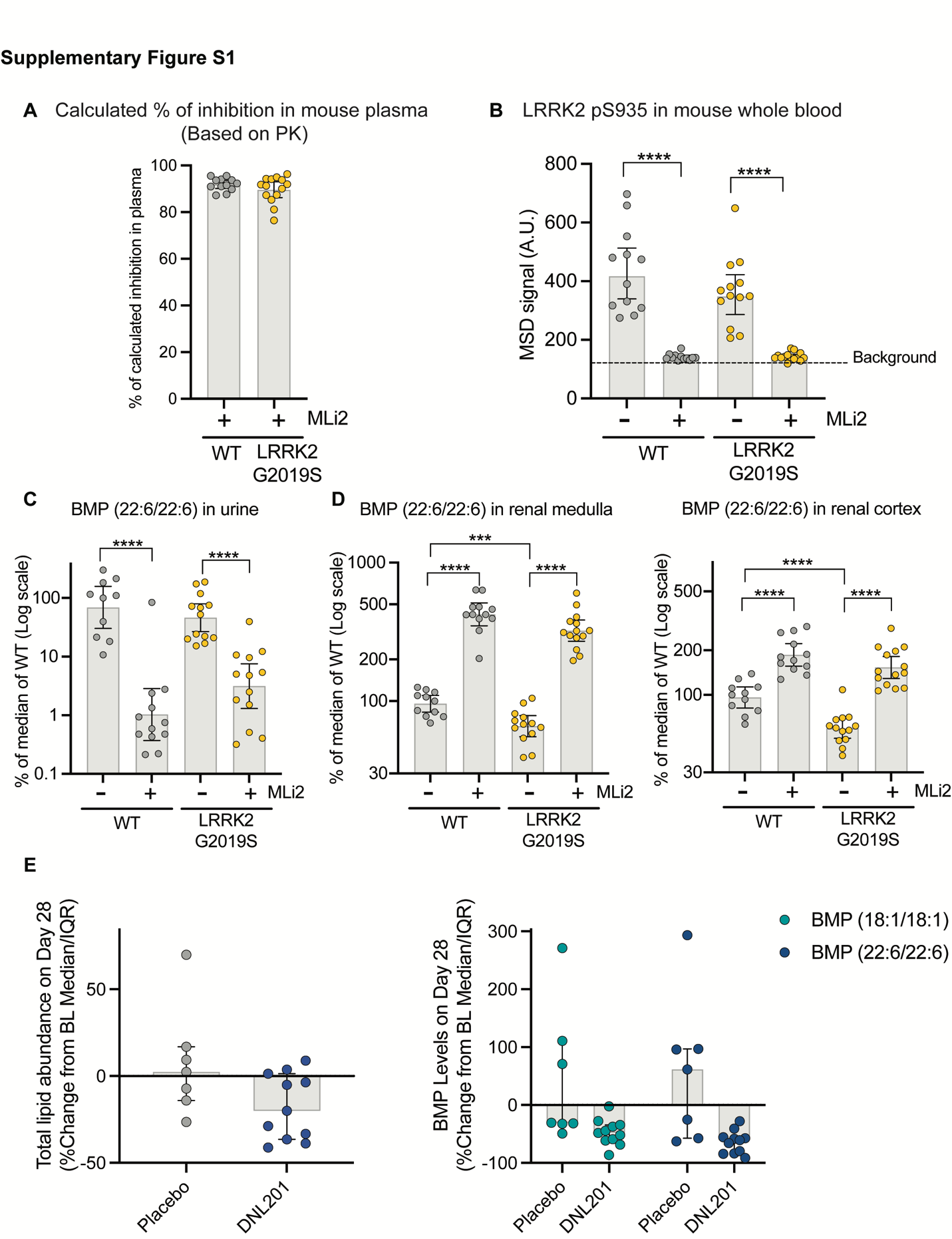

**Supplementary Figure S1: LRRK2 activity regulates BMP secretion in urine in mouse and in human subjects**. (A-B) Pharmacokinetics and pharmacodynamic analysis demonstrated >90% of LRRK2 kinase inhibition in peripheral in WT and LRRK2 G2019S KI mice dosed with MLi-2. (A) Calculated % of inhibition based on the unbound concentration of the drug in plasma demonstrated >90% inhibition. (B) The levels LRRK2 pS935 measured by MSD assay in whole blood showed significant reduction in WT and LRRK2 G2019S KI mice. Representative plots of BMP (22:6/22:6) in urine (C), renal medulla and renal cortex (D) from LRRK2 G2019S KI mice and WT littermates treated with or without MLi-2. Data are presented as % of the median values of WT-vehicle group and shown as geometric mean with 95% CI with p-values based on an ANCOVA model and statistical significance assessed at nominal levels. ***p < 0.001, ****p < 0.0001. E) Reduction of total lipid and BMP levels in urine exosomes collected from human subjects treated with DNL201 (n=11) compared with the placebo group (n=7). Lipid levels are expressed as percent change from pre- to post- dose. Data are shown as median with interquartile range.

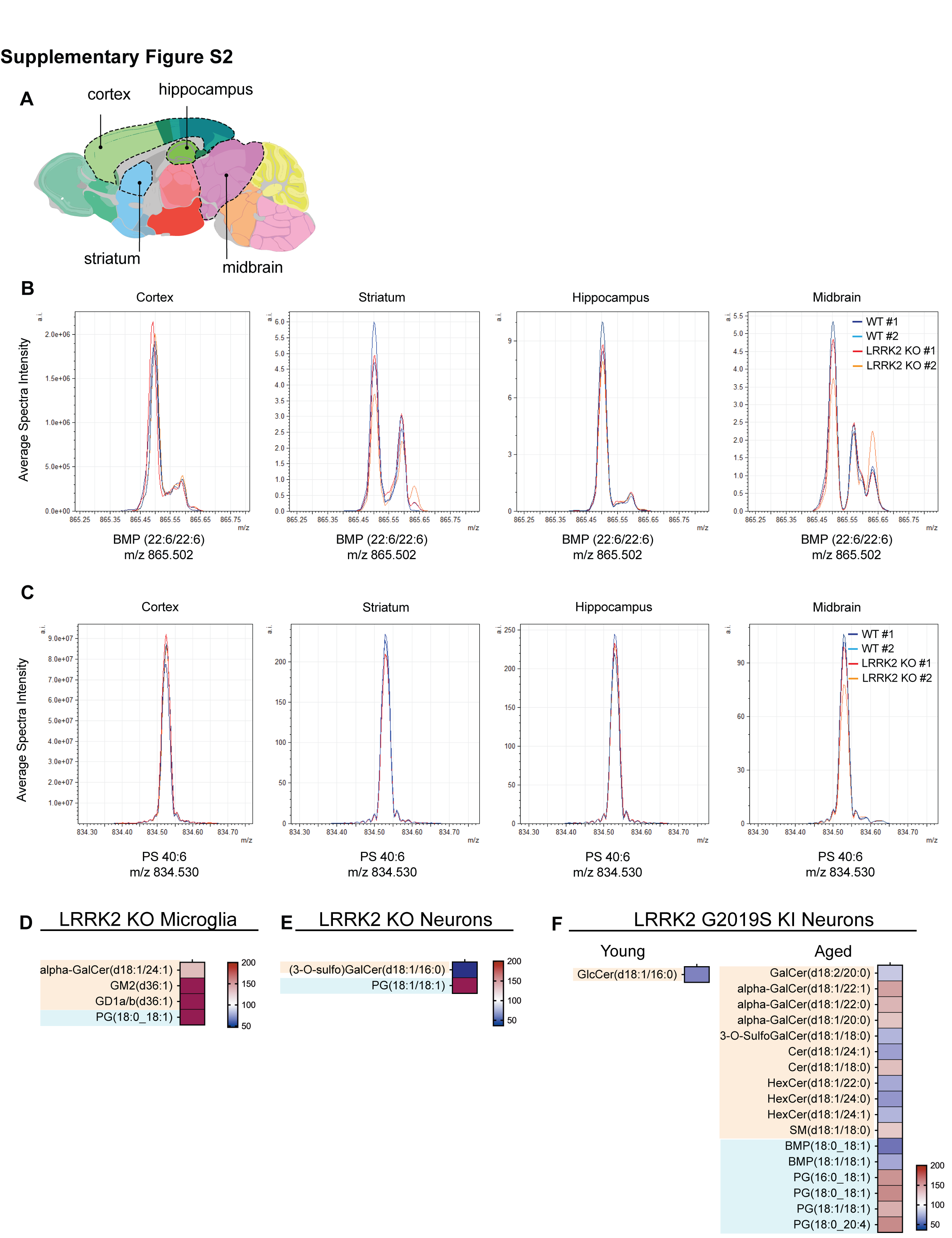
**Supplementary Figure S2: LRRK2 activity regulates glycosphingolipids in mouse brain and modestly impacts in BMP in cell-type specific manner.** (A) Schematic indicating approximate location of sagittal brain regions of interest (ROIs) for comparison of average spectra intensity in Fig. S2B and C. Spectra were averaged across each brain ROI from the mass spectrometry imaging experiments. Exported spectra at a mass/charge ratio (m/z) of (B) 865.502, corresponding to BMP (22:6/22:6) and (C) 834.530, corresponding to PS (40:6) are shown for selected ROIs. Data presented includes n = 2 mice per genotype with each color representing a biological replicate. Spectra show no significant differences in the intensities of BMP (22:6/22:6) or PS (40:6) in any of the selected brain regions. (D-F) Heatmaps demonstrated the changes in glycosphingolipids and BMP-related lipids in microglia from LRRK2 KO mice (D) and in neurons from LRRK2 KO mice (E) and LRRK2 G2019S KI mice at 5-6 month-old and 18-month old (F). The heatmaps were generated as % of change normalized to WT littermate controls. The analytes included had nominal p-values <0.10 for the genotype difference and then grouped based on lipid class. The BMP related lipids were shaded in cyan, and the glycosphingolipid species were shaded in orange. In the color scale, white depicts the WT amounts, as 100%, red shows an accumulation and blue shows a reduction.

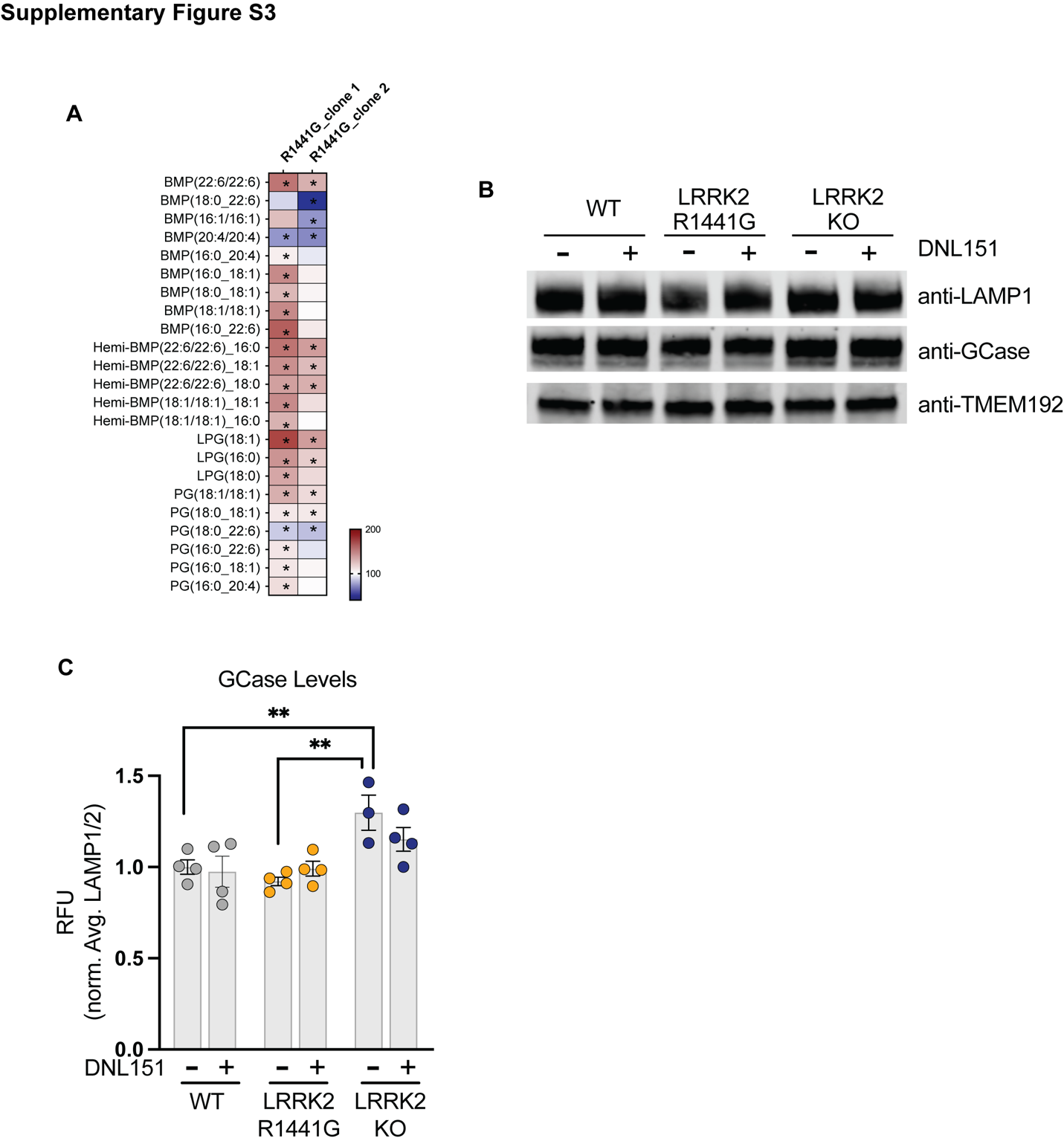
**Supplementary Figure S3: LRRK2 activity regulated glycosphingolipid catabolism in A549 cells but does not impact GCase levels in lysosomes.** A) Heatmap showing alternations in BMP-related lipid species in whole cell extracts from LRRK2 R1441G KI cells as compared to parental wild-type cells. The heatmaps were generated as % of change by normalizing the average of different groups to the average of the WT group. The analytes included had nominal p-values <0.10 for genotype differences in either clone and were grouped based on lipid class; *p<0.10. White in the color scale depicts the WT-vehicle amounts, as 100%, red shows an accumulation (capped at 350%), and blue shows a reduction B) Representative western blot analysis of lysosomes isolated from wild type, LRRK2 R1441G and GBA1 KO cells probed with anti-HA (to detect TMEM192-3x-HA), anti-LAMP1, anti-LAMP2 and anti-GCase antibodies. C) The signal from western blot analysis of lysosomes isolated from LRRK2 R1441G KI cells probed with an anti-GCase antibody was normalized to the average signals quantified using antibodies against LAMP1 and LAMP2 to normalize for lysosomal input; Data are shown as geometric mean ± SEM; n=3-4 independent experiments; **p < 0.01.

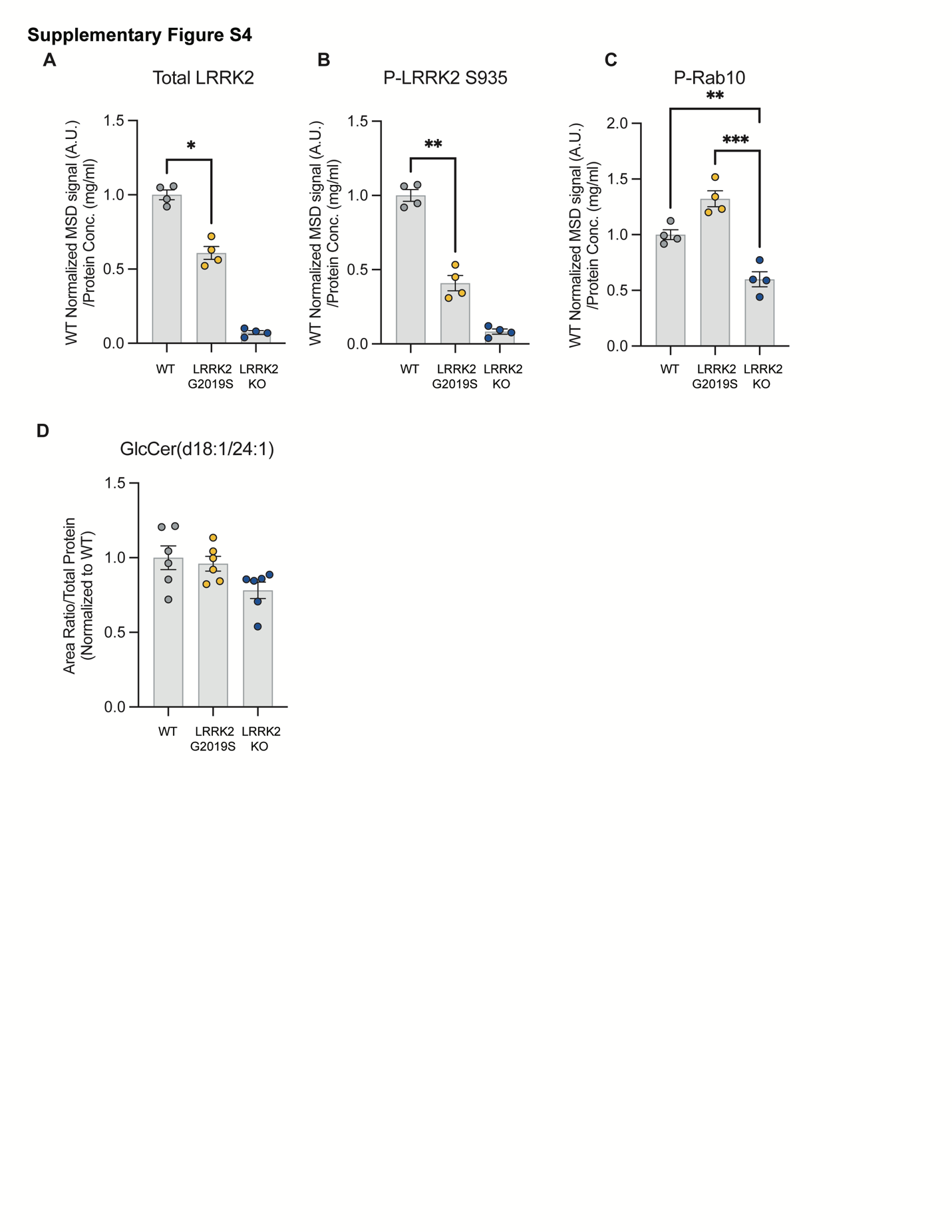
**Supplementary Figure S4: Analysis of LRRK2 levels, Rab10 phosphorylation, and GlcCer levels in LRRK2 G2019S and KO iMicroglia.** (A) Total-LRRK2, (B) pS935 LRRK2 and (C) pT73 Rab10 protein levels were measured in WT, LRRK2 G2019S and LRRK2 KO iMG cell lysates using MSD-based assays; n = 4 independent experiments; one-way ANOVA, Tukey’s method for multiple comparisons. (D) LC-MS-based analysis of GlcCer species shows no significant changes among the different LRRK2 genotypes in iMicroglia. *p < 0.05, **p < 0.01, ***p < 0.001. Data are shown as geometric mean ± SEM.

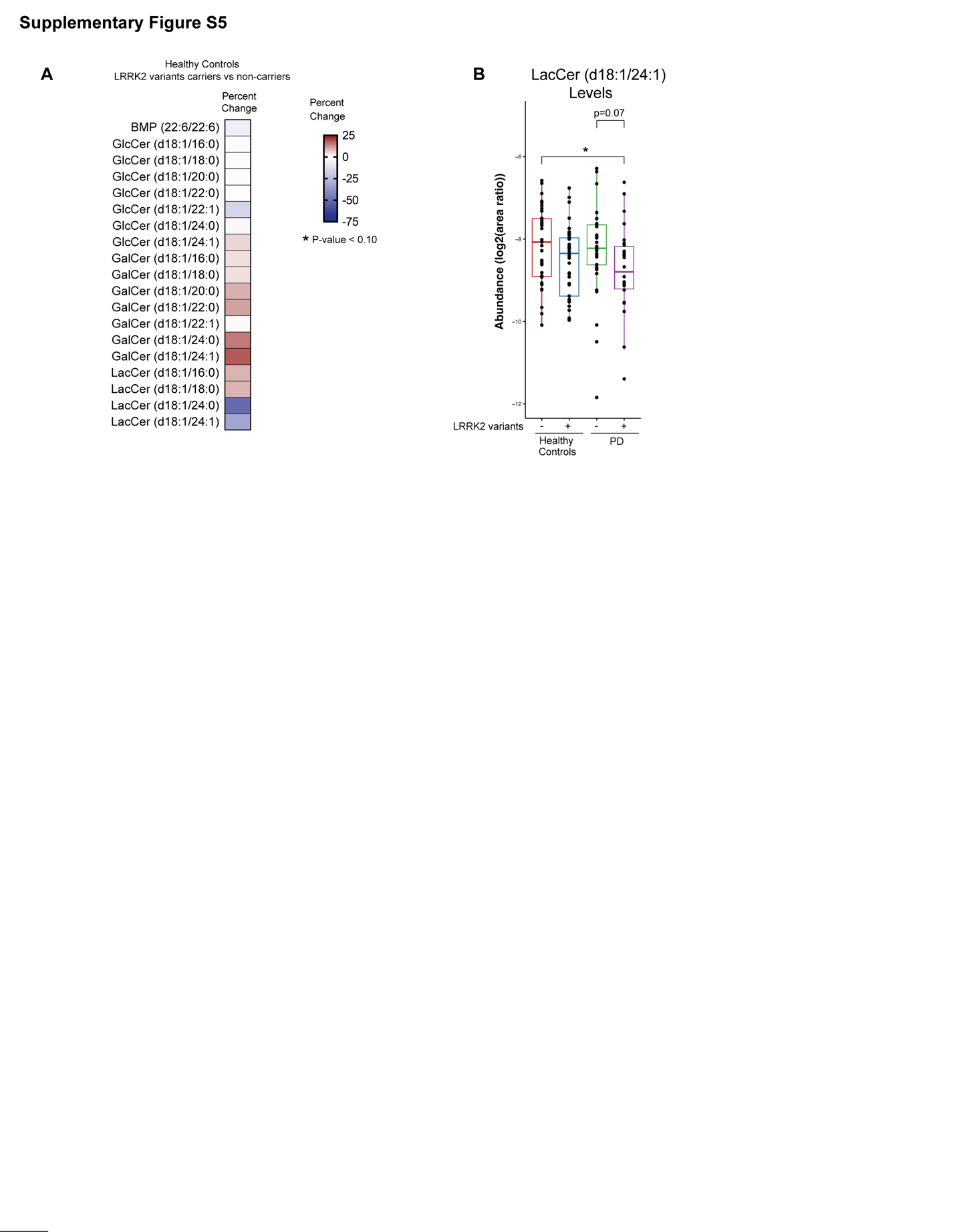

**Supplementary Figure S5: Analysis of BMP and GSL levels in human CSF from healthy subjects and PD patients with or without LRRK2 variants.** A) Heatmap showing % change in lipid abundance of BMP and GSL detected in human CSF from healthy subjects carrying LRRK2 variants compared to non-carriers. % changes and significance of effects were analyzed using robust linear model with sex and age as covariates. None of the analytes in the heatmap showed significant difference between the two groups, using unadjusted p <0.10 as cutoffs. B) Relative abundance of LacCer (d18:1/24:1) levels in CSF. Significance of change was analyzed by linear model with pairwise comparisons by Tukey’s honest significant difference test with significance set at unadjusted p value of 0.05. Main box and error bars depict interquartile ranges of top 75^th^ or bottom 25^th^ percentile and largest and smallest value with 1.5 times the interquartile ranges above and below 75^th^ or 25^th^ percentiles. Median 50^th^ percentile is shown as midline within each boxplot. *p < 0.05.

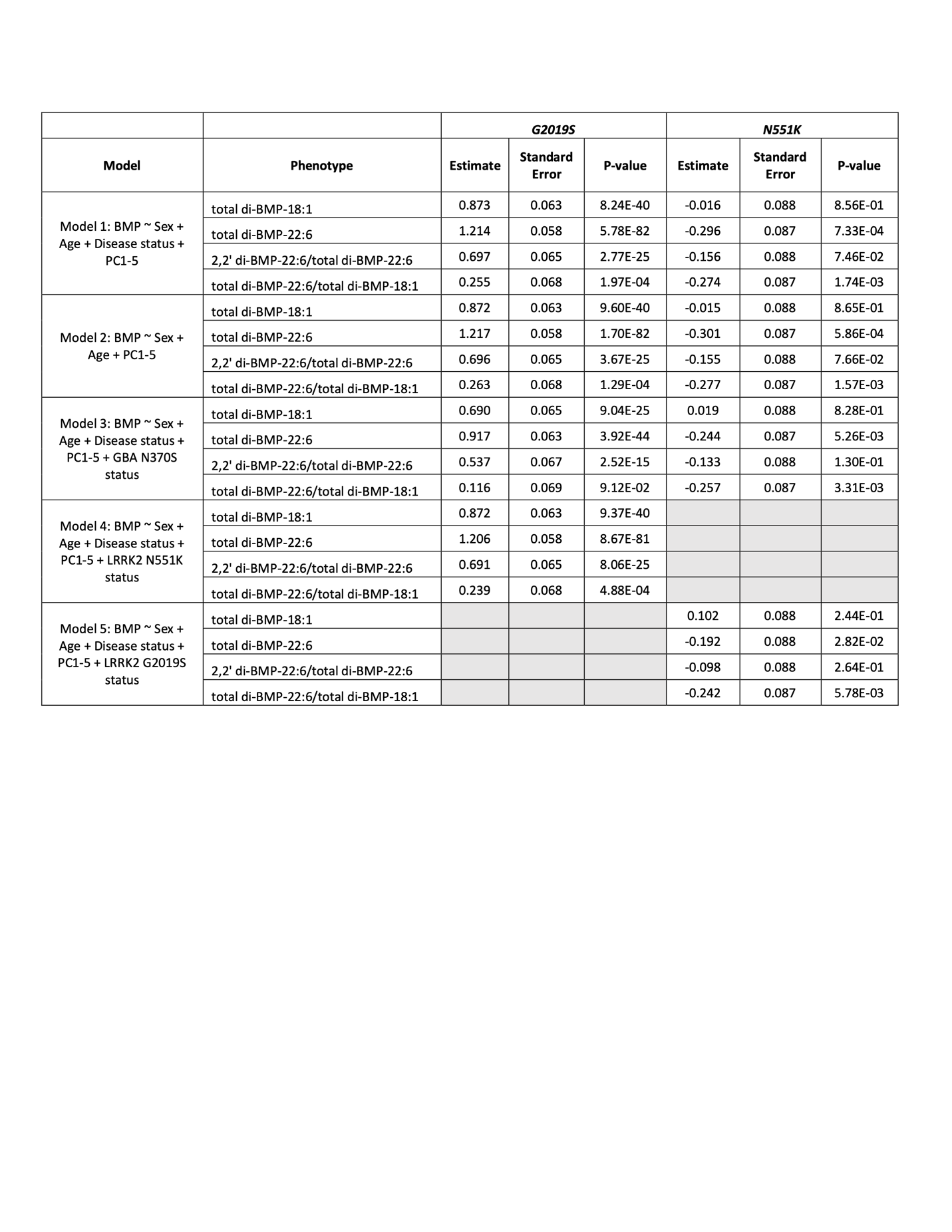
**Supplementary Table S1: Linear regression statistics for association testing of G2019S and N551K status on urine BMP levels in PPMI.** Urine BMP measurements were normalized to creatinine, log transformed, and fit in a linear model against the covariates listed under each “Model”. Residuals from each model where then inverse normal transformed and tested for association against G2019S status and N551K status. Estimate = standard deviation change in adjusted and transformed BMP levels for dosage of each G2019S/N551K allele.

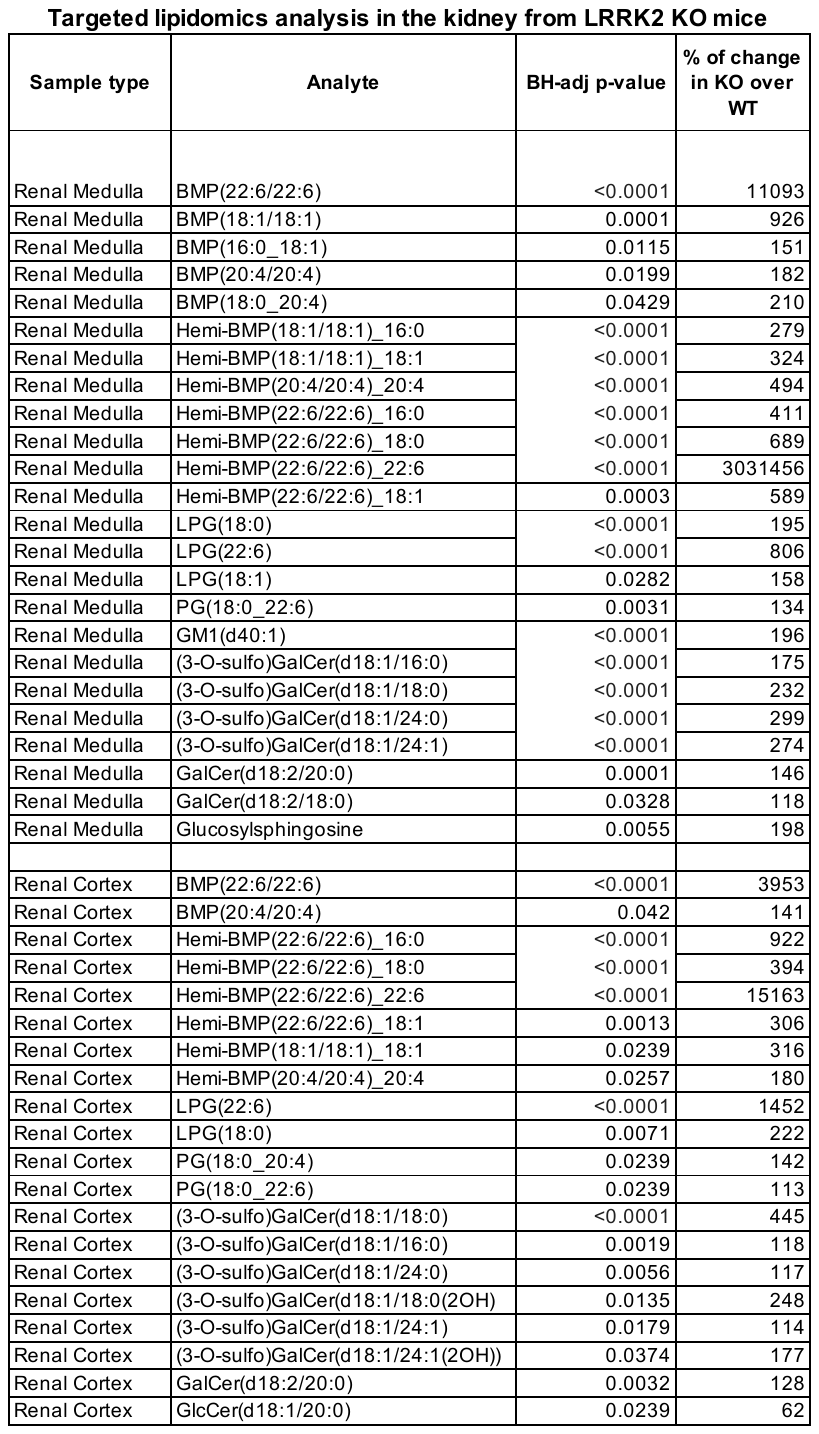

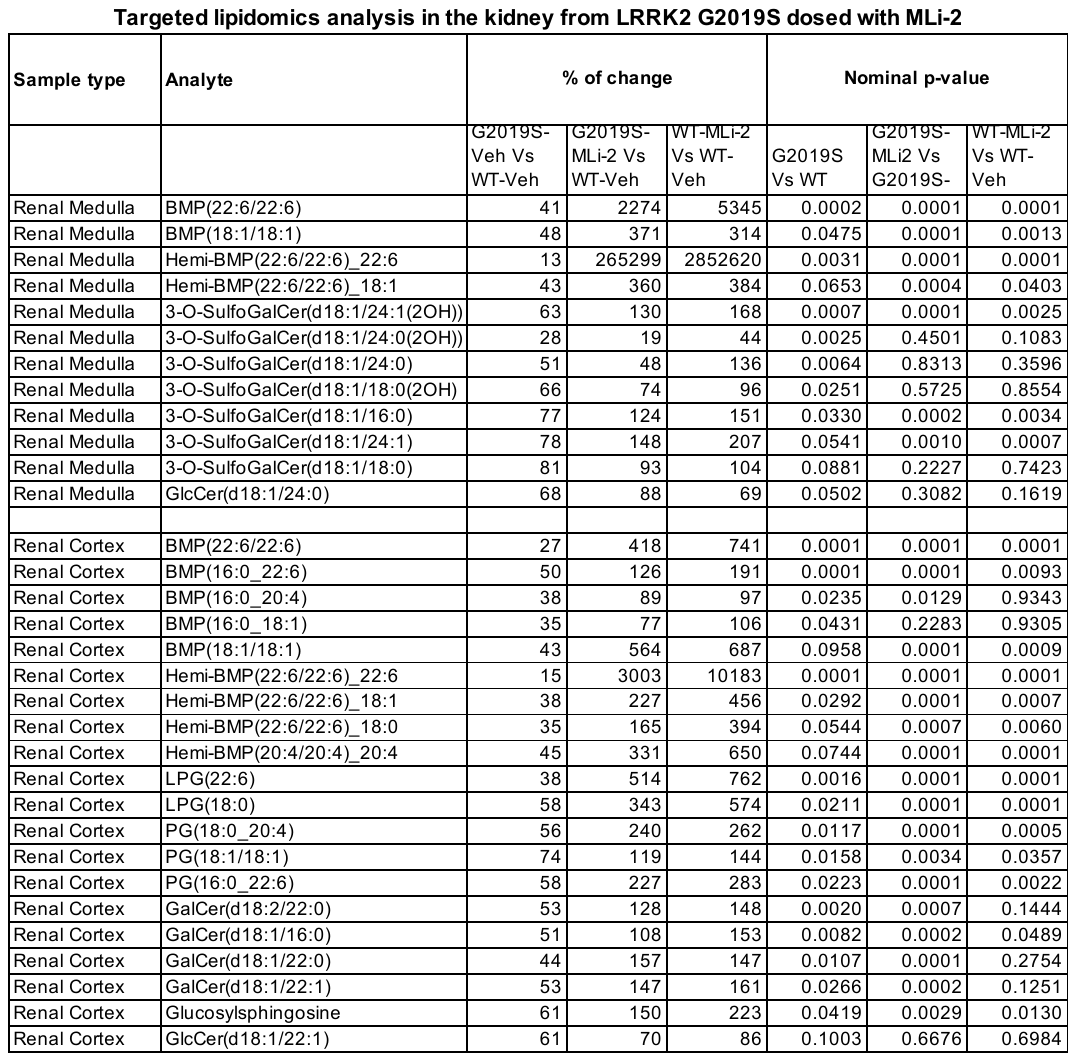

Note: p-values were rounded to 4 decimal points

**Supplementary Table S2: Targeted lipidomic analysis of BMP related lipids and glycosphingolipids in kidney from LRRK2 KO and G2019S KI mouse.** Lipids from a targeted panel were measured in by LC-MS/MS from renal cortex and renal medulla. The table included BMP related lipids and glycosphingolipids with BH-adjusted ANCOVA p-values <0.05 for the difference between KO Vs WT for table A, or nominal ANCOVA p-values <0.10 for the difference between G2019S KI Vs WT in table B.

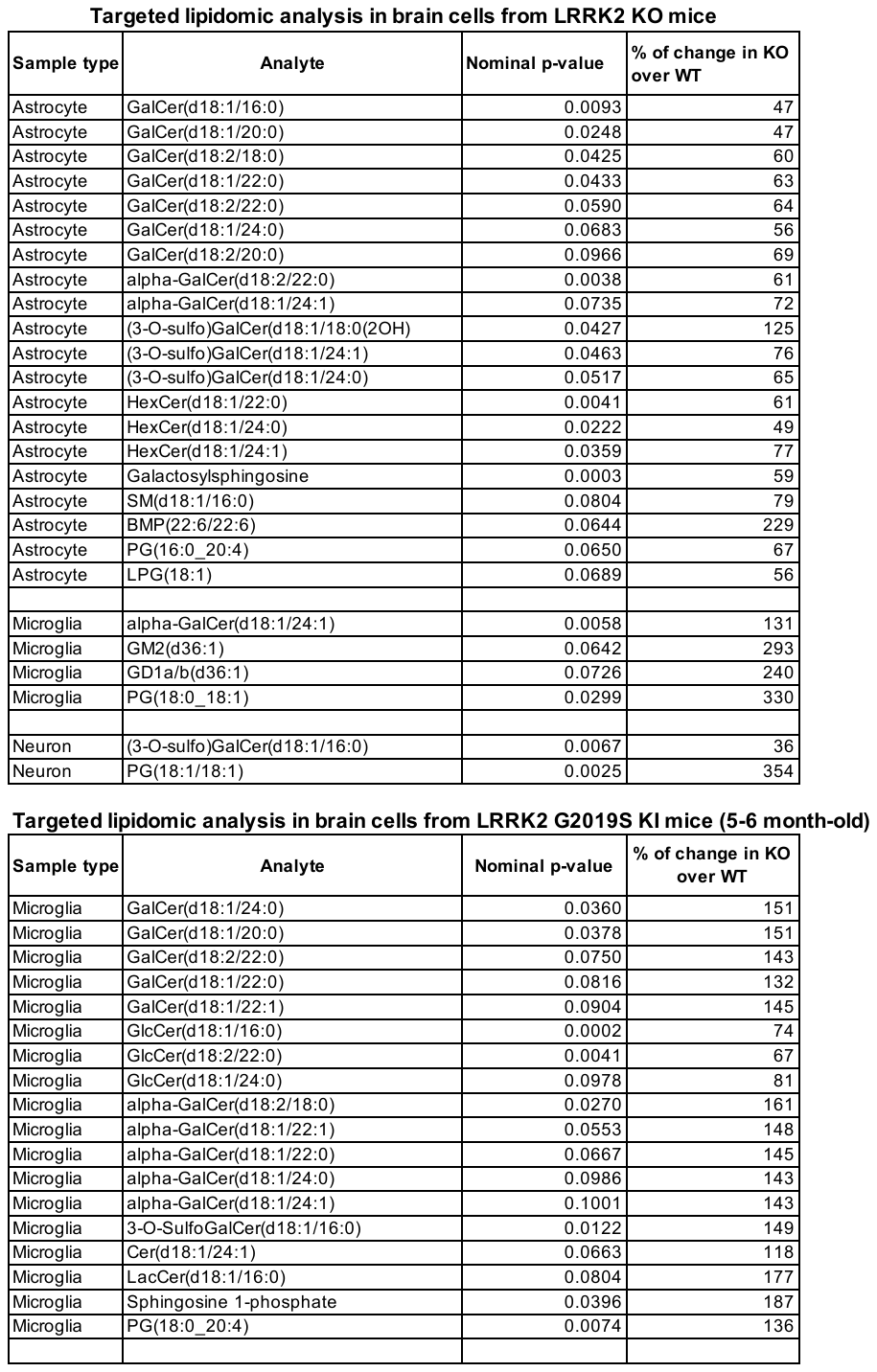

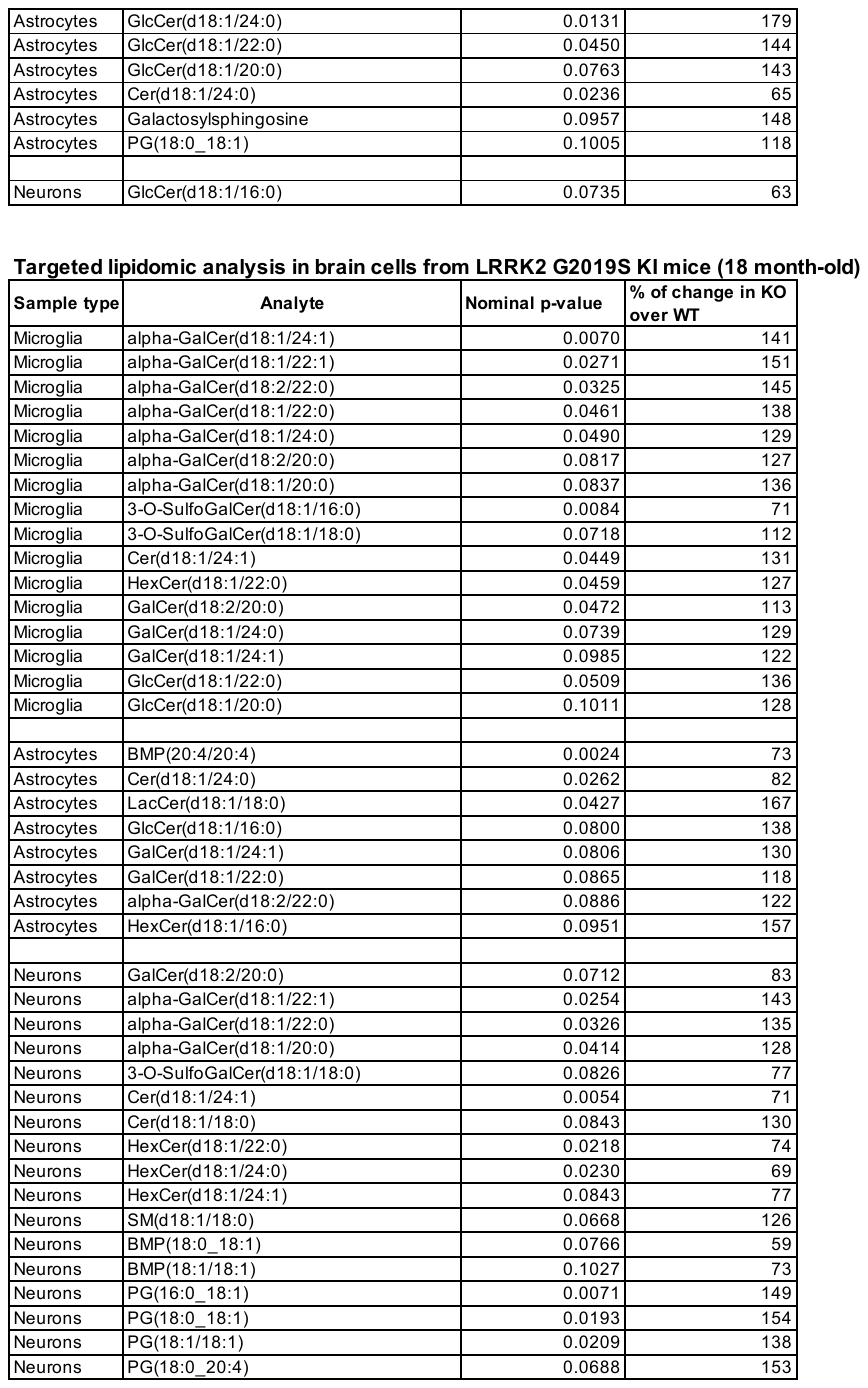

Note: p-values were rounded to 4 decimal points

**Supplementary Table S3: Targeted lipidomic analysis in CNS cells from LRRK2 KO and G2019S KI mice.** Lipids from a targeted panel were measured in by LC-MS/MS from FACS isolated astrocytes, microglia and neurons from LRRK2 mice. The table included BMP related lipids and glycosphingolipids with nominal p-values <0.10 for the genotype difference.

|  | LRRK2-  HC | LRRK2-  PD | LRRK2+  HC | LRRK2+  PD | Overall |
| --- | --- | --- | --- | --- | --- |
|  | (N=35) | (N=37) | (N=37) | (N=26) | (N=135) |
| Age |  |  |  |  |  |
| Mean (SD) | 53.0 (14.7) | 58.3 (11.5) | 50.0 (14.4) | 62.8 (11.5) | 55.5 (13.9) |
| Median  (Q1, Q3) | 54.0  (42.0, 64.0) | 59.0  (50.0, 67.0) | 50.0  (40.0, 61.0) | 65.0  (55.3, 70.3) | 57.0  (44.0, 66.0) |
| Min, Max | 24.0, 83.0 | 26.0, 77.0 | 27.0, 80.0 | 39.0, 80.0 | 24.0, 83.0 |
| Sex |  |  |  |  |  |
| Female | 19 (54.3%) | 13 (35.1%) | 17 (45.9%) | 13 (50.0%) | 62 (45.9%) |
| Male | 16 (45.7%) | 24 (64.9%) | 20 (54.1%) | 13 (50.0%) | 73 (54.1%) |

**Supplementary Table S4: Demographics of LRRK2 Cohort Consortium Participants with CSF Analyzed in this Study.** The demographics of the metabolomic and lipidomic CSF profiling study from the LRRK2 Cohort Consortium are summarized. HC: Healthy Control; PD: Parkinson’s disease; LRRK2-: Participants not carrying a LRRK2 pathogenic point mutation; LRRK2+: Participants carrying a LRRK2 pathogenic point mutation.
